# Chemical modification of hyaluronan oligosaccharides differentially modulates hyaluronan- hyaladherin interactions

**DOI:** 10.1101/2024.03.12.584658

**Authors:** Rebecca J. Dodd, Charles D. Blundell, Benedict M. Sattelle, Jan J. Enghild, Caroline M. Milner, Anthony J. Day

## Abstract

The glycosaminoglycan hyaluronan (HA) is a ubiquitous, non-sulphated polysaccharide with diverse biological roles mediated through its interactions with HA-binding proteins (HABPs). Most HABPs belong to the Link module superfamily, including the major HA receptor, CD44, and secreted protein TSG-6, which catalyzes the covalent transfer of Heavy Chains (HC) from inter-a-inhibitor (IaI) onto HA. The structures of the HA-binding domains (HABD) of CD44 (HABD_CD44) and TSG-6 (Link_TSG6) have been determined and their interactions with HA extensively characterized. The mechanisms of binding are different, with Link_TSG6 interacting with HA primarily via ionic and CH−π interactions, whereas HABD_CD44 binds solely via hydrogen bonds and van der Waals forces. Here we exploit these differences to generate HA oligosaccharides, chemically modified at their reducing ends, that bind specifically and differentially to these target HABPs. Hexasaccharides (HA_6_^AN^) modified with 2- or 3-aminobenzoic acid or 2-amino-4-methoxybenzoic acid (HA_6_-2AA, HA_6_-3AA, HA_6_-2A4MBA, respectively) had increased affinities for Link_TSG6 compared to unmodified HA_6_^AN^. These modifications did not increase the affinity for CD44_HABD. A model of HA_6_-2AA (derived from the solution dynamic 3D structure of HA_4_-2AA) was docked into the Link_TSG6 structure, providing evidence that the 2AA-carboxyl forms a salt bridge with Arginine-81. These modeling results informed a 2^nd^ series of chemical modifications for HA oligosaccharides, which again showed differential binding to the two proteins. Several modifications to HA_4_ and HA_6_ were found to convert the oligosaccharide into substrates for HC-transfer, whereas unmodified HA_4_ and HA_6_ are not. This study has generated valuable research tools to further understand HA biology.

## Introduction

Hyaluronan (HA) is a ubiquitous, non-sulfated, glycosaminoglycan composed of repeating disaccharides of glucuronic acid (GlcA) and *N*-acetylglucosamine (GlcNAc). HA is a major structural component of the extracellular matrix in all vertebrates, where it typically has a molecular weight between 10^5^ and 10^7^ Da, also being an essential constituent of the glycocalyx (sugar coat) that surrounds most cells (1–3). Synthesis and breakdown of HA within tissues is tightly regulated and increases in hyaluronan biosynthesis are often associated with disease. For example, HA accumulation occurs rapidly in the lungs of mice following influenza infection (4, 5) and SARS-CoV2 can also increase pulmonary HA in human and mouse models (6, 7); in this context, HA is believed to be in part responsible for increased fluid retention and impaired gas exchange leading to lung dysfunction. Conversely, the improved health span and resistance to cancer in the naked mole rat have been attributed to high levels of HA that accumulate in the tissues of this long-lived rodent (8). There is evidence that some roles of HA are dependent on its molecular weight (9–11). For example, high molecular weight HA (in addition to phosphatidylcholine) is essential for sufficient lubrication at articular joint surfaces (12–14) whereas production of HA fragments (<500 kDa) has been reported to be associated with wound healing and cell signaling responses (15, 16), as well as indicating severe COVID-19 infection (17). However, it is likely that some of the pro-inflammatory effects attributed to low molecular weight HA, *e.g*., via signaling through Toll-like receptors (18) are a result of LPS contamination in the HA preparations used (19).

The diversity of HA biology largely stems from its interactions with a repertoire of HA binding proteins (HABPs), also known as ‘hyaldherins’, leading to the formation of HA/protein complexes with distinct architectures engendering particular biochemical and biophysical properties (1, 20, 21). Different HABPs have their own spatial and temporal expression patterns and are associated with tissue response to damage or inflammation (20). Many but not all HABPs contain Link modules (20, 22) – a protein domain consisting of ∼90 amino acid residues that mediate binding to HA (1, 23). The secreted glycoprotein TSG-6 and the HA receptor CD44 are two members of the Link module superfamily whose structure and function have been studied in detail. TSG-6 (which contains a single Link module) is upregulated in response to inflammation and has potent anti-inflammatory and tissue protective properties (24, 25). One of TSG-6’s functional activities is to act as an enzyme mediating the covalent modification of HA with heavy chains (HCs) derived from the plasma proteoglycan inter-a-inhibitor (IaI) (26, 27); a process involving the formation of HC•TSG-6 intermediates (26, 28). The compositions and functions of the resulting HC•HA complexes are tissue and context dependent (see (25)). For example, expansion of the cumulus matrix is driven by the production of HC•HA and plays a critical role in ovulation and fertilization (27, 29) whereas accumulation of HC•HA matrices during lung inflammation is believed to contribute to pathology, *e.g.,* in influenza infection or allergic airway responses (4, 30). CD44, which contains an extended Link module (31, 32), is the major cell surface HA receptor and contributes to many biological processes, including anchoring of stromal cells into the extracellular matrix, and mediating leukocyte trafficking (*e.g.,* during inflammation) and metastasis (33, 34). Binding between HA and CD44 is tightly regulated through alternative splicing, post-translational protein modification and dependency on how the HA is presented (32, 35–37). For example, HA chains cross-linked by TSG-6 show enhanced binding to CD44^+^ lymphocytes (37) and HC•HA from synovial fluids of rheumatoid arthritis patients is more adhesive to CD44^+^ cells than HA alone (38) although HC•HA made in *in vitro* is not (36).

In addition to different biological functions, it is apparent that the mechanisms by which TSG-6 and CD44 interact with HA are distinct (31, 39), *i.e*., based on biochemical and structural studies with Link_TSG6, the independently folding Link module of human TSG-6, and the HA-binding domain (HABD) of CD44 (HABD_CD44) (31, 32, 39–43). The Link modules of both CD44 and TSG-6 are comprised of two α-helices and two triple stranded anti-parallel β-sheets; CD44’s HABD includes flanking sequences providing 4 additional β-strands that extend the globular structure (23, 31, 32, 41, 44).

For Link_TSG6, key HA-binding residues (Lys11, Tyr12, His45, Tyr59, Lys63, Phe70, Tyr78 and Arg81) are found in and around a shallow groove on the surface of the Link module, and upon binding there is a ligand-induced conformational change in the protein (39–42, 44); this occurs mostly within a loop region connecting the b4 and b5 strands. The conformation of the bound HA was inferred from molecular modeling with an HA octasaccharide (HA_8_^AN^; with a non-reducing terminal GlcA (A) and reducing terminal GlcNAc (N)) being docked into the HA-bound conformation of Link_TSG6 (41) based on experimentally derived restraints (39, 42). This revealed that in the refined model (39), the binding of Link_TSG6 to HA_8_^AN^ (where 7 of the sugar rings make contact with the protein) is mediated by a combination of stacking interactions between histidine/tyrosine amino acids and the sugar rings along with ionic interactions between arginine/lysine residues and the carboxyl moieties of the GlcA sugars (**Figure 1**). This extensive interaction network induces a pronounced kink between GlcNAc2-GlcA3 and GlcA3-GlcNAc4 relative to the conformer preferred in free solution (albeit with f/y angles that are energetically favorable (39)). These insights were obtained from the analysis of binding of Link_TSG6 to different sizes of HA oligosaccharides (HA_4_-HA_8_) using isothermal titration calorimetry (ITC) and NMR spectroscopy. This work also demonstrated that HA_8_^AN^ is the minimum length required for maximal binding; oligosaccharides shorter that HA_8_^AN^ have reduced affinity because they cannot make the full complement of interactions (39). The binding between Link_TSG6 and HA is also pH-dependent and is maximal at pH 6.0. At higher pH values, the affinity is reduced because of a change in the protonation state of His4 that disrupts a salt bridge and hydrogen bond network connecting this residue to the HA-binding site (40, 43).

**Figure 1.**
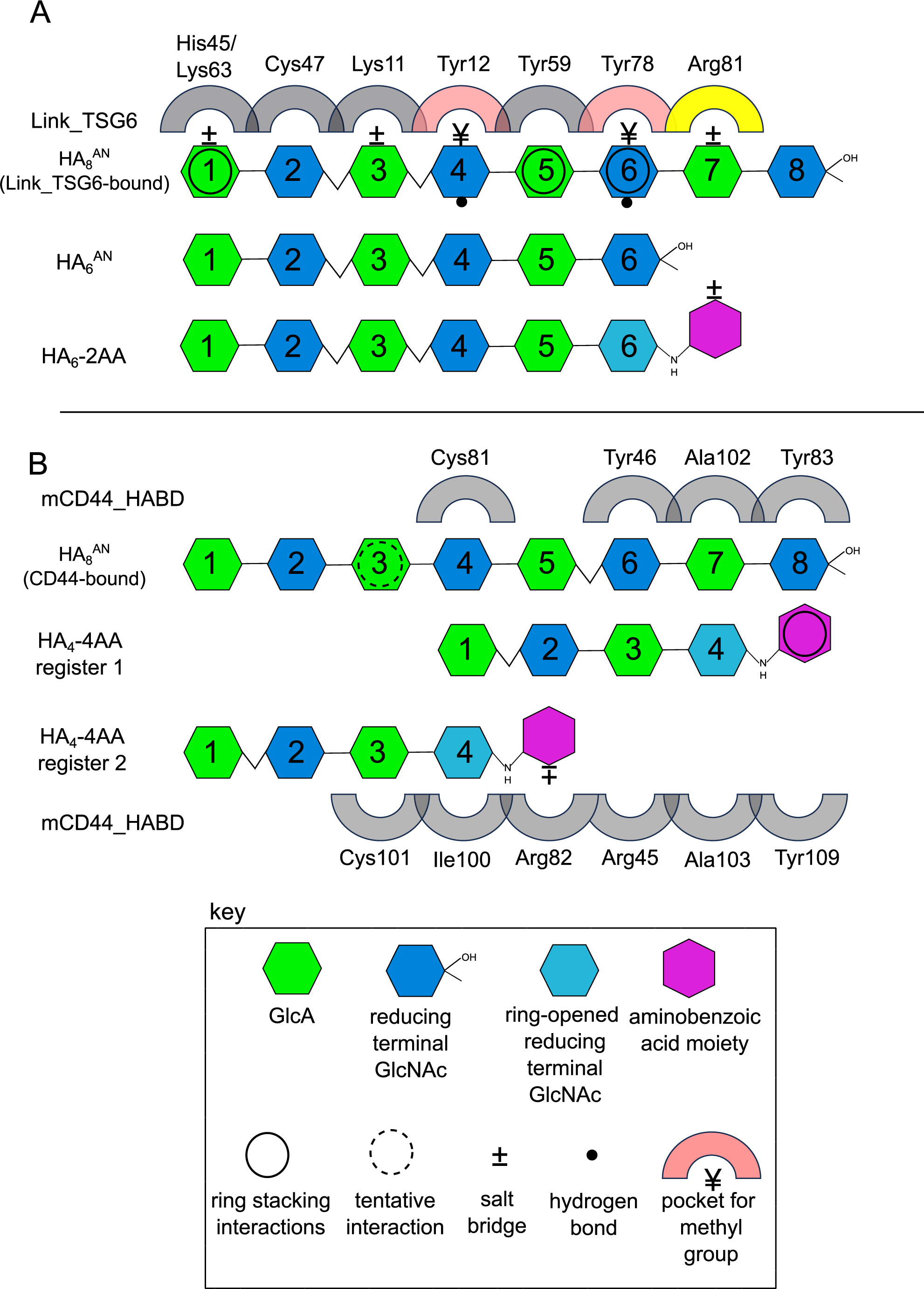
Model of interactions between HA oligosaccharides and CD44_HABD and Link_TSG6. Key HA binding residues of Link_TSG6 or mouse CD44_HABD (mCD44_HABD) are shown (grey arches) with the suggested binding registers of various unmodified (HA_6_^AN^, HA_8_^AN^) or modified HA oligosaccharides (HA_6_-2AA, HA_4_-4AA). **A**) Interactions of HA_6_^AN^ and HA_8_^AN^ with Link_TSG6 inferred from NMR, mutagenesis and ITC data between the oligosaccharide and protein are indicated as described in (39); black circles = CH−π interactions; ± = a salt bridge; • = a hydrogen bond; ¥ = methyl (Me) group in specificity pocket (light pink arch). Arg81 (the residue that was targeted for additional interactions) is colored in yellow. B) For CD44, some of the key binding residues are shown (for HA_8_^AN^) as described in (32) along with proposed binding registers for modified HA_4_-4AA.

The HA-binding site in CD44 is also well-defined, based on NMR spectroscopy and X-ray crystallography studies using both human and mouse proteins (31, 32). CD44 also has a shallow binding groove on the Link module surface in a similar position to that seen in TSG-6 (39). From the crystal structure of murine CD44 in complex with HA_8_^AN^ (31), it is apparent that the surface involved in the binding of HA to CD44 is smaller than that observed for Link_TSG6 (39), with only five sugar rings making contacts with the protein. However, from NMR experiments (with different sizes of HA oligomers), an octasaccharide (HA_8_^AN^) has been suggested to be the minimal length of HA that completely occupies the HA-binding site (32). Unlike Link_TSG6, the interaction between CD44 and HA does not involve either ionic or CH−π interactions (**Figure 1**), but rather is mediated by hydrogen bonds and van der Waals contacts (31). This difference in the molecular basis of HA binding to the two proteins likely explains the higher affinity seen for the interaction of HA_8_^AN^ with Link_TSG6 compared to CD44_HABD (31, 39), and is also consistent with their distinct physiological roles. In addition, CD44 and TSG-6 capture different HA conformations: binding of HA_8_^AN^ to Link_TSG6 results in a kink in the chain between GlcNAc2 and GlcA3 and between GlcA3 and GlcNAc4 (39), whereas binding of the same length oligosaccharide to CD44_HABD results in a kink between rings GlcNAc5 and GlcA6 (31).

From the structural and biochemical studies described above, it is clear that CD44 and TSG-6 interact with HA via distinct mechanisms, and it seems likely that this will be true of other HABPs, *i.e.,* as suggested from sequence analysis of the Link module superfamily (40, 42). Therefore, it should be feasible to exploit these differences in order to specifically target individual HA-binding proteins, *i.e.,* with CD44_HABD and Link_TSG6 providing an excellent model system to test this possibility. Hyaluronan oligosaccharides of different lengths can be readily purified (45, 46) and have been used in previous structural and functional studies on CD44 and TSG-6 (31, 32, 39, 42). Therefore, they represent a good starting point, especially since short HA oligomers provide ready-made scaffolds for chemical modification (47). Lu and Huang (48) used HA tetrasaccharides as scaffolds to generate compounds that could potentially inhibit binding of HA to CD44 in competition assays. In addition to single modifications to oligosaccharides, chemical derivatization of polymeric HA (*e.g.,* 10 kDa and above) has also been described as a strategy to generate inhibitors or probes for HABPs. Liu *et al.* (48) recently reported the generation of a library of carboxyl-modified HA polysaccharides containing two derivatives that could inhibit binding of native HA to CD44; the inhibitory potential of these derivatives increased with the size of the HA modified, likely due to the multivalency and superselectivity of CD44-HA binding (10, 11, 48–50). However, some modifications to polymeric HA reduce binding affinity, for example sulfation of the 6O-groups of GlcNAc prevents binding of the modified HA to CD44 (51, 52).

Here, we describe the design, synthesis, and activity of two series of chemically modified HA oligosaccharides and demonstrate that specific modifications can differentially alter the affinities of these oligosaccharides for either Link_TSG6 or CD44_HABD. Furthermore, we show that certain modifications of HA oligosaccharides can alter their activity as substrates for HC-transfer, demonstrating the potential of modified HA oligosaccharides as tool compounds to further explore HA biology.

## Results

### Chemical modification of HA oligosaccharides with aminobenzoic acid

We have previously modified HA oligosaccharides at their reducing terminus with 2-aminobenzoic acid (2AA) to generate fluorescent probes (53). This modification was selected here for our initial studies, in part because of the existing methodology, but also since 2AA contains a carboxyl group, introducing an additional negative charge to the HA oligosaccharide in a similar register to that of the next GlcA ring. We hypothesized that aminobenzoic modified oligosaccharides would show increased affinity for Link_TSG6 over CD44_HABD given that the HA-binding groove of Link_TSG6 contains basic residues (*e.g.,* Arg81) capable of making ionic interactions with this moiety (39, 41, 42). To investigate whether the position and orientation of the introduced carboxyl group were important, we also generated oligosaccharides modified with 3-aminobenzoic acid (3AA) and 4-aminobenzoic acid (4AA; **Table S1**). These modifications were carried out by reductive amination (**Figure 2A**) performed simultaneously on a pool of oligosaccharides (*e.g.,* HA_4_^AN^, HA_6_^AN^, and HA_8_^AN^_),_ resulting from the digestion of sodium hyaluronate with ovine testicular hyaluronidase (45); see details in Experimental Procedures.

**Figure 2.**
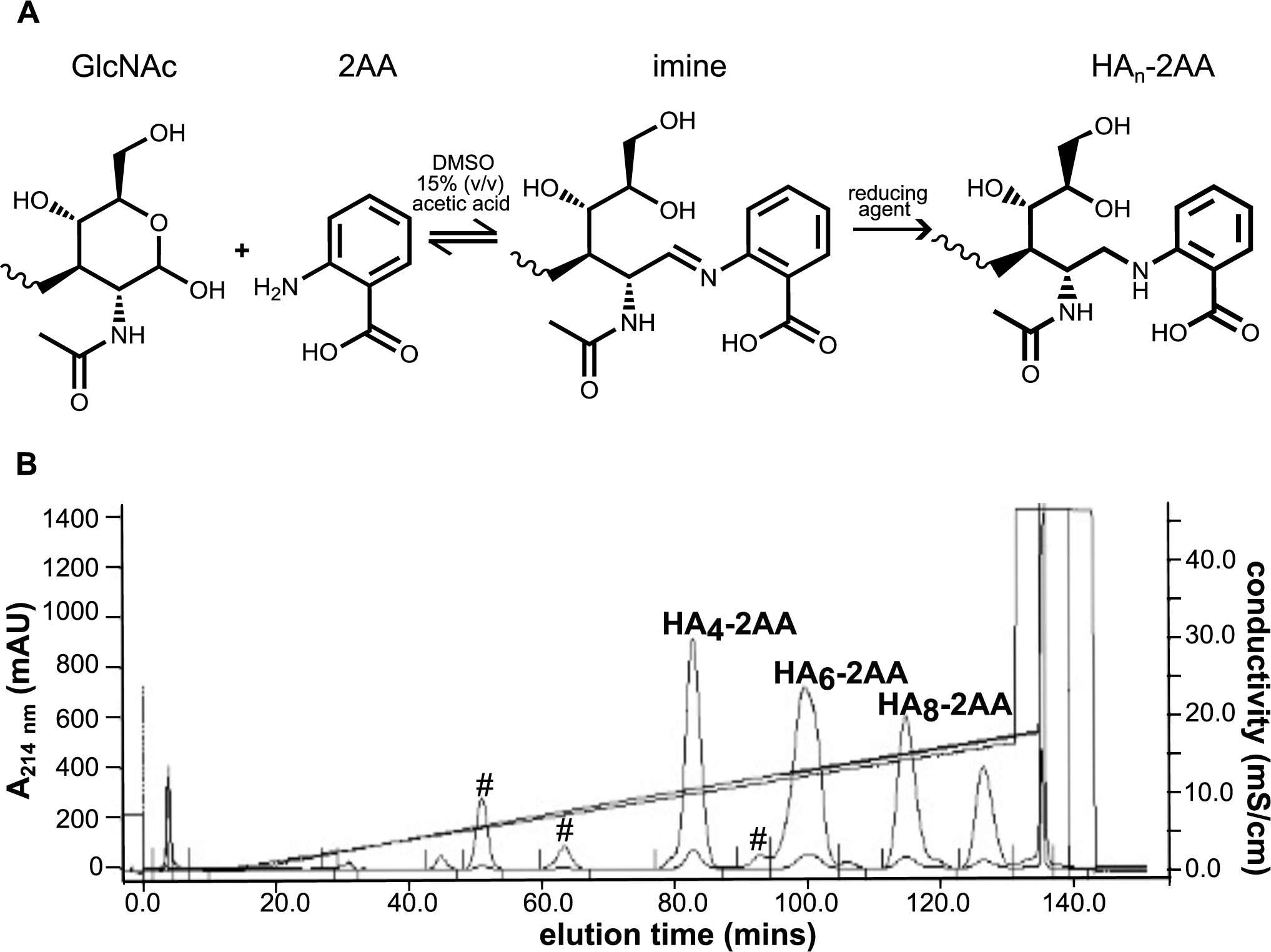
Modification of HA_n_^AN^ oligosaccharides. **A**) Reaction scheme for reductive amination of HA oligosaccharides with reducing terminal GlcNAc, illustrated for modification with 2-amino-benzoic acid (2AA). Reactions were incubated at 65 °C in DMSO with 15% (v/v) CH3COOH in the presence of 1 M 2-picolene borane as reducing agent. **B**) Representative anion exchange chromatogram showing purification of 2-AA modified oligosaccharides of different length. Elution of oligosaccharides was monitored at 214 nm and modified oligosaccharide peaks are labelled, as determined by MALDI-TOF MS and NMR (see **Table S2**). Peaks indicating remaining unmodified oligosaccharides are indicated with #.

The 2AA-, 3AA-, and 4AA-modified oligosaccharides were first purified from unreacted chemical reagents by size exclusion chromatography (SEC). The modified oligosaccharides of differing lengths (*e.g.,* HA_4_^AN^-HA_10_^AN^) were then separated from each other and from the unmodified oligomers (to baseline resolution) using anion exchange chromatography (AEX; **Figure 2B**); aminobenzoic acid modified HA oligosaccharides eluted at greater salt strengths compared to unmodified oligomers of the same length because of the introduction of the additional carboxylate group. The identity and purity of the oligosaccharides were determined by matrix-assisted-laser-desorption-ionization time-of-flight mass spectrometry (MALDI-TOF; **Table S2** and **Figure S1**) and 1D-^1^H NMR spectroscopy (**Figure 3** and **Figure 4**). The observed molecular masses of modified and unmodified oligosaccharides were found to be within 1.5 Da of their theoretical values (**Table S2**). From the NMR analyses, differences were observed (as expected) in the amide and aromatic proton region for the modified oligosaccharides with resonances arising from the aminobenzoic acid group (2AA, 3AA or 4AA; **Figure 4**) and the loss of resonances from the closed reducing terminal rings (*i.e.,* α-NH and β-NH groups; **Figure 3**). However, the remainder of the spectra were essentially unchanged compared to the unmodified oligosaccharides (**Figure 3**). For example, resonances from the non-reducing terminal GlcNAc amides were still observed and the volume of the internal GlcNAc amide peak increased with the length of oligosaccharide, as described previously (54).

**Figure 3.**
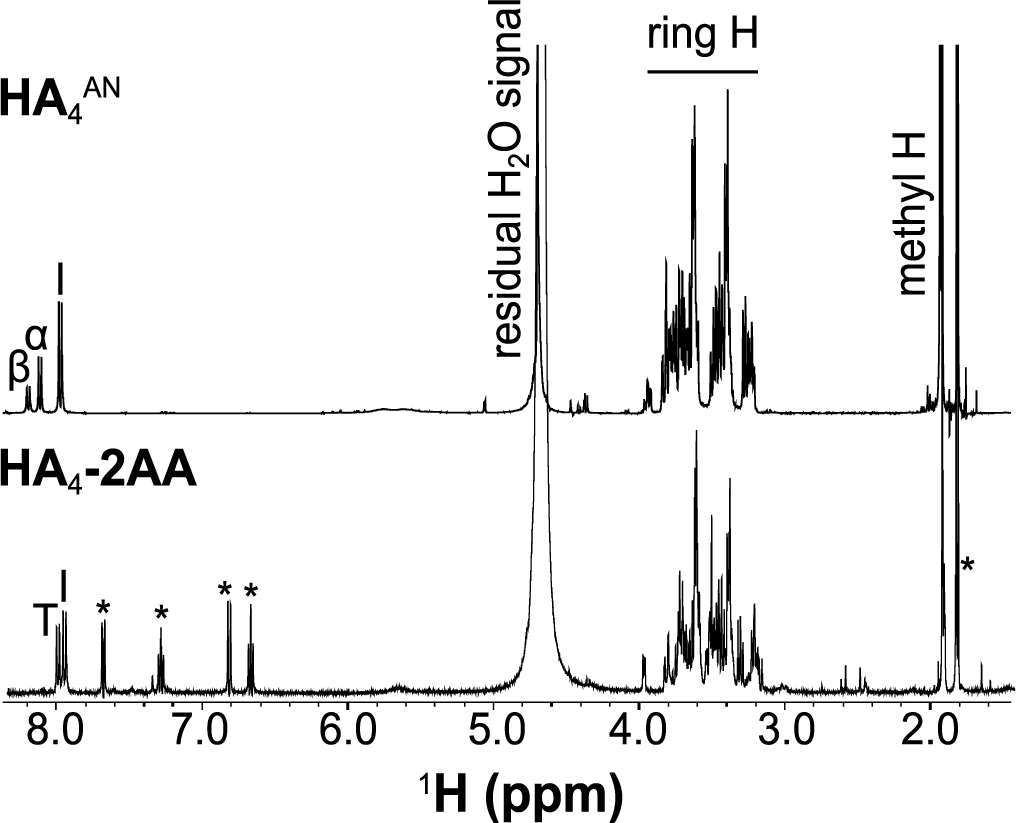
NMR analysis of HA_4_-2AA. Comparison of ^1^H-1D NMR spectra of HA_4_^AN^ (top) and HA_4_-2AA (bottom) (pH 6.0, 25 °C, 500 MHz). In HA_4_^AN^, resonances from the closed α- or β-anomers of the reducing terminal GlcNAc amide are visible (labeled a and b, respectively), along with the internal GlcNAc amide (labeled I). Modification with 2AA results in minimal perturbation of the oligosaccharide ring H region, whereas, in the amide region, the resonances from closed α and β are collapsed into a single new resonance (T) by the chemical modification and new resonances arising from the 2AA group are introduced (indicated with *).

**Figure 4.**
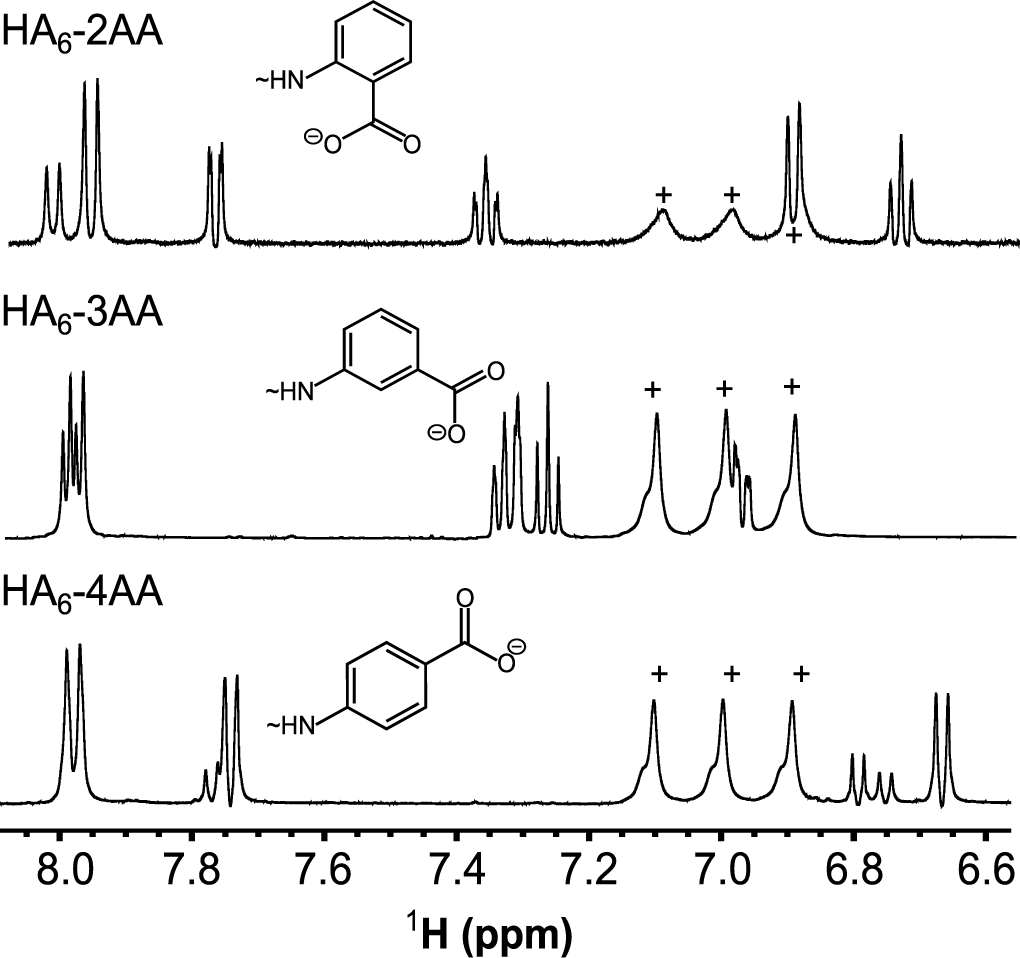
NMR analysis of aminobenzoic acid-modified HA hexasaccharides. Portions of ^1^H-1D NMR spectra recorded on HA_6_^AN^ modified with 2AA (top), 3AA (middle) or 4AA (bottom), showing the spectral differences in the aromatic region associated with the different aminobenzoic acid moieties (illustrated schematically above each spectrum). Resonances from residual ammonium ions (from SEC) present in the preparations of HA_6_-3AA and HA_6-_4AA are indicated with +. All spectra were recorded at 25 °C on a spectrometer operating at 500 MHz.

### Analysis of interactions of aminobenzoic-modified HA oligosaccharides with CD44 and TSG-6

The effect of aminobenzoic acid modifications on the affinities of the HA_6_^AN^ and HA_8_^AN^ oligosaccharides for Link_TSG6 was assessed using isothermal titration calorimetry (ITC; **Figure 5**). HA_6_^AN^ modified with 2AA, 3AA, and 4AA aminobenzoic acid moieties were all found to have increased affinity (**Table 1**), with the largest changes observed with 2AA and 3AA (190% and 220% binding, respectively, compared to HA_6_^AN^), driven by favorable changes in entropy compared to the unmodified 6-mer. Conversely, 3AA modification of HA_8_^AN^ did not improve the binding affinity for Link_TSG6, with addition of 2AA and 4AA resulting in a reduction in the *K*_D_ value (16% and 42% of binding, respectively, compared to HA_8_^AN^) due to large unfavorable changes in the entropy of interaction, partially compensated by increased enthalpies (**Table 1**).

**Figure 5.**
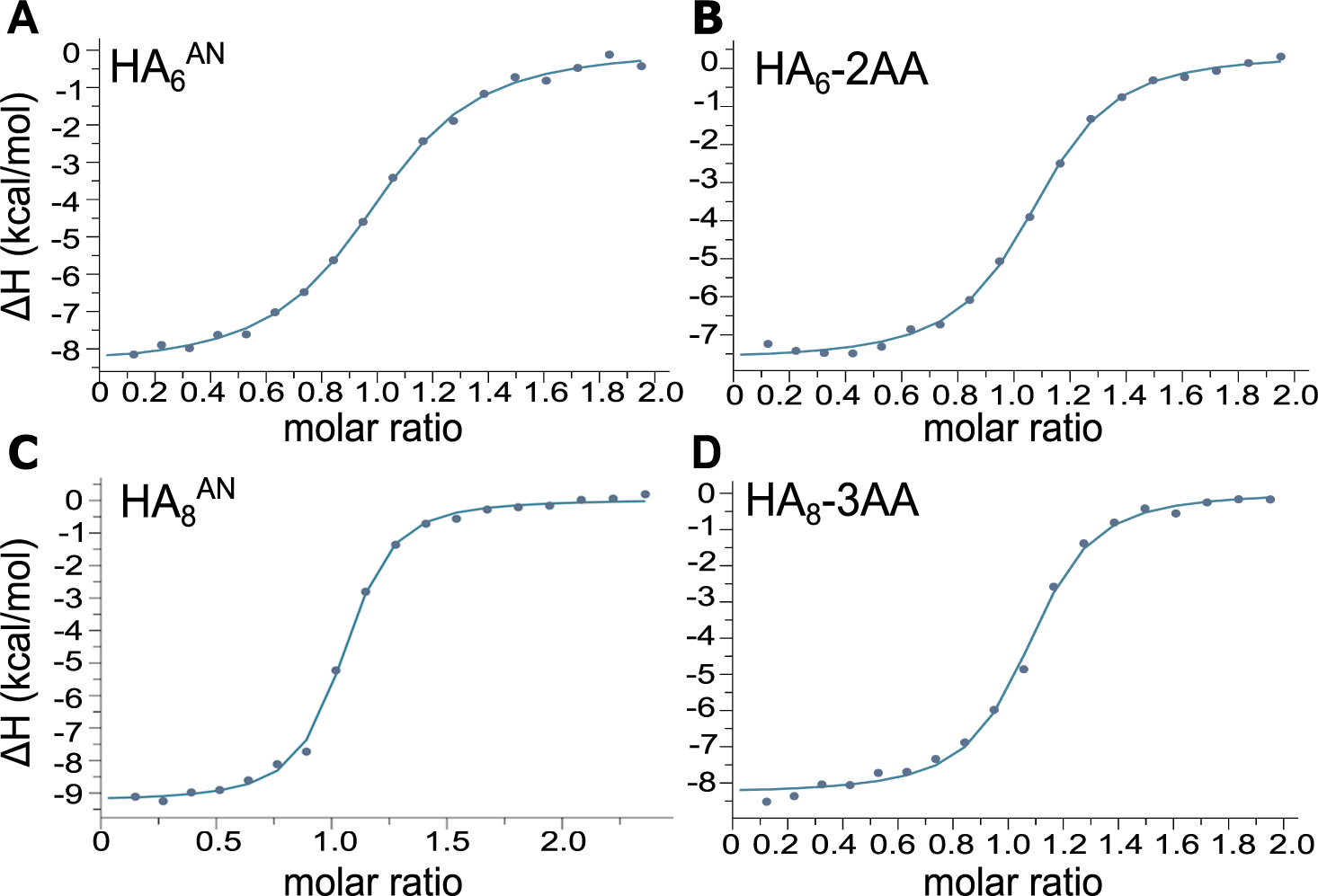
HA_6_-2AA has increased affinity for Link_TSG6 compared to unmodified HA_6_^AN^. Comparison of ITC plots for the interaction of Link_TSG6 with HA oligosaccharides in 5 mM MES, pH 6.0, at 25 °C: **A**) HA_6_^AN^ **B**) HA_6_-2AA **C**) HA_8_^AN^ and **D**) HA_8_-3AA. All titrations were performed with 0.29 mM oligosaccharide solutions injected (18 x 2 µL) into 0.029 mM Link_TSG6 in the cell. In **A-D**, data are fit to a one-site binding model to derive the dissociation constant (**Table 1**). The titrations shown are representative of at least four independently performed experiments for each oligosaccharide.

**Table 1:**
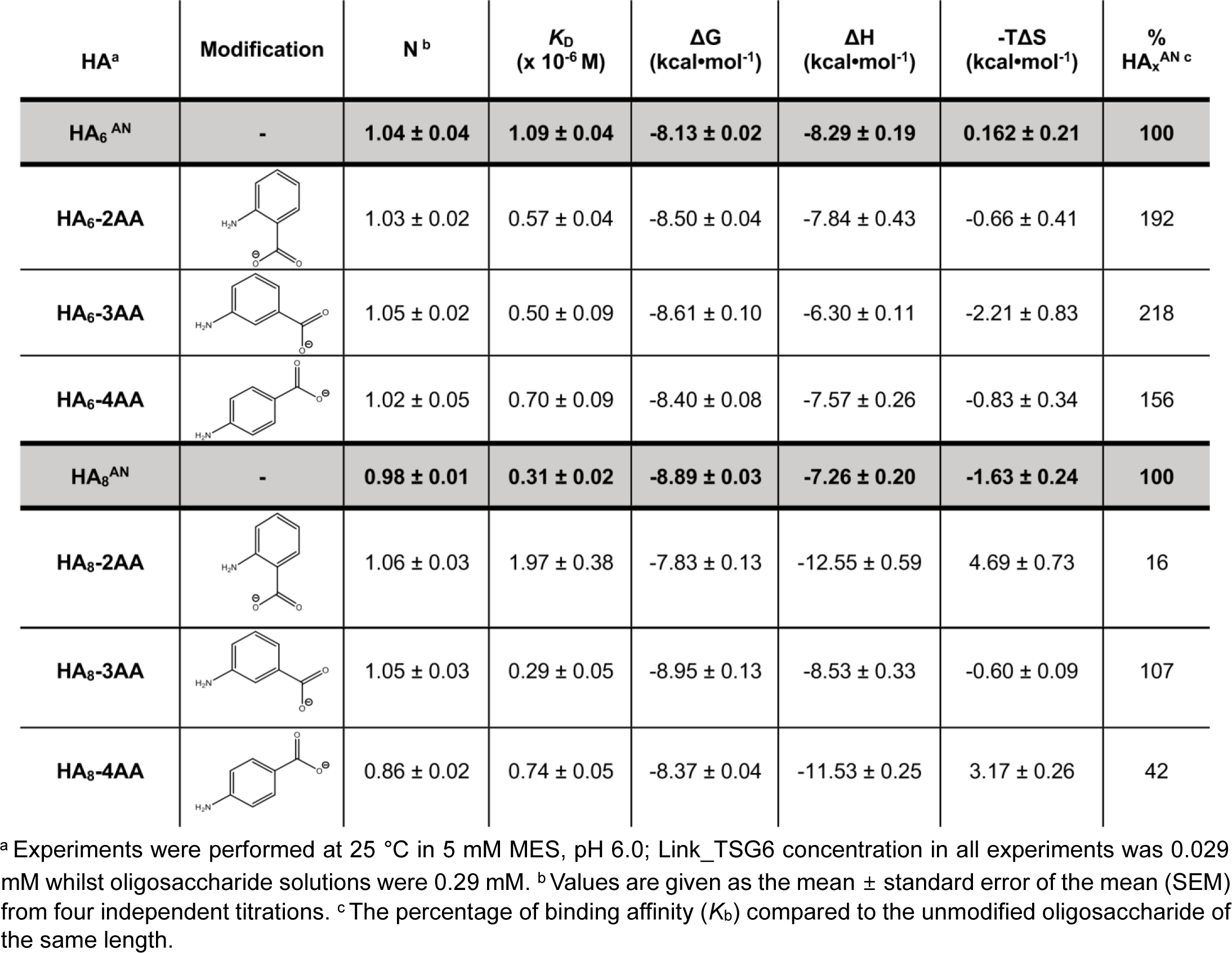
Dissociation constants and thermodynamic parameters measured by ITC for interactions of Link_TSG6 with the 1^st^ series of (aminobenzoic acid) modified HA_6_^AN^ and HA_8_^AN^ oligosaccharides.

| HA<sup>a</sup> | Modification | N<sup>b</sup> | K<sub>D</sub><br>(x 10<sup>-6</sup> M) | ΔG<br>(kcal•mol<sup>-1</sup>) | ΔH<br>(kcal•mol<sup>-1</sup>) | -TΔS<br>(kcal•mol<sup>-1</sup>) | %<br>HA<sub>x</sub><sup>AN c</sup> |
| --- | --- | --- | --- | --- | --- | --- | --- |
| HA<sub>6</sub><sup>AN</sup> | - | 1.04 ± 0.04 | 1.09 ± 0.04 | -8.13 ± 0.02 | -8.29 ± 0.19 | 0.162 ± 0.21 | 100 |
| HA<sub>6</sub>-2AA |  | 1.03 ± 0.02 | 0.57 ± 0.04 | -8.50 ± 0.04 | -7.84 ± 0.43 | -0.66 ± 0.41 | 192 |
| HA<sub>6</sub>-3AA |  | 1.05 ± 0.02 | 0.50 ± 0.09 | -8.61 ± 0.10 | -6.30 ± 0.11 | -2.21 ± 0.83 | 218 |
| HA<sub>6</sub>-4AA |  | 1.02 ± 0.05 | 0.70 ± 0.09 | -8.40 ± 0.08 | -7.57 ± 0.26 | -0.83 ± 0.34 | 156 |
| HA<sub>8</sub><sup>AN</sup> | - | 0.98 ± 0.01 | 0.31 ± 0.02 | -8.89 ± 0.03 | -7.26 ± 0.20 | -1.63 ± 0.24 | 100 |
| HA<sub>8</sub>-2AA |  | 1.06 ± 0.03 | 1.97 ± 0.38 | -7.83 ± 0.13 | -12.55 ± 0.59 | 4.69 ± 0.73 | 16 |
| HA<sub>8</sub>-3AA |  | 1.05 ± 0.03 | 0.29 ± 0.05 | -8.95 ± 0.13 | -8.53 ± 0.33 | -0.60 ± 0.09 | 107 |
| HA<sub>8</sub>-4AA |  | 0.86 ± 0.02 | 0.74 ± 0.05 | -8.37 ± 0.04 | -11.53 ± 0.25 | 3.17 ± 0.26 | 42 |

<sup>a</sup> Experiments were performed at 25 °C in 5 mM MES, pH 6.0; Link\_TSG6 concentration in all experiments was 0.029 mM whilst oligosaccharide solutions were 0.29 mM. <sup>b</sup> Values are given as the mean ± standard error of the mean (SEM) from four independent titrations. <sup>c</sup> The percentage of binding affinity (K<sub>b</sub>) compared to the unmodified oligosaccharide of the same length.

Using NMR spectroscopy, we assessed the chemical shift perturbations of ^15^N-labeled Link_TSG6 in complex with either HA_6_-2AA or HA_8_-2AA (compared to unmodified HA_6_^AN^ and HA_8_^AN^; **Figure 6**); comparison overlays illustrating differences are shown in **Figure S2 and S3,** respectively The overall similarities of the [^1^H-^15^N]-HSQC spectra for Link_TSG6 in the presence of unmodified and modified oligosaccharides indicates that both HA_6_-2AA and HA_8_-2AA were similarly bound in the HA binding groove of Link_TSG6, *i.e.,* based on our previous extensive NMR analyses (39, 41, 42). Consistent with the reducing termini of the oligosaccharides being located near Tyr78 and Arg81 (see **Figure 1**), the largest chemical shift perturbations for Link_TSG6 bound by either HA_6_-2AA or HA_8_-2AA (compared to HA_6_^AN^ and HA_8_^AN^, respectively) were observed in this region (**Figure 6**). For HA_6_-2AA, which has ∼2-fold increased affinity (**Table 1**), most perturbations were localized to the β4-β5 loop, whereas for the longer oligosaccharide HA_8_-2AA (that has ∼6-fold reduced affinity), perturbations were generally larger and localized more widely, *e.g.,* to residues within the β4-β5 and β5-β6 loops, and β6 strand. Our previous NMR analyses revealed that a conformational change in the β4β5 loop was essential to open the binding groove of Link_TSG6 and allow HA to bind (41). Furthermore, ^1^H and ^15^N chemical shift perturbations for ^15^N-Link_TSG6 with HA_4_^AN^, HA_6_^AN^, HA_8_^AN^, and HA_10_^AN^ were all very similar and likewise showed the largest perturbations in the β4-β5 loop. Thus, our finding here that the binding of HA_8_-2AA to Link_TSG6 resulted in additional (large) perturbations in the β5-β6 loop and β6 strand indicates that HA_8_-2AA (and HA_8_-4AA) could induce different local conformational changes (compared to HA_8_^AN^ and HA_8_-3AA), which result in the unfavorable entropy and reduced binding affinities seen for the interactions of Link_TSG6 with modified HA_8_^AN^ oligosaccharides (**Figure 6**, **Table 1** and **Figure S3**).

**Figure 6.**
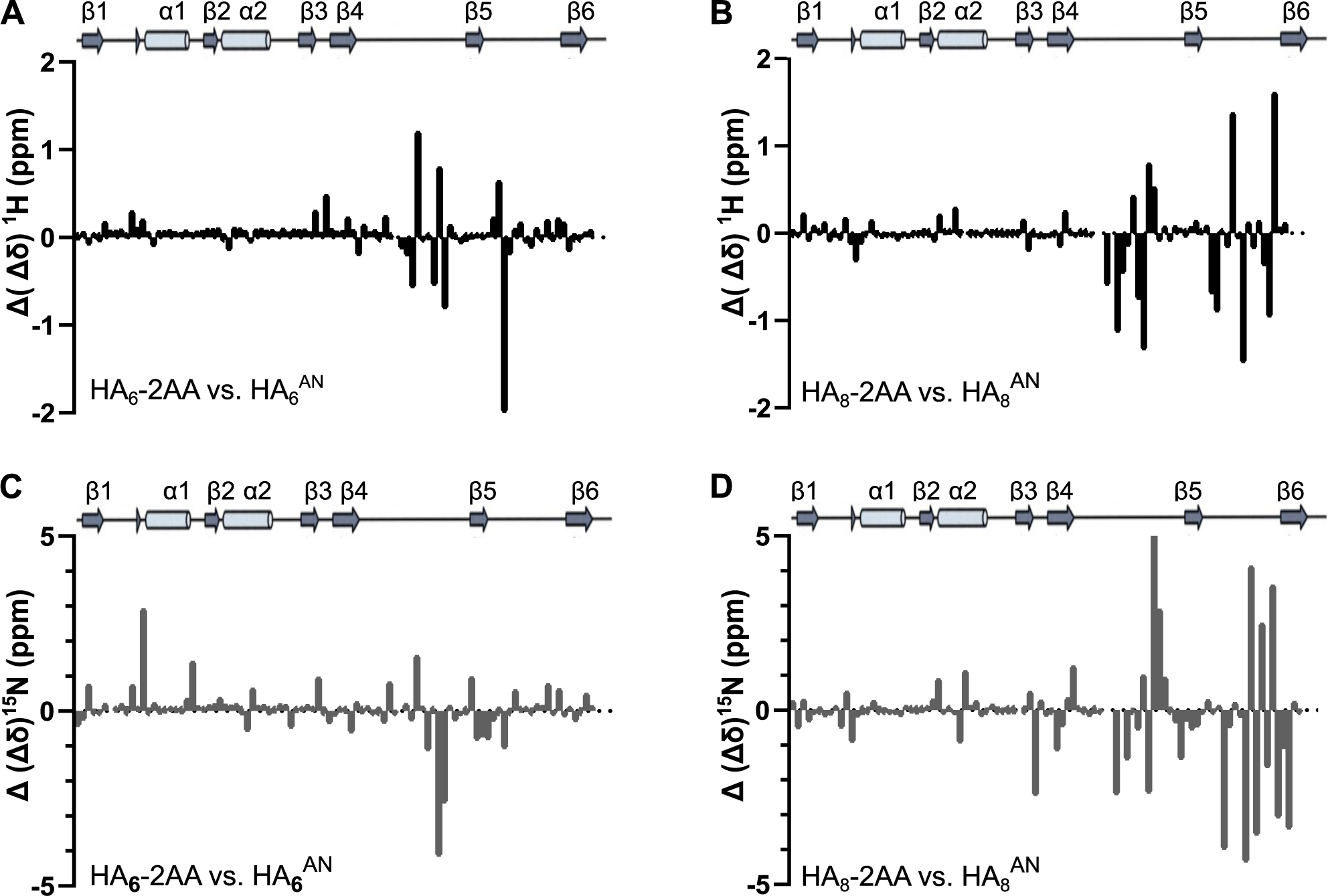
Modified HA_n_-AA oligosaccharides are accommodated in the HA-binding groove of Link_TSG6. Chemical shift perturbations were measured from [^1^H-^15^N]-HSQC spectra recorded on ^15^N-lablled Link_TSG6 in the presence of modified and unmodified HA oligosaccharides (see **Figure S2** and **Figure S3**). Plots show the change (Δ(Δδ)) in ^1^H (**A**, **B**) and ^15^N (**C**, **D**) chemical shifts for Link_TSG6 bound by HA_6_-2AA compared to those induced by binding of HA_6_^AN^ (**A**, **C**) and HA_8_-2AA compared to HA_8_^AN^ (**B**, **D**). In **A-D**, the secondary structure elements of Link_TSG6 (as determined in (41)) are shown above each graph.

We also measured the affinities of the aminobenzoic acid-modified oligosaccharides with CD44. Here a new construct for the HABD of CD44 was generated, comprising residues 20-169 of the human CD44 protein with an N-terminal six histidine tag (named: hisCD44_HABD^20-169^). This removed 9 flexible residues from the C-terminus of the CD44^20-178^ construct, in accordance with our previous studies, which concluded that residues 21-169 define the structural domain for the HABD (32). The 1D-^1^H spectra for hisCD44^20-169^ and CD44^20-178^ (at pH 7.5) were found to be essentially identical and a [^1^H-^15^N]-HSQC recorded on ^15^N-labeled CD44_HABD^20-169^ showed that the backbone amide resonances were well-dispersed with improved resolution in the center of the spectrum compared to the previously studied CD44^20-178^ construct (**Figure S4**).

The affinity of the interaction between HA and the CD44_HABD (from human and mouse) is known to be much weaker than for Link_TSG6 (31, 32, 39, 41). Thus, to overcome the requirement for large amounts of protein and HA oligosaccharides in ITC experiments (31), microscale thermophoresis (MST) was employed instead to study the CD44HA interactions. MST experiments showed that the hisCD44_HABD^20-169^ construct bound to HA_10_^AN^ and HA_8_^AN^ (**Figure 7A, B**) with a similar affinity (∼45 µM and ∼50 µM) to that seen previously for CD44^20-178^ (31). Unmodified HA_4_^AN^ and HA_6_^AN^ oligosaccharides were found to bind with weaker affinities compared to HA_8_^AN^ (**Table 2**), consistent with the observations that oligosaccharides smaller than 8-mers cannot make the full complement of interactions in the HA binding groove (31). For HA_8_^AN^, the 2AA-, 3AA- and 4AA-modifications all reduced the binding affinity for hisCD44_HABD^20-169^ compared to the unmodified oligosaccharide (**Table 2**). HA_8_-4AA had the biggest reduction in binding (to 11% of the HA_8_^AN^ value; **Figure 7D** and **Table 2**). Modified HA_6_^AN^ oligosaccharides (all of which bound more tightly to Link_TSG6; **Table 1**) also had reduced affinity for hisCD44_HABD^20-169^ (11-47% of binding; **Table 2**). Our previous NMR studies have shown that HA_4_^AN^ binds to the CD44_HABD and gives similar chemical shift perturbations in comparison to HA_6_^AN^ and HA_8_^AN^ (32). Here, using MST, we have determined the affinity of the HA_4_^AN^ interaction with hisCD44_HABD^20-169^ as ∼2.2 mM (**Table 2**). We found that modification of HA_4_^AN^ with 4AA led to the greatest increase in the affinity (451% compared to HA_4_^AN^), whereas HA_4_-2AA and HA_4_-3AA increased binding by 183% and 260% respectively.

**Figure 7.**
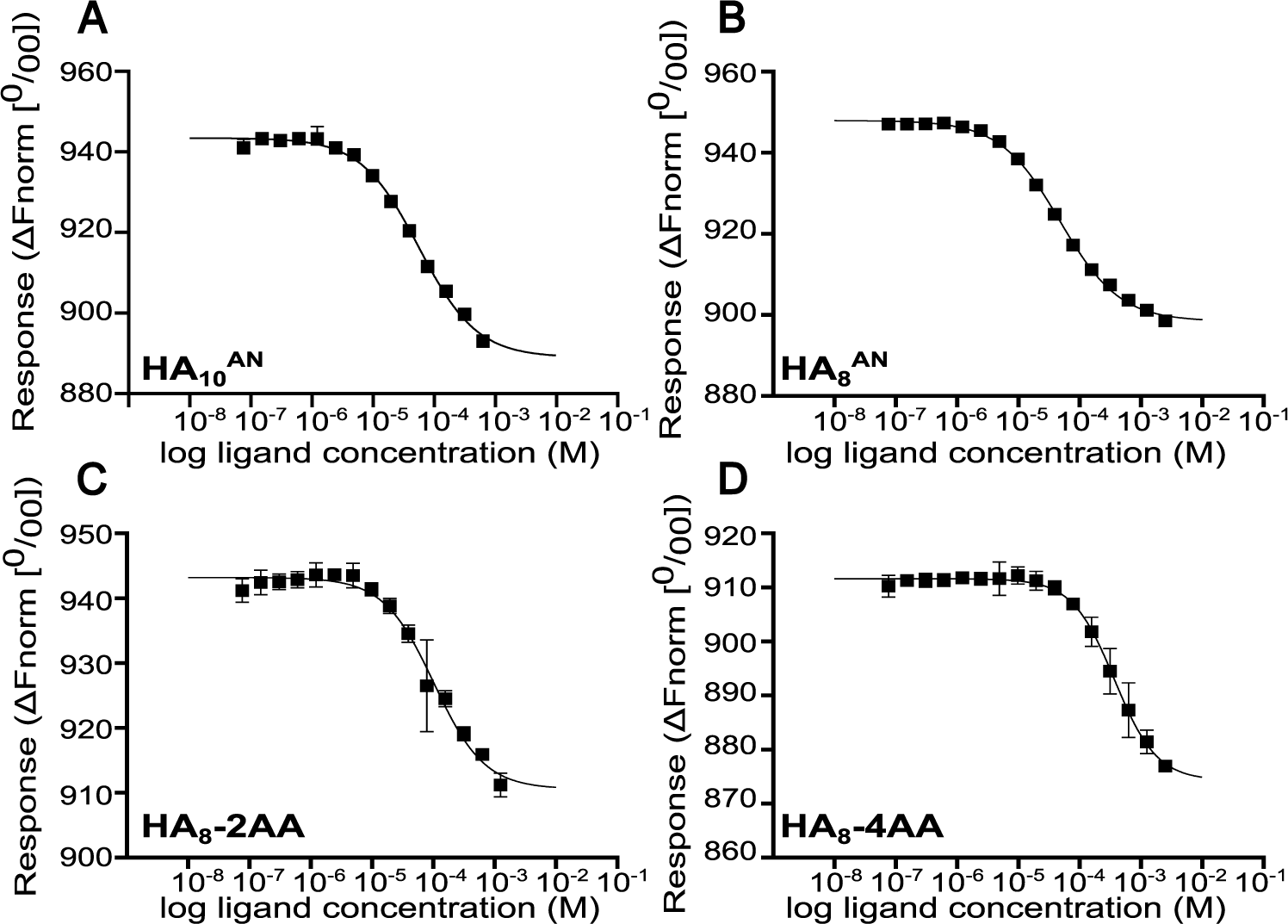
MST analysis of hisCD44_HABD^20-169^ interactions with HA oligosaccharides. Binding curves from MST experiments with hisCD44_HABD^20-169^ and selected HA oligosaccharides: **A**) HA_10_^AN^, **B**) HA_8_^AN^, **C**) HA_8_-2AA, and **D**) HA_8_-4AA. Experiments were performed with 10 nM hisCD44_HABD^20-169^ (labelled with red-Tris-NTA 650 dye) and 5-15 mM HA oligosaccharide in 10 mM HEPES, pH 7.5, 150 mM NaCl and 0.05% Tween-20. All measurements were performed at 25 °C with 20% LED and medium MST power. Plots show the normalized fluorescence change (ΔFnorm [0/00]) as the concentration of HA oligosaccharide is increased, where the data points are the average values (± standard deviation) from three independently performed titration experiments. Data were fit to a 1:1 binding model (65, 99) to derive dissociation constants (**Table 2**) for the interactions.

**Table 2.**
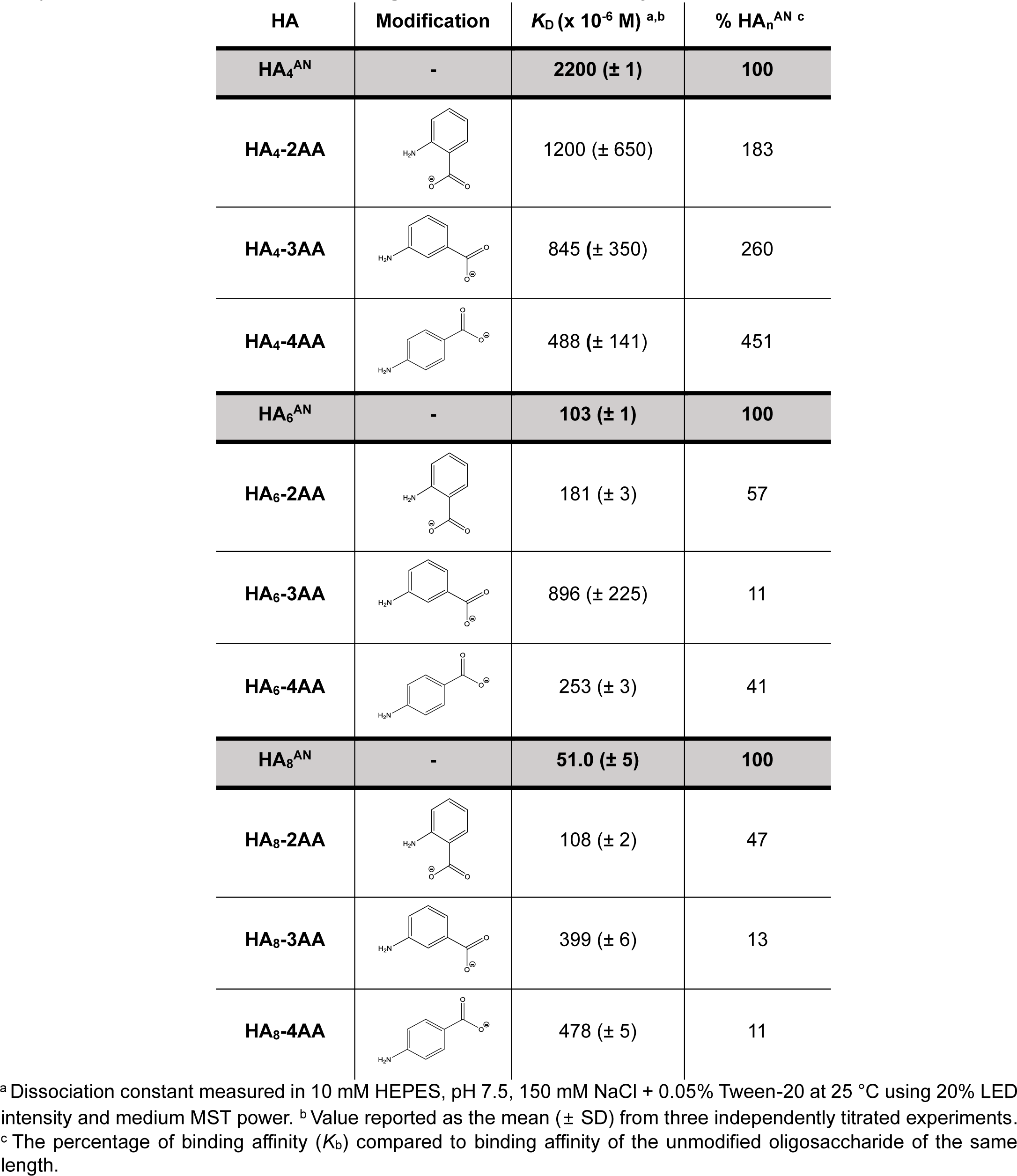
Dissociation constants for interactions of hisCD44_HABD^20-169^ with aminobenzoic acid (2AA, 3AA and 4AA) modified HA_4_^AN^, HA_6_^AN^ and, HA_8_^AN^ oligosaccharides measured by MST.

| HA | Modification | $K_D$ ($\times 10^{-6}$ M)<sup>a,b</sup> | % HA<sub>n</sub><sup>AN</sup><sup>c</sup> |
| --- | --- | --- | --- |
| **HA<sub>4</sub><sup>AN</sup>** | - | **2200 (± 1)** | **100** |
| **HA<sub>4</sub>-2AA** | | 1200 (± 650) | 183 |
| **HA<sub>4</sub>-3AA** | | 845 (± 350) | 260 |
| **HA<sub>4</sub>-4AA** | | 488 (± 141) | 451 |
| **HA<sub>6</sub><sup>AN</sup>** | - | **103 (± 1)** | **100** |
| **HA<sub>6</sub>-2AA** | | 181 (± 3) | 57 |
| **HA<sub>6</sub>-3AA** | | 896 (± 225) | 11 |
| **HA<sub>6</sub>-4AA** | | 253 (± 3) | 41 |
| **HA<sub>8</sub><sup>AN</sup>** | - | **51.0 (± 5)** | **100** |
| **HA<sub>8</sub>-2AA** | | 108 (± 2) | 47 |
| **HA<sub>8</sub>-3AA** | | 399 (± 6) | 13 |
| **HA<sub>8</sub>-4AA** | | 478 (± 5) | 11 |

<sup>a</sup> Dissociation constant measured in 10 mM HEPES, pH 7.5, 150 mM NaCl + 0.05% Tween-20 at 25 °C using 20% LED intensity and medium MST power. <sup>b</sup> Value reported as the mean ( ± SD) from three independently titrated experiments.
<sup>c</sup> The percentage of binding affinity ( $K_b$ ) compared to binding affinity of the unmodified oligosaccharide of the same length.

### Determination of the solution dynamic 3D structure of HA_4_-2AA-modified HA by NMR spectroscopy

As described above, HA_6_-2AA binds to Link_TSG6 with ∼2-fold higher affinity than the unmodified 6-mer HA_6_^AN^. In this regard, the ‘bioactive’ conformation of a ligand when it is bound by protein is often similar to its more populated conformers in solution (55–57). In order to understand the molecular basis of the affinity increase, and therefore enable rational design of modified oligosaccharides with further improved binding, we determined the solution dynamic 3D structure of HA_4_-2AA as the first step towards generating a model of a HA_6_-2AA/Link_TSG6 complex. HA_4_-2AA was chosen for structural analysis because the conformational behavior of the opened reducing terminal GlcNAc ring and attached 2AA moiety was the new information sought, and (by having two fewer saccharide rings) it gave reduced spectral overlap than HA_6_-2AA.

**Figure S5** shows the 2D structure of HA_4_-2AA and the residue and atom nomenclature used. A pH value of 9.4 was found to be optimal for collection of restraints involving the exchangeable aniline (X5 HN11) and amide (T4 HNA1) protons (**Figure S6**). To allow comparison with previously published values for HA_4_^AN^ (58) (**Table S3** and **Table S4),** chemical shifts were assigned at both pH 9.4 in 5% (v/v) D_2_O and pH 6.0 in 100% (v/v) D_2_O; the only significant differences in ^1^H or ^13^C chemical shifts between HA_4_-2AA and HA_4_^AN^ are due to the 2AA modification (**Table S3**, **Table S4** and **Figure S7**). Since the non-reducing terminal 2 residues do not have significantly different ^1^H chemical shift values from those in HA_4_^AN^, the dynamic 3D structure of these residues in HA_4_-2AA is therefore indistinguishable from that determined previously for HA oligosaccharides (59, 60). The negligible differences in chemical shift between pH 9.4 and 6.0 further indicates that the dynamic 3D structure of HA_4_-2AA does not change across this pH range.

The solution dynamic 3D structure of HA_4_-2AA was determined using the methodology described in Blundell *et al.* (55). An internal hydrogen bond was inferred within the 2AA moiety (from aniline X5 HN11 to oxygen X5 O7A) from the low value of the aniline proton temperature coefficient (-2.6 ppb/K; **Table S5**). Conformation-dependent ^3^*J*_HH_ scalar couplings were measured throughout the opened GlcNAc ring (residue T4), and comparison of their values with those for closed GlcNAc rings in HA oligosaccharides (**Table S6**) indicates that the conformational behavior of the opened ring is significantly different; 9 structural restraints were derived from ^3^*J*_HH_ scalar couplings (**Table S7**). Nuclear Overhauser enhancements (NOEs) were measured from a [^1^H-^1^H]-NOESY spectrum and categorized into NOEs (**Table S8**) and noNOEs (**Table S9**) as described previously (55). In total, 144 structural restraints were used to determine the solution dynamic 3D structure of residues G3-T4-X5 of HA_4_-2AA (**Table S10**).

Overall, the solution dynamic 3D behavior of a total of 11 rotatable bonds were measured in the G3-X5 terminus of HA_4_2AA. The model parameters for each torsion in these residues, which define the overall shape of the molecule and best fit the experimental data, are given in **Table S11**, along with the complete dynamic model for the whole of HA_4_-2AA (**Figure S8**). Many of the torsions in the backbone of the molecule adopt unimodal behaviors, giving these portions limited local shapes. Three torsions in the opened GlcNAc ring (T4 C2-C3, T4 C1-C2 and T4 C1-X5 HN11), however, display more complex, co-dependent conformational behavior, and, since these comprise part of the backbone of the molecule, their behavior strongly affects the reducing terminal’s gross shape. The co-dependency of these torsions means that as one torsion adopts a particular value, the others move in tandem to accommodate it, and, overall, all three torsions together manifest 5 modes, producing 5 distinct shapes (Conformers 1-5) about which the molecule librates in solution (**Table 3**).

**Table 3.** Co-dependent conformational behavior of torsions within residues T4-X5 of HA_4_-2AA.

| T4 C2-C3<sup>a</sup> | T4 C1-C2 | T4 C1-X5 NH11 | Population, $\pi^d$ | Mode |
| --- | --- | --- | --- | --- |
| $\mu = 60^\circ$ <sup>b</sup> $\sigma = 20.0$ <sup>c</sup> | $\mu = -60^\circ$<br>$\sigma = 4.7$ | $\mu = -127^\circ$<br>$\sigma = 4.4$ | 0.35 | 1<br>(Figure 8C) |
| $\mu = 180^\circ$<br>$\sigma = 20.0$ | | $\mu = 116.9^\circ$ $\sigma = 4.4$ | 0.35 | 2<br>(Figure 8D) |
| | | $\mu = -95.8^\circ$<br>$\sigma = 4.4$ | 0.22 | 3<br>(Figure 8E) |
| $\mu = -60^\circ$<br>$\sigma = 20.0$ | $\mu = 180^\circ$<br>$\sigma = 4.7$ | $\mu = -146.9^\circ$<br>$\sigma = 4.4$ | 0.06 | 4<br>(Figure 8F) |
| | | $\mu = 85.3^\circ$<br>$\sigma = 4.4$ | 0.02 | 5<br>(Figure 8G) |

<sup>a</sup> Torsion angle as defined in **Table S11**. <sup>b</sup> $\mu$ is the torsion mean angle of a given mode. <sup>c</sup> $\sigma$ is the libration about the mean angle for a given mode. <sup>d</sup> $\pi$ is the occupancy of a given mode, expressed as a fraction of the total for the whole co-dependent system.

In **Figure 8**, the dynamic 3D structure of HA_4_-2AA (represented as ensembles of conformations) is compared to that of HA_5_^AA^ (60), *i.e.,* an unmodified HA of equivalent in length. In Conformers 1, 2 and 3 (combined population = 0.91; **Figure 8C, D, E**) the T4 GlcNAc acetamido group is in a very similar position to that of native HA, whereas it is considerably relocated in Conformers 4 and 5 (combined population = 0.09; **Figure 8F, G**). The position of the 2AA carboxylate group is different from HA_5_^AA^ in all conformers of HA_4_^AN^-2AA, although Conformers 1, 2 and 3 position it most similarly. The marked conformational change in torsion T4 C3-C4 of the opened GlcNAc ring relative to the closed ring of unmodified HA positions the hydroxyl groups (O4, O5 and O6) in T4 in a very different arrangement (**Figure 8** and **Figure 9**). As a result, it is probable that in HA_4_-2AA, the T4 HO5 rather than T4 HO4 hydroxyl (61) forms the inter-residue hydrogen bond with the anomeric oxygen of the previous GlcA ring.

**Figure 8.**
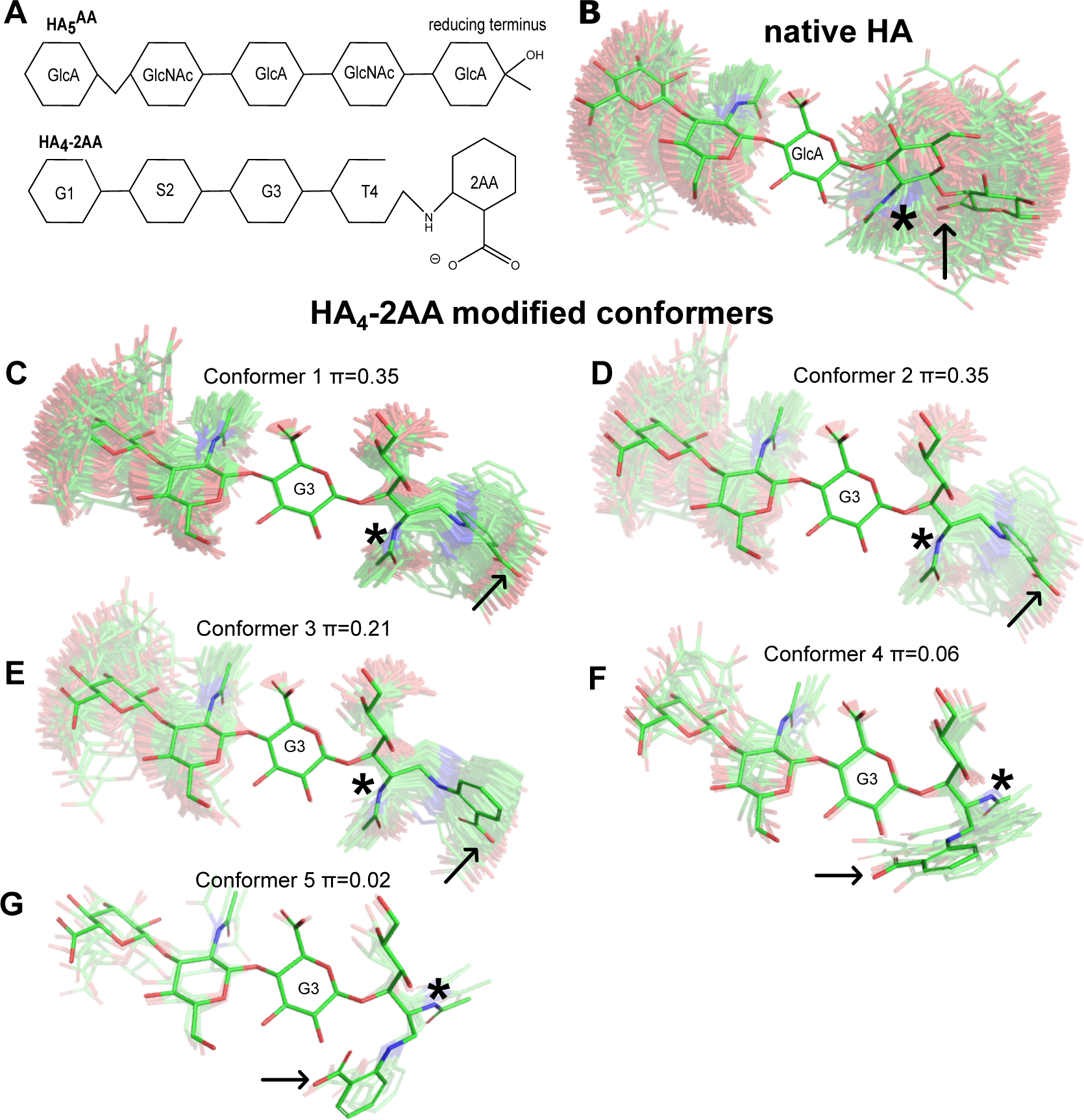
Comparison of solution conformational behavior of HA_4-_2AA with HA_5_AA. **A)** 2D structures of HA_5_^AA^ and HA_4_-2AA, showing residue nomenclature. **B**) Unmodified HA_5_^AA^ adopts one conformer in solution (bright conformation) about which it liberates (faded conformations); 250 conformations are shown, determined as described in (55). **C-F**) An ensemble of 250 conformations were determined for HA_4_^AN^-2AA where these adopt 5 different conformers (Conformer 1-5) of differing occupancies (π). Conformers 1-5 are each shown as the mean conformation (bright) along with their associated librational range (faded). All conformations are overlaid on the ring atoms of the central GlcA residue (G3 in HA_4_-2AA). Carbon atoms are shown in green, oxygen atoms in red and nitrogen atoms in blue; hydrogen atoms are omitted for clarity.

**Figure 9:**
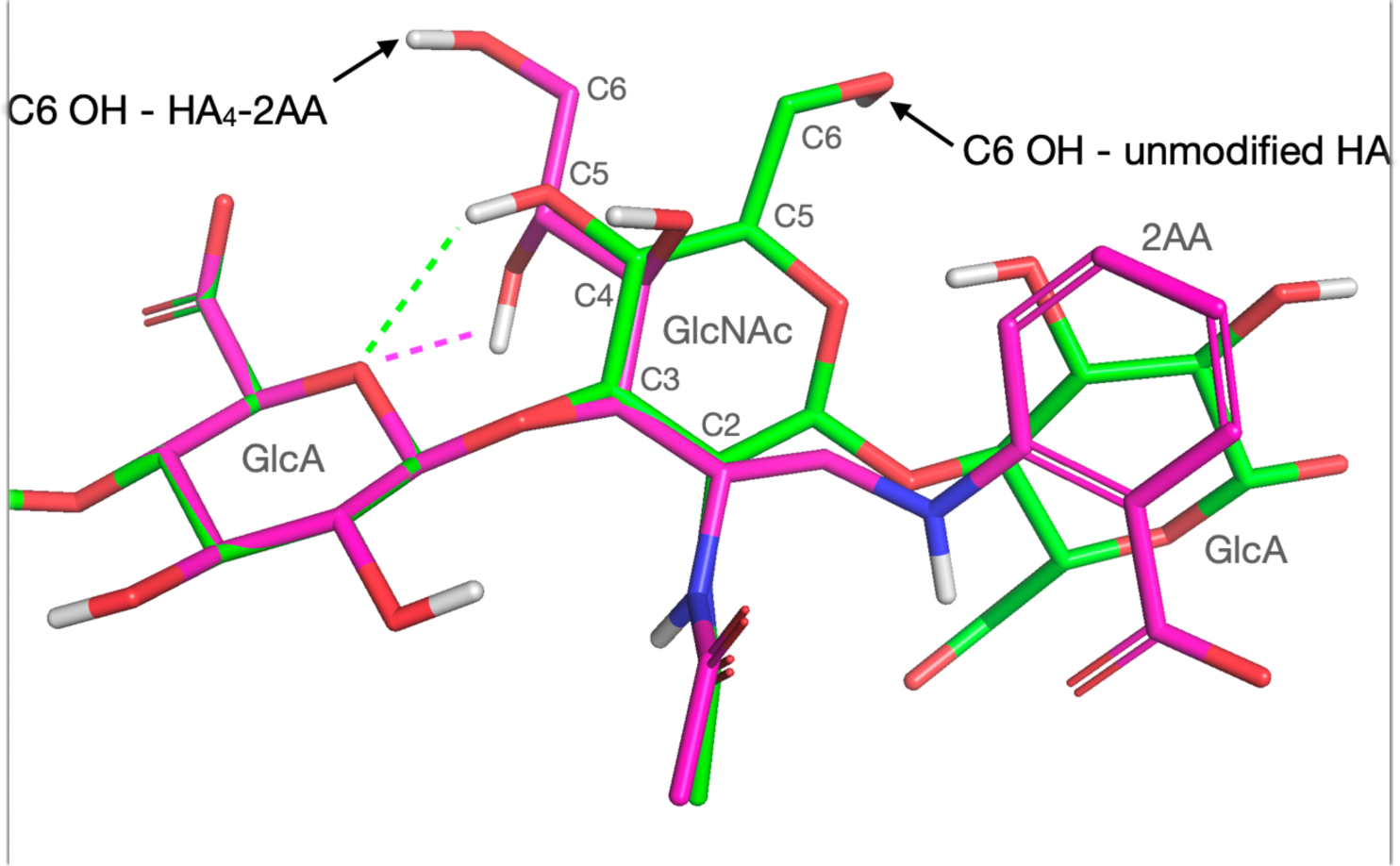
Change in conformation of the GlcNAc ring and its hydroxyl groups caused by the chemical modification. Conformer 3 of the reducing terminus of HA_4_-2AA (with the most populated mode of T4 C5-C6 (180°)) compared to that of the equivalent residues in unmodified HA (60). The change in position of the C6 hydroxyl (C6 OH) group is indicated with arrows. Carbon atoms are shown in magenta for HA_4_-2AA and in green for unmodified HA. All oxygen, nitrogen and hydrogen atom are shown in red, blue, and white, respectively. The conformations are overlaid on the heavy atoms of the internal GlcA of unmodified HA and HA_4_-2AA (*i.e*., G3).

### Generation of a model of the HA_6_-2AA/Link_TSG6 complex

To investigate how the new reducing terminal of HA_6_-2AA could be accommodated in the HA-binding groove of Link_TSG6 and thereby rationalize the changes in affinity observed with the 2-aminobenzoic acid modified oligosaccharides, a molecular model of the HA_6_-2AA/Link_TSG6 complex was made. Keeping the bound conformation of the non-reducing terminal 4 residues from HA_8_^AN^, the dynamic 3D structure of the opened ring and 2AA moiety of HA_6_-2AA was modeled according to the solution behavior determined for HA_4_-2AA (**Figure S9**). The resulting ensemble of 250 conformations of HA_6_-2AA were then individually docked with rigid body docking into the lowest-energy HA-bound conformation of Link_TSG6 (accession code: 1o7c; (41)).

Analysis of the docking results revealed that HA_6_-2AA conformations representing libration around Conformer 2 could locate the X5 carboxylate group in the most similar position to that of the unmodified bound HA (**Figure 10A**, **B** and **Figure S9**). Indeed, when the 250 conformations of HA_6_-2AA were evaluated using GoldScore (62)), conformations from the Conformer 2 librational distribution gave the highest values (**Figure 10C**), while some conformations from Conformers 1 and 3 also gave good scores. Librations from Conformers 4 and 5, however, clearly gave poorer complementarity. In the top-ranking conformation of HA_6_-2AA (a conformation from the librational distribution of Conformer 2) the backbone of the opened GlcNAc residue (T4) lies against Tyr78, possibly contributing to hydrophobic interactions, while the carboxylate group of the 2AA moiety is orientated towards Arg81 of Link_TSG6, allowing it to make a salt bridge (**Figure 10D**); this is consistent with the favorable enthalpy changes and overall increased affinity seen in the ITC experiments (**Table 1**). This model of the HA_6_-2AA/Link_TSG6 complex also suggests that the aromatic ring of the 2AA group can be accommodated in a hydrophobic ‘specificity’ pocket (see below), which accommodates the methyl group of GlcNAc in unmodified HA (39, 41, 42).

**Figure 10:**
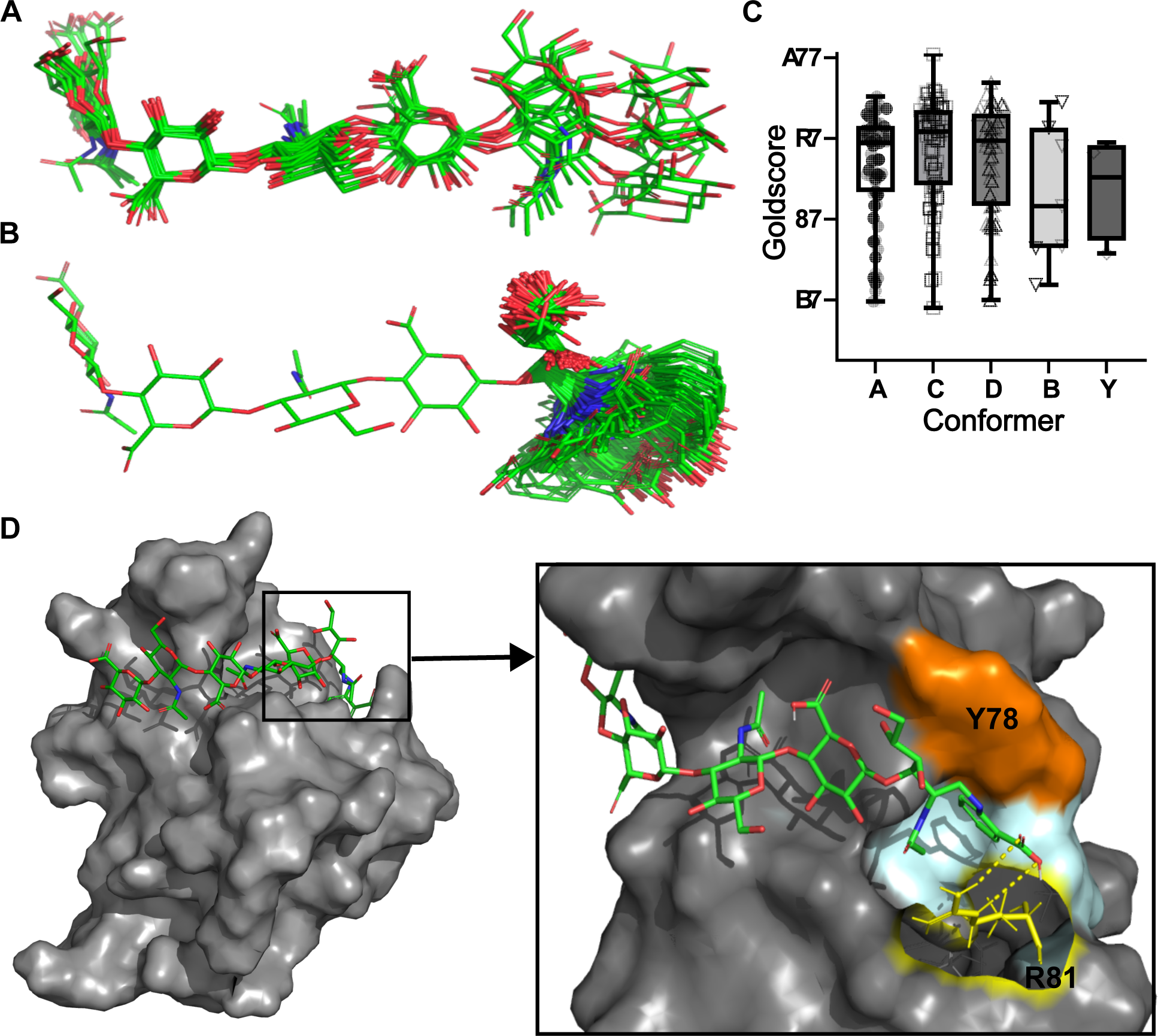
Modeling the interaction of HA_6_-2AA with Link_TSG6. **A**) The 10 conformations of HA_8AN_ bound to Link_TSG6 taken from (39), showing only residues 1-7 to correspond to the size and composition of HA_6_-2AA. **B**) Subset of the ensemble of model conformations of HA_6_-2AA (see **Figure S9**) showing conformations representing libration around Conformer 2 (Figure 8). **C**) GoldScore scores for each of the 250 conformations of modeled HA_6_-2AA, grouped by the Conformer they librate around (see Figure 8); higher scores represent more favorable molecular interactions. Boxplots show the mean score and interquartile range per Conformer group, whiskers indicate the minimum and maximum score per group. **D**) Rigid-body docking of the highest ranked HA_6_-2AA conformation (a libration from Conformer 2) into the lowest energy HA_8_^AN^-bound conformation of Link_TSG6. In **A**), **B**) and **D**), carbon atoms are shown in green, oxygen atoms in red, and nitrogen atoms in blue; hydrogen atoms are omitted for clarity. In **D**) the protein surface is shown in grey with key HA-binding residues (Tyr78 (Y78) and Arg81 (R81)) labelled; Y78 is colored orange, and carbon atoms of the exposed side chain of R81 are shown in yellow. The hydrogen-bonds of the potential salt-bridge to the X5 carboxylate of HA_6_-2AA are indicated by dashed yellow lines (nitrogen atoms of R81 are shown in blue and hydrogen atoms in white). The hydrophobic pocket (comprised of sidechains from Val57, Gly79 and Ile80) where the 2AA group could be accommodated is colored light blue.

### Rational design of a 2^nd^ series of modified HA oligosaccharides and their interactions with CD44_HABD and Link_TSG6

Modeling of the interaction of HA_6_-2AA with Link_TSG6 (described in the previous section), revealed that the aromatic ring of 2AA could potentially be accommodated in a hydrophobic pocket formed by residues Gly79, Ile80, Arg81, Leu82, Val57 and Trp88 of Link_TSG6 (**Figure 10D**). Thus, we hypothesized that further increases in binding affinity to Link_TSG6 could be achieved by modifications to the reducing terminus of HA that either a) enhanced the salt bridge with Arg81 and/or b) promoted aromatic ring stacking or hydrophobic interactions with the specificity pocket. In order to test whether the salt bridge could be strengthened, we modified HA_n_^AN^ with 2-amino-4-methoxybenzoic acid (2A4MBA). This group was selected because the acidity of the carboxylate is increased by the presence of the methoxy group in the *para* position on the benzoic acid ring. The HA-2A4MBA oligosaccharides (and the other ‘2^nd^ series’ modified oligomers) were generated and characterized as described before (**Table S1** and **Table S2**). Using ITC, we found that HA_6_-2A4MBA indeed had increased affinity for Link_TSG6 compared to HA_6_^AN^ (238% of binding) and also had a lower *K*_D_ value and increased enthalpy (**Table 4**) compared to the interaction with HA_6_-2AA (**Table 1**); *i.e.,* consistent with our intended design to increase the strength of the salt bridge. Conversely, the longer oligosaccharide, HA_8_-2A4MBA had reduced affinity for Link_TSG6 (21% of binding; **Table 4**), which was a similar reduction to that seen with HA_8_-2AA (**Table 1**).

**Table 4.**
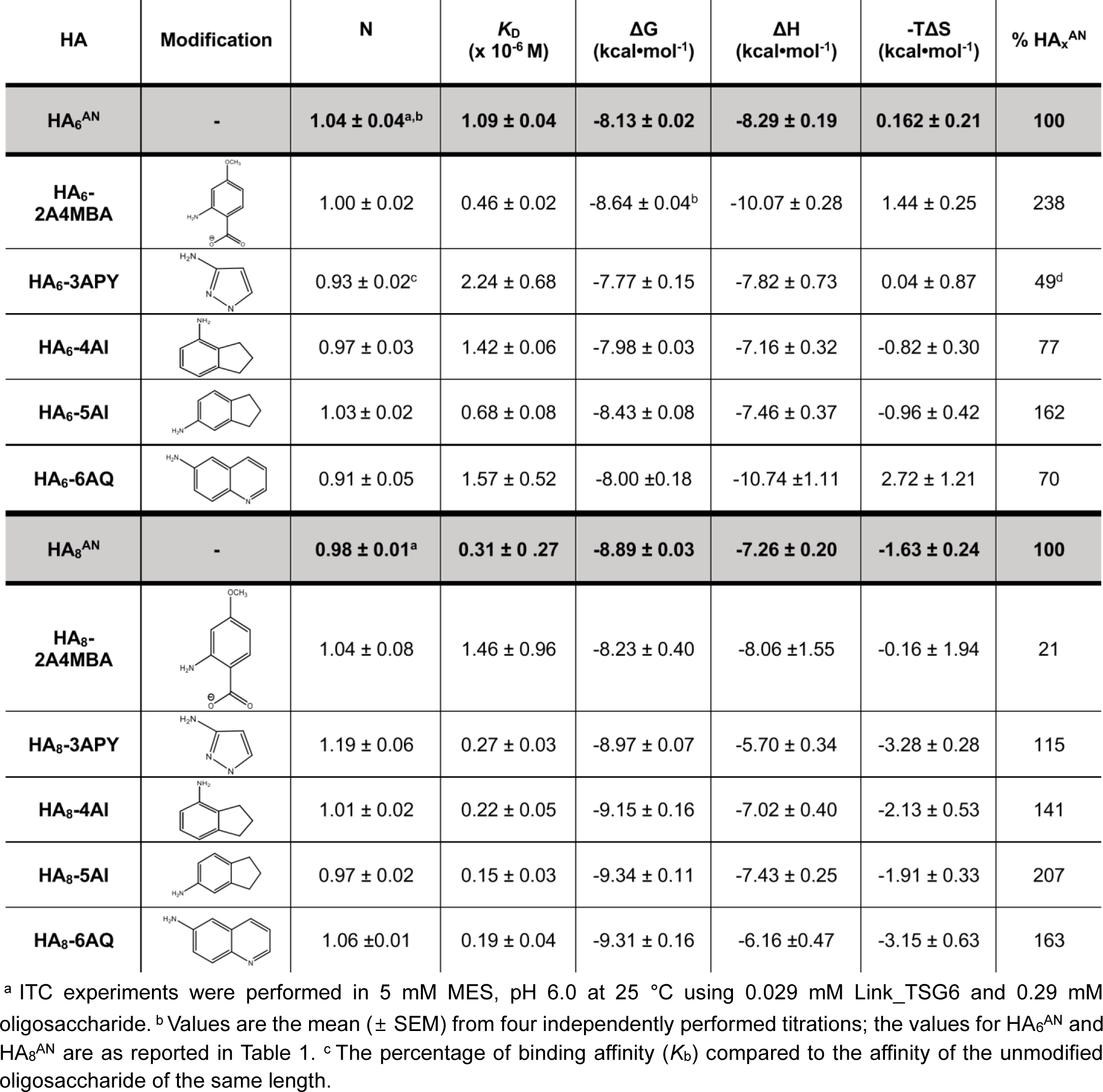
Dissociation constants and thermodynamic parameters measured by ITC for interactions of Link_TSG6 with the 2^nd^ series of modified HA_6_^AN^ and HA_8_^AN^ oligosaccharides.

| HA | Modification | N | $K_D$<br>($\times 10^{-6}$ M) | $\Delta G$<br>(kcal·mol<sup>-1</sup>) | $\Delta H$<br>(kcal·mol<sup>-1</sup>) | -T$\Delta S$<br>(kcal·mol<sup>-1</sup>) | % HA<sub>x</sub><sup>AN</sup> |
| --- | --- | --- | --- | --- | --- | --- | --- |
| HA<sub>6</sub><sup>AN</sup> | - | 1.04 ± 0.04<sup>a,b</sup> | 1.09 ± 0.04 | -8.13 ± 0.02 | -8.29 ± 0.19 | 0.162 ± 0.21 | 100 |
| HA<sub>6</sub>-2A4MBA |  | 1.00 ± 0.02 | 0.46 ± 0.02 | -8.64 ± 0.04<sup>b</sup> | -10.07 ± 0.28 | 1.44 ± 0.25 | 238 |
| HA<sub>6</sub>-3APY |  | 0.93 ± 0.02<sup>c</sup> | 2.24 ± 0.68 | -7.77 ± 0.15 | -7.82 ± 0.73 | 0.04 ± 0.87 | 49<sup>d</sup> |
| HA<sub>6</sub>-4AI |  | 0.97 ± 0.03 | 1.42 ± 0.06 | -7.98 ± 0.03 | -7.16 ± 0.32 | -0.82 ± 0.30 | 77 |
| HA<sub>6</sub>-5AI |  | 1.03 ± 0.02 | 0.68 ± 0.08 | -8.43 ± 0.08 | -7.46 ± 0.37 | -0.96 ± 0.42 | 162 |
| HA<sub>6</sub>-6AQ |  | 0.91 ± 0.05 | 1.57 ± 0.52 | -8.00 ± 0.18 | -10.74 ± 1.11 | 2.72 ± 1.21 | 70 |
| HA<sub>8</sub><sup>AN</sup> | - | 0.98 ± 0.01<sup>a</sup> | 0.31 ± 0.27 | -8.89 ± 0.03 | -7.26 ± 0.20 | -1.63 ± 0.24 | 100 |
| HA<sub>8</sub>-2A4MBA |  | 1.04 ± 0.08 | 1.46 ± 0.96 | -8.23 ± 0.40 | -8.06 ± 1.55 | -0.16 ± 1.94 | 21 |
| HA<sub>8</sub>-3APY |  | 1.19 ± 0.06 | 0.27 ± 0.03 | -8.97 ± 0.07 | -5.70 ± 0.34 | -3.28 ± 0.28 | 115 |
| HA<sub>8</sub>-4AI |  | 1.01 ± 0.02 | 0.22 ± 0.05 | -9.15 ± 0.16 | -7.02 ± 0.40 | -2.13 ± 0.53 | 141 |
| HA<sub>8</sub>-5AI |  | 0.97 ± 0.02 | 0.15 ± 0.03 | -9.34 ± 0.11 | -7.43 ± 0.25 | -1.91 ± 0.33 | 207 |
| HA<sub>8</sub>-6AQ |  | 1.06 ± 0.01 | 0.19 ± 0.04 | -9.31 ± 0.16 | -6.16 ± 0.47 | -3.15 ± 0.63 | 163 |

<sup>a</sup> ITC experiments were performed in 5 mM MES, pH 6.0 at 25 °C using 0.029 mM Link\_TSG6 and 0.29 mM oligosaccharide. <sup>b</sup> Values are the mean (± SEM) from four independently performed titrations; the values for HA<sub>6</sub><sup>AN</sup> and HA<sub>8</sub><sup>AN</sup> are as reported in Table 1. <sup>c</sup> The percentage of binding affinity ( $K_b$ ) compared to the affinity of the unmodified oligosaccharide of the same length.

With the aim of enhancing hydrophobic interactions between modified HA oligosaccharides and Link_TSG6, HA_6_^AN^ and HA_8_^AN^ were modified with 3-aminopyrazole (3APY), 4-aminoindan (4AI), 5-aminoindan (5AI) and 6-amino-quinolone (6AQ; **Table S1** and **Table S2**). HA_6_-5AI had an increased affinity for Link_TSG6 (∼162% binding compared to HA_6_^AN^), which was driven by a favorable change in entropy (**Table 4**). Modification of HA_6_^AN^ with 4AI, 6AQ or 3APY did not enhance the interaction for Link_TSG6 compared to HA_6_^AN^ (**Table 4**); in the case of HA_6_-6AQ (70% of the affinity with HA_6_^AN^), there was an increase in enthalpy, but this was offset by a large unfavorable change in entropy. Modification of HA_8_^AN^ with 3APY, 4AI, 5AI, and 6AQ all resulted in increased affinities for Link_TSG6 (**Table 4**), with the largest gain seen with 5AI (HA_8_-5AI; 207% of binding affinity compared to HA_8_^AN^). HA_8_-5AI has the highest overall affinity of any of the modified oligosaccharides generated in this study, due to favorable changes in both enthalpy and entropy when compared to the unmodified HA_8_^AN^.

The interactions of the 2^nd^ series of modified oligosaccharides with hisCD44_HABD^20-169^ were also assessed using MST (**Table 5**); except for HA_n_^AN^ modified with 6AQ, which could not be analyzed by MST due to fluorescence interference from the aminoquinoline group (63–65). We found that modification of HA_6_^AN^ with 5AI resulted in increased affinity for hisCD44_HABD^20-169^ (133% of binding compared to the unmodified 6-mer), whereas the 2A4MBA and 4AI modifications reduced binding affinity (52% and 28% of HA_6_^AN^ binding, respectively), and modification with 3APY did not greatly alter the affinity of the interaction (91% compared to HA_6_^AN^). Apart from 2A4MBA (which increased binding by 141%), none of the modifications to HA_8_^AN^ resulted in improved binding affinities for hisCD44_HABD^20-169^, with HA_8_-3APY, HA_8_-4AI and HA_8_-5AI exhibiting 23, 37 and 96%, respectively, compared to the unmodified HA oligosaccharide.

**Table 5.**
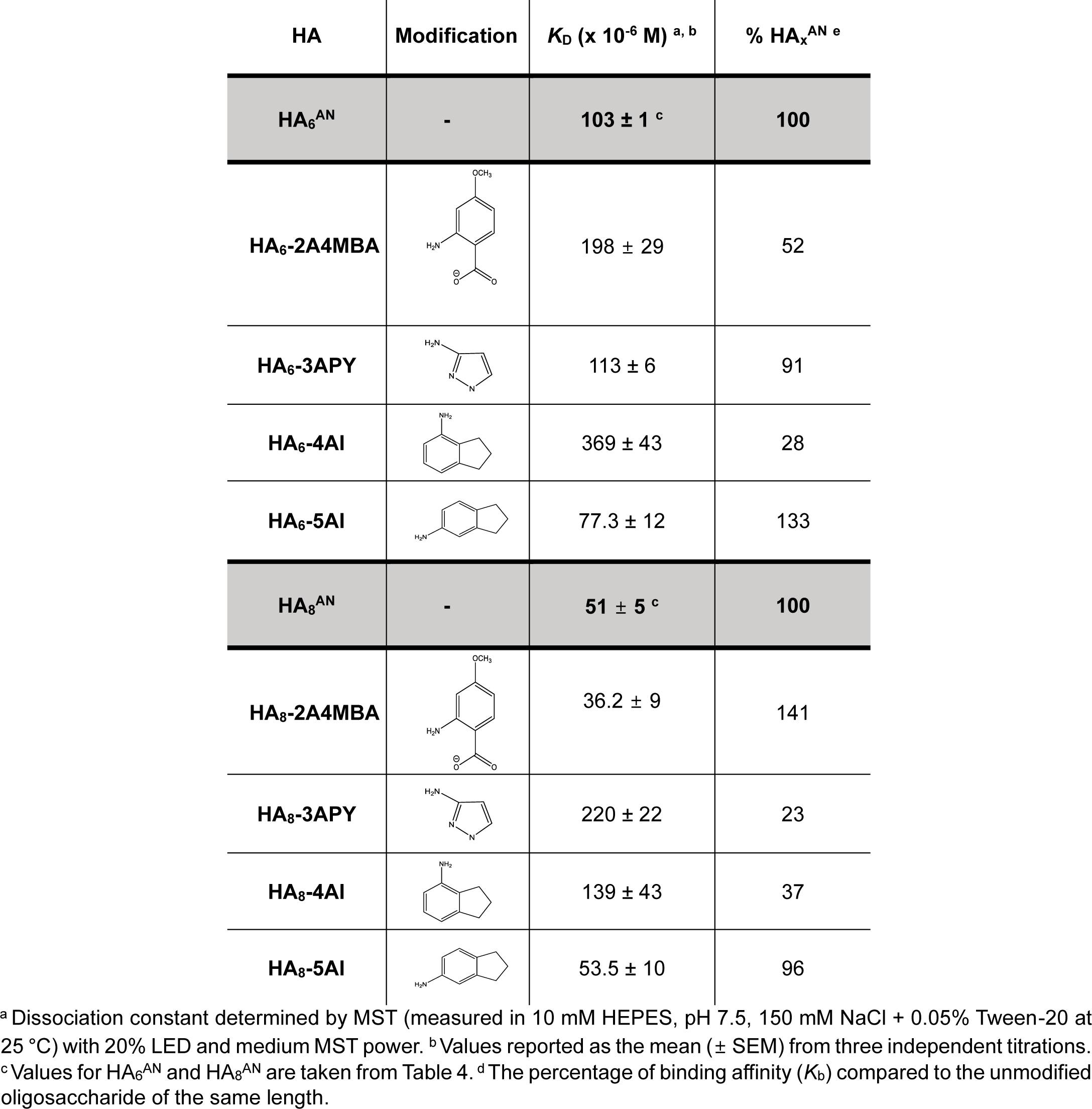
Dissociation constants measured by MST for interactions of hisCD44_HABD^20-169^ with the 2^nd^ series of modified HA_6_^AN^ and HA_8_^AN^ oligosaccharides.

| HA | Modification | $K_D$ ($\times 10^{-6}$ M) <sup>a, b</sup> | % HA<sub>x</sub><sup>AN</sup> <sup>e</sup> |
| --- | --- | --- | --- |
| HA<sub>6</sub><sup>AN</sup> | - | 103 ± 1 <sup>c</sup> | 100 |
| HA<sub>6</sub>-2A4MBA |  | 198 ± 29 | 52 |
| HA<sub>6</sub>-3APY |  | 113 ± 6 | 91 |
| HA<sub>6</sub>-4AI |  | 369 ± 43 | 28 |
| HA<sub>6</sub>-5AI |  | 77.3 ± 12 | 133 |
| HA<sub>8</sub><sup>AN</sup> | - | 51 ± 5 <sup>c</sup> | 100 |
| HA<sub>8</sub>-2A4MBA |  | 36.2 ± 9 | 141 |
| HA<sub>8</sub>-3APY |  | 220 ± 22 | 23 |
| HA<sub>8</sub>-4AI |  | 139 ± 43 | 37 |
| HA<sub>8</sub>-5AI |  | 53.5 ± 10 | 96 |

<sup>a</sup> Dissociation constant determined by MST (measured in 10 mM HEPES, pH 7.5, 150 mM NaCl + 0.05% Tween-20 at 25 °C) with 20% LED and medium MST power. <sup>b</sup> Values reported as the mean ( ± SEM) from three independent titrations. <sup>c</sup> Values for HA<sub>6</sub><sup>AN</sup> and HA<sub>8</sub><sup>AN</sup> are taken from Table 4. <sup>d</sup> The percentage of binding affinity ( $K_b$ ) compared to the unmodified oligosaccharide of the same length.

One of the major goals of this study was to generate modified HA oligosaccharides that differentially modulated binding to CD44 and TSG-6 (summarized in **Table 6**). The largest differential was seen for HA_6_-3AA (218% binding for Link_TSG6 *vs* 11% binding for hisCD44_HABD^20-169^), resulting in a 20-fold relative change in affinity for Link_TSG6 over CD44_HABD. HA_8_-3AA showed the second biggest differential effect with a nearly 10-fold change overall (107% binding to Link_TSG6 *vs* 11% binding to hisCD44_HABD^20-169^). Conversely, modification of HA_8_^AN^ with 2A4MBA best improved the binding for CD44_HABD compared to Link_TSG6 (∼7-fold; 21% of binding to Link_TSG6 *vs* 141% binding to hisCD44_HABD^20-169^). Together these data demonstrate that it is indeed feasible to make chemical modifications to HA that enhance the binding affinity and differentially modulate HA’s interactions with particular HABPs.

**Table 6.** Summary of relative binding affinities for the interactions of modified HA oligosaccharides with hisCD44_HABD^20-169^ and Link_TSG6, and substrate activity of the modified oligomers in HC-transfer assays.

|  | HA | % Binding<br>affinity<br>Link_TSG6 | % Binding<br>affinity<br>hisCD44_HABD<sup>20-169</sup> | Relative discrimination<br>Link_TSG6:<br>hisCD44_HABD<sup>20-169</sup> | Substrate for<br>HC-transfer |
| --- | --- | --- | --- | --- | --- |
|  | HA<sub>4</sub><sup>AN</sup> | n.d<sup>a</sup> | 100 | n.d | No |
| modified<br>HA<sub>4</sub><sup>AN</sup> | HA<sub>4</sub>-2AA | n.d | 183<sup>c</sup> | - | No<sup>d</sup> |
|  | HA<sub>4</sub>-3AA | n.d | 260 | - | No |
|  | HA<sub>4</sub>-4AA | n.d | 451 | - | Yes |
|  | HA<sub>4</sub>-2A4MBA | n.d | n.d | - | Yes |
|  | HA<sub>6</sub><sup>AN</sup> | 100 | 100 | 1.00 | No |
| modified<br>HA<sub>6</sub><sup>AN</sup> | HA<sub>6</sub>-2AA | 192<sup>b</sup> | 57 | 3.40 | Yes |
|  | HA<sub>6</sub>-3AA | 218 | 11 | 19.80 | No |
|  | HA<sub>6</sub>-4AA | 156 | 41 | 3.80 | Yes |
|  | HA<sub>6</sub>-2A4MBA | 238 | 52 | 4.60 | Yes |
|  | HA<sub>6</sub>-3APY | 49 | 91 | 0.54 | Yes |
|  | HA<sub>6</sub>-4AI | 77 | 28 | 2.75 | No |
|  | HA<sub>6</sub>-5AI | 162 | 133 | 1.20 | No |
|  | HA<sub>8</sub><sup>AN</sup> | 100 | 100 | 1.00 | Yes |
| modified<br>HA<sub>8</sub><sup>AN</sup> | HA<sub>8</sub>-2AA | 16 | 47 | 0.34 | Yes |
|  | HA<sub>8</sub>-3AA | 107 | 13 | 8.20 | Yes |
|  | HA<sub>8</sub>-4AA | 42 | 11 | 3.80 | Yes |
|  | HA<sub>8</sub>-2A4MBA | 21 | 141 | 0.15 | Yes |
|  | HA<sub>8</sub>-3APY | 115 | 23 | 5.00 | Yes |
|  | HA<sub>8</sub>-4AI | 141 | 37 | 3.80 | Yes |
|  | HA<sub>8</sub>-5AI | 207 | 96 | 2.20 | Yes |

<sup>a</sup> Not determined – affinity too low to be reliably measured by ITC. <sup>b</sup> Relative change in binding affinity compared to unmodified oligosaccharide of the same length as measured by ITC. <sup>c</sup> Relative change in binding affinity compared to unmodified oligosaccharide of the same length as measured by MST. <sup>d</sup> Substrate activity of modified HA oligosaccharide determined from *in vitro* HC-transfer assays.

### The effect of chemical modifications of HA oligosaccharides on TSG-6-mediated HC transfer

It is well established that HA can become covalently modified with HCs from the IaI family of proteoglycans in a biochemical process catalyzed by TSG-6 (25). Previous work has shown that the shortest HA^AN^ oligosaccharide that can act as a substate in HC transfer is an octasaccharide, where neither 4-mers nor 6-mers of HA become covalently attached to HC (39, 66). In this regard, HA_4_^AN^ and HA_6_^AN^ (unlike HA_8_^AN^) do not act as competitive inhibitors of HC transfer onto polymeric HA, and thus cannot inhibit processes such as cumulus expansion that requires the formation of high molecular weight HC•HA complexes (66). To begin to investigate the utility of our modified oligosaccharides as tool compounds, we assessed their activity as substrates for HC-transfer using an *in vitro* HC-transfer assay (26). Reactions were performed using purified IaI (from human plasma) and recombinant human TSG-6 (rhTSG-6) with the reaction products analyzed by SDS-PAGE (not shown) and Western blotting using an anti-HC1 antibody. Consistent with previous findings (39), while HA_8_^AN^ (and HA_14_^AN^, which we use as a positive control) acted as substrates, HA_4_^AN^ and HA_6_^AN^ did not (**Figure 11B, C**). We found that none of the chemical modifications described here substantially affected the ability of HA_8_^AN^ to act as a substrate, although HA_8_-4AA did appear to be a somewhat weaker substrate compared to HA_8_^AN^ (**Figure 11D, F**).

**Figure 11:**
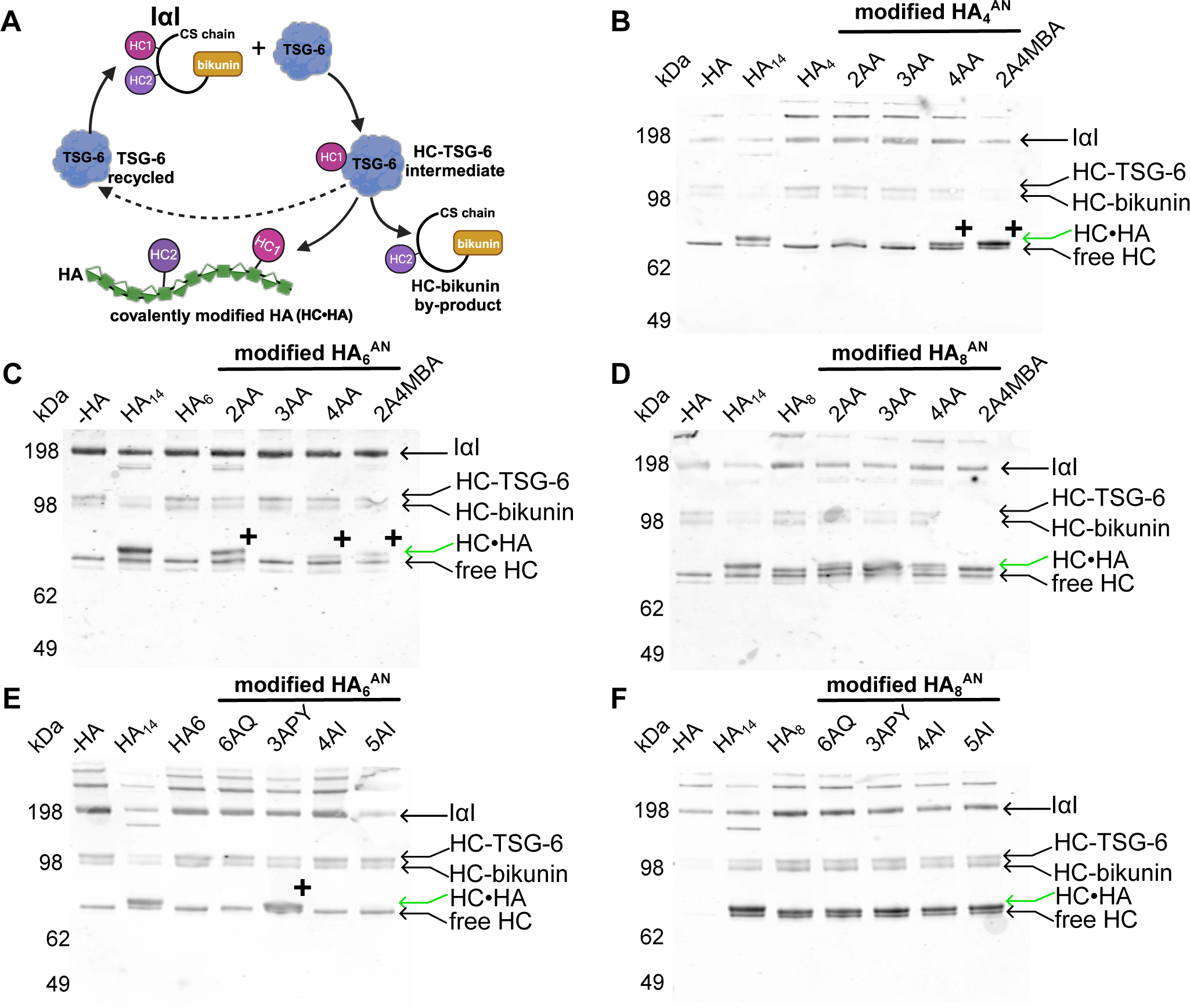
Modified HA_n_^AN^ oligosaccharides as substrates for HC-transfer. **A**) Schematic of TSG-6-mediated transfer of HA from IαI onto HA to form covalent HC•HA complexes; adapted from (25) and created using BioRender. TSG-6 reacts with IaI to form a covalent intermediate (HC•TSG-6) with either HC1 or HC2 (illustrated for HC1) along with a HC•bikunin by-product. The HC is then transferred from HC•TSG-6 onto HA (forming an ester bond with the C6 OH of a GlcNAc residue); this releases TSG-6, which can then react with another IaI molecule. (**B-E**) Western blots showing the results of HC-transfer reactions performed with 15 µM HA (modified HA oligosaccharides and unmodified controls), 1.8 µM IαI and 2.85 µM rhTSG-6 incubated for 2 hours at 37 °C: **B**) the 1st series of modified HA_4_^AN^; **C**) 1st series of modified HA_6_^AN^; (**D**) the 1st series of modified HA_8_^AN^; **E**) the 2nd series of modified HA_6_^AN^; **F**) the 2nd series of modified HA_8_^AN^. HA_14_^AN^ was used as a positive control throughout and unmodified HA_4_^AN^, HA_6_^AN^, and HA_8_^AN^ were used as controls in (**B**), (**C, E**) and (**D**, **F**), respectively. All blots were probed with anti-HC1 anti-sera (CP7, (98)). The running positions of IaI, HC•TSG-6, HC•bikunin and free HCs are shown to the right-hand side of the blots. HC•HA complexes run above the free HC, where ‘+’ indicates modified HA_4_^AN^ and HA_6_^AN^ oligomers that act as substrates for HC transfer, whereas the corresponding unmodified tetra- and hexasaccharides do not. Blots are representative of three independent experiments for each oligosaccharide.

Significantly, we found that several modifications changed the unreactive 6-mer (HA_6_^AN^) into a substrate for HC transfer. HA_6_-2AA (**Figure 11C**) and HA_6_-3APY (**Figure 11E**) both acted as good substrates for transfer, as indicated by the appearance of a band with an apparent molecular weight of ∼85 kDa (labelled with + on **Figure 11B, C**). Fainter bands were also seen with HA_6_-4AA and HA_6_-2A4MBA (**Figure 11C**) indicating these modifications modify the hexasaccharide to become weak substrates. No HC•HA_6_ complexes were observed when HA_6_^AN^ was modified with 3AA, 6AQ, 4AI or 5AI (**Figure 11E**), indicating that only the presence of certain chemical groups (on its reducing termini) leads to substrate activity. Remarkably, of the modified tetrasaccharides available, HA_4_-4AA and HA_4_-2A4MBA, were both able to act as good substrates (**Figure 11B**). These data clearly indicate that chemical modification of HA can alter its biochemical properties, and that some of the novel reagents made here may have potential as research tools in the investigation of HA biology.

## Discussion

Here we have described the design, synthesis, and characterization of the binding of 20 modified HA oligosaccharides (HA_4_^AN^-HA_8_^AN^) to two HABPs. We have identified several modifications that resulted in enhanced affinity of the oligosaccharide for either Link_TSG6 or CD44_HABD (compared to the unmodified oligosaccharide of the same length). Additionally, we have identified modifications that widen the discrimination between Link_TSG6 and CD44_HABD, as well as modified HA oligomers that become substrates for TSG-6-mediated HC transfer.

For CD44, we found that the largest gains in affinity were achieved with tetrasaccharides modified with aminobenzoic acid; HA_4_-2AA, HA_4_-3AA, and HA_4_-4AA all had increased affinity compared to HA_4_^AN^ alone. HA_4_-4AA had the highest affinity of those tested with a 4.5-fold increase compared to HA_4_^AN^ (HA_4_-4AA *K*_D_= 488 µM vs. HA_4_^AN^ *K*_D_ = 2200 µM), although this was still weaker than with HA_6_^AN^ or HA_8_^AN^ (**Table 2**). This is consistent with previous findings that these longer oligosaccharides are necessary for high-affinity binding to CD44, which occurs through the full complement of interactions within the HABD (31, 32). Interestingly, for HA_6_^AN^ and HA_8_^AN^ we found that most modifications to these lengths of oligosaccharide reduced the affinity of the interaction. Only HA_8_-2A4MBA and HA_6_-5AI had modest improvements in affinity (1.4-fold and 1.3-fold, respectively; **Table 5**). However, consistent with the CD44 HA-binding site containing many hydrophobic residues (31, 67), we found that the 2^nd^ series of modifications (3APY, 4AI and 5AI) resulted in smaller perturbations for HA_6_^AN^ and HA_8_^AN^ compared to 2AA, 3AA and 4AA. The increases in affinity seen here for the aminobenzoic acid-modified tetrasaccharides were similar to the findings of Lu and Huang (67), whose lead compound was a modified thiol glycoside attached to the closed reducing terminal ring of HA_4_^AN^. *In silico* modeling (67) suggested that the m-benzyl phenyl moiety of the compound could fit into the hydrophobic pocket (consisting of Cys81, Tyr83, Ile92 and Cys101), within the HA binding groove of CD44 (31, 67), π-π stacking against the tyrosine sidechain. Thus, the aromatic rings of the aminobenzoic acid moieties studied here could be involved in similar stacking interactions within this pocket, resulting in the improved affinities observed (see **Figure 1**). Our data are also consistent with recent work (48) demonstrating that polymeric HA (10 kDa) modified with multiple aromatic amines generated the best inhibitor of binding of unmodified HA to CD44 (although direct comparison of these IC_50_ values with the *K*_D_ values from our study is not possible). The (modest) gain in affinity achieved with modified HA_4_^AN^, reported here, supports the strategy of using HA oligosaccharide as scaffolds to enhance the binding to target HABPs. However, the affinity achieved with HA_4_-4AA (*K*_D_ = 488 µM) was very similar to that of a small-molecule ‘fragment’ identified by screening against CD44 (2-[(4-methyl1H-imidazol-5-yl)methyl]-1,2,3,4-tetrahydroisoquinolin-8-amine), which had *K*_D_ values of 0.4 mM and 0.5 mM for binding to the human and mouse CD44-HABDs, respectively (68). Interestingly, this compound does not occupy the HA-binding groove, but rather binds adjacent and orthogonal to it (68), and likely competes for binding via stabilizing the protein in its low affinity conformation (31).

In our study we targeted improving the affinity of HA binding to Link_TSG6, rather than to CD44, so further gains in affinity should be achievable for the latter, *i.e.*, despite CD44 often being considered ‘undruggable’ (48, 68–70). For example, based on our analysis of the structures for CD44_HABD in complex with a tetrasaccharide (67) or octasaccharide (31), modifying the non-reducing termini of GlcA residues in HA_4_^AN^ and HA_6_^AN^ with an appropriate acidic moiety should allow salt bridges to be formed with Arg82 and Arg155, respectively. Given that CD44-HA binding involves no ionic interactions (31), these modifications would be anticipated to lead to significant increases in affinity. Certainly, modified oligosaccharides with enhanced binding to CD44 would be expected to function as highly effective competitive inhibitors of polymeric HA-CD44 binding, compared to unmodified oligomers, due to the superselectivity of the CD44HA interactions (11, 50). As such the development of further tool compounds with greater specificity for CD44 appears warranted.

In addition to generating modified HA oligosaccharides that showed enhanced binding to CD44, we have also identified different modified oligomers that have increased affinity for Link_TSG6 (the isolated HABD of TSG-6). While modified HA_4_^AN^ were not studied in our ITC experiments with Link_TSG6 (since the large amount of material required makes them ‘incompatible’ with calorimetry experiments), we found that, unlike CD44, some modifications to HA_6_^AN^ and HA_8_^AN^ could improve the affinity for Link_TSG6 (*e.g*., 5AI modification of HA_6_^AN^ and HA_8_^AN^ led to increases of *K*_D_ by 1.6- and 2.1-fold, respectively); see **Table 1** and **Table 4**. This is not surprising given that we targeted improving the interaction with Link_TSG6 based on its different mode of binding compared to CD44 (39); see discussion below. Modification of HA_6_^AN^ with 2A4MBA resulted in the greatest increase in binding affinity for Link_TSG6 (*K*_D_ = 0.46 µM; 2.4-fold increase compared to HA_6_^AN^). The increased affinity for HA_6_-2A4MBA is likely due to the formation of an additional salt bridge between the carboxylate of the 2A4MBA moiety and the side chain of Arg81, consistent with the higher enthalpy of the interaction (compared to the unmodified oligomer; **Table 4**), as was predicted from the docking of the solution structure derived model for 2AA-modified HA into the Link_TSG6 structure; *i.e*., with HA_6_-2A4MBA forming a stronger ionic interaction than HA_6_-2AA, because the -OCH_3_ acts as an electron withdrawing group on the 2-aminobenzoic acid ring. This validates both the strategy we employed to enhance the HA-Link_TSG6 interaction, as well as the interaction network and the model of the Link_TSG6/HA_8_ complex proposed in Higman *et al*. (39).

Importantly, we found that HA_6_-3AA and HA_8_-3AA were the best compounds for discriminating binding to Link_TSG6 over CD44_HABD^20-169^ (with 19.8-fold and 8.2-fold relative differences in affinity, respectively), *i.e.,* compared to the unmodified HA_n_^AN^ oligomers (**Table 6**). On the other hand, HA_8_-2A4MBA was the modified oligosaccharide that could best discriminate binding of an octasaccharide to CD44 over Link_TSG6 (6.7-fold relative improvement); this modification reduced binding of HA_8_^AN^ to Link_TSG6 (4.8-fold) and increased the affinity to the CD44_HABD (1.4-fold). The ability to differentially alter the binding of HA to these two proteins is consistent with the different modes of binding proposed for CD44 and TSG-6 (31, 39, 42), underpinned by the differences in amino acid sequence position (and residue type) utilized within their Link module domains (46, 53). It is possible that further discrimination between CD44_HABD and Link_TSG6 could be achieved through design of oligosaccharides modified at both the reducing and non-reducing termini, *i.e*., utilizing HA oligosaccharides with either GlcA or GlcNAc as the non-reducing sugar (39, 45, 46). In this regard, Higman *et al*. (39) identified that HA_7_^AA^ or HA_7_^NN^ had only slightly reduced binding affinities for Link_TSG6 compared to HA_8_^AN^ (72% and 68%, respectively), whereas HA_8_^NA^ was found to have a 1.9-fold greater affinity, suggestive that these oligomers may provide good starting points for future investigation.

It may be possible to extend our approach to other members of the Link module family. In this regard, like TSG-6 (41, 42, 71), ionic interactions are known to contribute to HA binding for other HABPs, *e.g*., HAPLN1 (72, 73) and LYVE-1 (74), whereas aggrecan and neurocan, like CD44, exhibit salt-strength independent interactions (75, 76). In the case of Link_TSG6, ionic bonds only contribute to ∼25% of the free energy of binding to HA (71) illustrating the importance of the contribution from non-basic amino acids via hydrogen bonds and CH-π ring stacking interactions. The 3 tyrosine residues in TSG-6 that contribute to HA binding (27, 40, 41) are also highly conserved in many (*e.g*., HAPLN1-4, the 4 lecticans (aggrecan, brevican, neurocan and versican) and stabilin-2/HARE) but not all (*e.g*., LYVE-1) Link module containing proteins (42). One major difference between the HAPLNs/lecticans and TSG-6 is that the former proteins all have an HA-binding domain composed of 2 contiguous Link modules that both contribute to HA binding (see (77, 78)). This ‘type-C’ HABD (20, 22, 79) requires a minimum of 10 saccharides for an optimal interaction with HA (77), and as such this length difference compared to CD44 and TSG-6 could potentially be exploited to make tool compounds that are selective for this class of HABPs.

In this study, we determined the solution dynamic 3D structure of HA_4_-2AA and found the modified GlcNAc-2AA portion librates around 5 distinct conformers. Applying this behavior to a model of HA6-2AA, an ensemble of 250 representative conformations were docked into the Link_TSG6 structure (in its HA-bound conformation (41)); see **Figure 10**. The best scoring conformation (*i.e*., the best fit into the HA-binding groove) was from Conformer 2, which is one of the most highly populated conformers in solution (see **Figure 8**). This is consistent with the finding that the common conformers of ligands observed in solution are often very similar to their bioactive conformations (55, 56, 80). Importantly, HA is highly conformationally dynamic (81–83), such that different HAPBs likely capture different HA conformations that are frequently visited in solution, leading to distinct architectures of HA/protein complexes when multiple protein molecules interact with the same HA chain, *i.e.,* through propagating local structural effects to the polymer level (1, 21).

As described in Higman *et al.*, HA_7_ oligosaccharides (both HA_7_^AN^ and HA_7_^NN^) were the shortest HA oligosaccharides that act as substrates for HC-transfer (39). In the present study, we have found that some chemically modified HA_6_^AN^ and HA_4_^AN^ oligosaccharides could become covalently coupled to an HC (*e.g*., HA_4_-AA, HA_4_-2A4MBA, HA_6_-2AA, HA_6_3AA, HA_6_-2A4MBA and HA_6_-3APY); see **Figure 11**. This is interesting, and surprising, given the previous findings regarding the minimum length of oligosaccharide required to accept HCs from TSG-6 (39, 66). Here, consistent with our previous data on unmodified HA oligomers (39), we have found that there is no correlation between the affinity of an oligosaccharide for Link_TSG6 and its substrate potential in TSG-6-mediated HC-transfer assays (using full-length TSG6); see **Table 6**. For example, certain modified HA_8_^AN^ oligosaccharides (*e.g*., HA_8_-2AA) had reduced affinity for Link_TSG6 (6.25-fold) but retained full activity as substrates in the transfer reaction. As noted previously (27), while the Link module of TSG-6 contributes to HA recognition in the context of the covalent HC•TSG-6 intermediate, it is unlikely that the entire HA-binding groove is involved in this process; based on the finding that of the 3 tyrosine’s involved in HA binding in ‘free’ TSG-6 (*i.e*., Tyr12, Tyr59 and Tyr78, which are numbered Tyr47, Tyr94 and Tyr113, respectively, in the full-length protein), only the mutation of Tyr47 to phenylalanine impairs TSG-6-mediated HC transfer onto HA. Moreover, the interaction of IaI with TSG-6 and the formation of the HC•TSG-6 intermediate inhibits the binding of HA to TSG-6 (36). Thus, the covalently attached HC likely partially occludes the HA-binding groove in TSG-6 and forms part of a composite interaction site that only involves a few residues of the Link module (27, 39) including Tyr47. Consistent with this, we have found here, that HA_4_-4AA and HA_4_-2A4MBA act as good substrates for HC transfer, where these oligosaccharides only have 3 intact sugar rings in addition to the chemically modified open ring. Overall, this indicates that the HA recognition site in HC•TSG-6 is smaller than has been anticipated and, also, is unlikely to utilize the binding mechanism we have identified here for the interaction of the modified sugars with Link_TSG6 given that the Tyr78/Arg81- end of the binding groove does not appear to be involved in HC transfer (27).

Our results may further suggest that the conformation of the modified HA oligosaccharide is an important determinant as to whether the oligosaccharide can accept the HC during transfer. It is known that HCs are transferred from the hydroxyls of GalNAc in the CS chain of IαI (where they are linked via ester bonds (84, 85) to Ser-28 of TSG-6 (27), and then transferred to the C6 hydroxyls of a GlcNAc in HA (86, 87), *i.e.*, via two sequential TSG-6-mediated transesterification reactions (26). Thus, it seems likely that the modified HA oligosaccharides that act as substrates can be accommodated in the composite binding site in the HC•TSG-6 complex in such a way that promotes transfer of the HC onto the C6 hydroxyl group of a suitably positioned GlcNAc residue. From the solution dynamic 3D structure of HA_4_2AA determined here, we observed that the 6C hydroxyl group of the reducing terminal (open) GlcNAc ring is in a somewhat different position compared to that in the unmodified HA_n_^AN^ (**Figure 9**). However, we do not know if it is this or the other GlcNAc (labelled S2 in **Figure 8**) that accepts the ester bond on HC transfer. Furthermore, it is not clear why HA_4_-2AA does not act as a substrate for HC transfer, whereas HA_4_-4AA and HA_4_-2A4MBA do, and whether differences in the orientations of C6 OH groups in these modified oligomers contribute to their different activities. This seems likely given that the carboxylate group of 2AA contributes to the conformational behavior of HA_4_-2AA via forming a hydrogen bond to the aniline proton (X5 HN11), where such an interaction is not possible in HA_4_-4AA (because of the different position of the COOH moiety on the benzene ring). Thus, a number of different solution conformations may bind to the HC•TSG-6 complex where only some of these are correctly oriented so as to be able to accept the ester bond connecting the HC to Ser28. Additional investigations are clearly required to understand the molecular mechanism of HC Transfer. From the work described here, it may be possible to further develop the chemically modified HA oligosaccharides that act as substrates to allow labeling of the amino acid residues in the vicinity of the active site. Understanding this mechanism is of medical importance given the role of HC•HA matrix formation in various physiological and pathological processes ranging from ovulation to lung disease (4–6, 17, 27, 30, 88).

Several of the chemical groups selected in this study (*e.g*., 2AA, 3AA and 6AQ) are already popular as derivatization reagents for carbohydrates (64, 89–93), owing to their fluorescent properties (53, 64). This, combined with the findings described here for the altered binding of chemically modified HA oligosaccharides to CD44 and TSG-6, highlights the utility of such modifications in generating reagents to probe HA biology. Our work provides a foundation for the further development of tool compounds, and perhaps specific inhibitors, for the study and therapeutic modulation of HA-protein interactions in complex tissue environments.

### Experimental procedures

#### Production of recombinant proteins

Link_TSG6 and ^15^N-labeled Link_TSG6 were expressed and purified as described previously (44, 94). A novel construct consisting of residues 20-169 of human CD44 and an N-terminal six-histidine tag (denoted hisCD44_HABD^20-169^), was expressed in *E. coli*, refolded, and purified using a method adapted from Banerji *et al.* (95) for the related CD44_HABD^20178^ construct. Briefly, refolding of hisCD44_HABD^20-169^ solubilized from inclusion bodies was carried out by rapid dilution (50-fold) into 50 mM Tris-HCl, pH 8.0, 1 mM EDTA, 0.5 M L-arginine, 1 mM L-cysteine and 0.2 mM L-cystine, followed by overnight incubation at 4 °C. Protein was then purified using a combination of nickel-affinity chromatography (IMAC) and size exclusion chromatography (SEC), followed by high-performance liquid chromatography (HPLC) on a Jupiter C5 300 Å column (Phenomenex). Where ^15^N-labelled hisCD44_HABD^20-169^ or Link_TSG6 were required for NMR spectroscopy, *E. coli* were grown in M9 minimal media (0.6% (w/v) Na_2_HPO_4_, 0.3% (w/v) KH_2_PO_4_, 0.5% (w/v) NaCl, 0.1 mM CaCl_2_, 1 mM MgSO_4_, 0.052% (w/v) ^15^N-NH_4_Cl, 0.2% (w/v) glucose, 0.085% (w/v) DIFCO yeast nitrogen base without NH_4_Cl) and purified as for the wild-type proteins. Samples of purified proteins were analyzed by electrospray ionization mass spectrometry (ESI-MS), showing they were <1 Da from the theoretical molecular mass for the unlabeled proteins and with 99.6% and 99.9% isotope incorporation for ^15^N-labeled hisCD44_HABD^20-169^ and Link _TSG6, respectively; 1D ^1^H and [^1^H-^15^N]-HSQC NMR spectroscopy (0.1 mM protein, pH 7.5, 10% (v/v) D_2_O, 25 °C, 0.3 mM DSS-d_6_ reference, Bruker Avance 500 MHz spectrometer) showed they were correctly folded.

#### Production and chemical modification of HA oligosaccharides

Hyaluronan oligosaccharides with an even number of monosaccharides and reducing terminal GlcNAc residues (termed HA_n_^AN^) were produced by digestion of 10 mg/mL sodium hyaluronate (HASK-1; Lifecore Biomedical) with 1 kU/mL ovine testicular hyaluronidase (Calbiochem) in 150 mM NaCl, 100 mM NaAc, pH 5.2 for 20 hours (45, 46). A 20-hour digestion at 37 °C resulted in a mixture of oligosaccharide lengths, predominantly HA_4_^AN^, HA_6_^AN^ and HA_8_^AN^, with some HA_10_^AN^ (45). To produce modified oligosaccharides, reductive amination reactions were performed based on a method adapted from Ruhaak *et al.* (93). The reagents, all sourced from Sigma Aldrich, are detailed in **Table S1**. Oligosaccharides (in the digestion mixture) were combined with a reactant amine compound at 500 mM in DMSO, 15% (v/v) acetic acid and then combined with 1 M 2-picolene borane (in a final ratio of 2:1:1 (v/v/v), oligosaccharide: amine: reducing agent) before incubation at 65 °C for 2 hours in a heat block. Reactions were terminated by addition of 5 volumes of deionized water immediately followed by SEC on a XK16/20 column packed in-house with BioGel P2 beads (BioRad). The column was equilibrated in 50 mM CH_3_COONH_4_, pH 7.0, running at 1 mL/min. Modified and unmodified oligosaccharides of different lengths were separated using anion exchange chromatography (AEX) on a HiPrep XK 16/10 column (GE Healthcare) at a flow rate of 5 mL/min using a gradient of 0 – 150 mM NaCl over 140 mins. AEX fractions were collected, lyophilized, reconstituted in 1 mL of deionized H_2_O, and desalted on a BioGel P-2 column (as described in (45)). Fractions of 3 mL were collected (for each species) and lyophilized after reconstitution in 20 mL of H_2_O, five times, to reduce residual CH_3_COONH_4_. Purified oligosaccharides were analyzed by matrix-assisted laser desorption ionization time-of-flight (MALDI-TOF) mass spectrometry (using a matrix of 10 mg/mL 6-aza-2-thiothymine in a 1:1 (v/v) solution of acetonitrile and 20 mM ammonium citrate). Modified oligosaccharides used in further analyses were within <1.5 Da of the expected molecular weight. 1D-^1^H NMR spectroscopy was used to provide further data as evidence of correct modifications. None of the 2^nd^ series of modified tetrasaccharides (HA_4_^AN^ with 3APY, 4AI or 5AI) had the expected molecular weights (data not shown) so these were excluded from further analyses.

#### Measuring HA-protein interactions

##### Link_TSG6 – Isothermal Titration Calorimetry

ITC experiments were performed using a MicroCal PEAQ-ITC instrument (Malvern Instruments). Oligosaccharide stock solutions were prepared at 0.29 mM (in 5 mM MES, pH 6.0) and injected (18 x 2 µL with 120 s spacing between injections) into the cell containing ∼200 µL Link_TSG6 (0.029 mM) in matched buffer at 25 °C. Data were fit to a one site binding model, after subtracting the heats from the last three injections to account for heat of dilution of the oligosaccharide into buffer. Each oligosaccharide was analyzed in four independently performed ITC experiments.

##### NMR spectroscopy

[^1^H-^15^N]-HSQC spectra were recorded for samples of ^15^N-labelled Link_TSG6 (0.1 mM, pH 6.0, 10% (v/v) D_2_O, 90% (v/v) H2O, 25 °C, 0.3 mM DSS-d_6_ reference) on a Bruker Avance 500 MHz spectrometer. After an initial reference spectrum for ^15^N-labelled Link_TSG6 was collected, oligosaccharide (either HA_6_^AN^, HA_8_^AN^, HA_6_-2AA, or HA_8_-2AA) was added incrementally up to a 10-fold molar excess. Backbone amide resonances were assigned by comparison to those of both free (PDB accession code 1o7b; Biological Magnetic Resonance Databank (BMRB) accession code: 6392) and HA_8_^AN^-bound (PDB: 1o7c; BMRB: 6393) Link_TSG6 reported previously (41).

##### CD44_HABD – Microscale Thermophoresis

For hisCD44_HABD^20-169^, interactions with HA oligosaccharides were assessed using MST. Protein (200 nM) was labelled with 5 µM red-Tris-NTA-650 dye following the manufacturer’s instructions in 10 mM HEPES, pH 7.4, 150 mM NaCl, 0.05% (v/v) Tween-20 (interaction buffer). Stock solutions of HA oligosaccharides (5 mM in interaction buffer) were serially diluted (1:1 in interaction buffer) 16 times before each concentration was mixed with an equal volume of labelled protein (10 nM final concentration hisCD44_HABD^20-169^). Samples were loaded into standard glass capillaries (NanoTemper) and thermophoresis measured at 25 °C (with active temperature control) using a Monolith NT.115pico instrument (NanoTemper) running NT Control software (version 1.0.1). All experiments were performed in triplicate and data analyzed with the ‘*K*_D_ fit model’ in the NT Analysis software (version 2.0.2).

##### Analysis of HA_4_-2AA and determination of its solution dynamic 3D structure by NMR spectroscopy

1D and 2D homonuclear and heteronuclear NMR experiments were performed on samples of HA_4_-2AA to allow resonance assignment and determination of its solution dynamic 3D structure. All samples were analyzed on a Bruker Avance 500 MHz spectrometer fitted with a QXI (quadrupole) cryoprobe or a Bruker Avance-DRX 800 MHz spectrometer with a TCI cryoprobe.

A pH titration was performed using [^1^H]-1D spectra recorded on a single sample of 1 mM HA_4_-2AA, 5% (v/v) D_2_O, 95% (v/v) H_2_O at 5 °C, adjusting the pH value between 2.0 and 11.0. A temperature series of [^1^H]-1D spectra was recorded on a single sample of 1 mM HA_4_-2AA, 5% (v/v) D_2_O, 95% (v/v) H_2_O, pH 9.4 at 5, 15, 25 and 37 °C. Other spectra used for assignment and measurement of structural restraints ([1H]-1D, [^1^H-^1^H]-COSY, [^1^H-^1^H]-TOCSY, [^1^H^-13^C]-HSQC, [^1^H^13^C]-HSQC-TOCSY, [^1^H-^1^H]-NOESY (mixing time 700 ms)) were recorded at natural abundance using standard parameters (55, 58) at 5 °C on samples of 1 mM HA_4_-2AA, 5% (v/v) D_2_O, 95% (v/v) H_2_O, pH 9.4 or 3 mM HA_4_-2AA, 100% (v/v) D_2_O, pH 6.0.

The solution dynamic 3D structure of the reducing terminus of HA_4_-2AA (penultimate GlcA residue, modified reducing terminal GlcNAc residue, and the aminobenzoic acid moiety) was determined according to the methodology described in (56). Three kinds of structural restraints were used in the structure calculations: NOEs and noNOEs were measured from a [^1^H-^1^H]-NOESY spectrum (700 ms mixing time) recorded on a sample of 3 mM HA_4_-2AA, 100% (v/v) D_2_O, pH 6.0. ^3^*J*_HH_ conformation-dependent scalar couplings were measured directly from a [^1^H]-1D spectrum (1 mM HA_4_-2AA, 5% (v/v) D_2_O, pH 9.4), and the temperature coefficient (Δδ_H_/ΔT) of the aniline proton was calculated by linear regression of its chemical shift with temperature (1 mM HA_4_-2AA, 5% (v/v) D_2_O, pH 9.4). For the rest of the structure, the unmodified portion of HA_4_-2AA was given the dynamic 3D structure parameters of HA_6_^AN^ determined previously (60) consistent with the identity of chemical shifts and NOE cross-peaks for HA oligosaccharides (see Results). Torsions involving hydroxyls and carboxylate groups (which in water cannot be measured with NMR data) were given values based on the dynamic 3D structure used for HA_6_^AN^ (60), expectations from mining the Cambridge Structural Database (CSD; (96)) and consideration of their positions within each determined conformer.

##### Generation of a structural model for the HA_6_-2AA/Link_TSG6 complex

The conformational parameters from the solution dynamic 3D structure of HA_4_-2AA were used to model appropriate conformations for rigid-body docking into the NMR structure of Link_TSG6 in its HA_8_^AN^-bound conformation (PDB accession code 1o7c; (41)). An ensemble of 250 conformations of HA_6_-2AA was generated by taking the (fixed) conformation for the 4 non-reducing terminal residues from the Link_TSG6-bound structure of HA_8_^AN^ (as reported in (39)) and sampling conformations for the reducing terminal (opened) GlcNAc and 2AA residues according to the measured solution conformational behavior of HA_4_-2AA. The GOLD software program (CSD release 2019.3, (62, 96)) was then used to dock each of the 250 conformations of HA_6_-2AA into the NMR structure of Link_TSG6 in its HA_8_^AN^bound conformation (PDB accession code 1o7c; (41)). The starting pose for the 5 non-reducing terminal monosaccharides was maintained as reported in (39) such that the modeling investigated how the reducing terminal modified GlcNAc and 2AA residues could be accommodated in the HA-binding groove of Link_TSG6; both protein and ligand conformations were kept rigid throughout the docking. Ensemble member docked poses were analyzed using the GoldScore metric scoring function to score each member in terms of its hydrogen bonding energy, van der Waals energy, and ligand torsion strain. All poses were written into structure-data format (SDF) files and visually inspected using Maestro (V12.4) and PyMol (v2.4, Schrӧdinger, USA) software.

##### HC•HA transfer assays

Covalent transfer of HC from IαI onto unmodified and chemically modified HA oligosaccharides was analyzed using the assay described previously (26). All reactions were performed using a buffer of 20 mM HEPES, pH 7.5, 150 mM NaCl, 5 mM MgCl_2_. In a ‘standard’ assay, HA oligosaccharides were at 15 μM, human IaI (97)1.8 at μM (based on a molecular weight of 180 kDa), and recombinant human TSG-6 (rhTSG-6; R&D Systems) at 2.8 μM (based on a molecular weight of 35 kDa). Reaction mixtures were analyzed by SDS-PAGE using 4-12% bis-tris polyacrylamide gels (Invitrogen). For Western blot analysis, proteins were transferred onto nitrocellulose membrane (GE Healthcare) and incubated with antisera against HC1 (CP7, 1:10,000; (98)). Bound primary antibody was detected using goat-anti-rabbit IgG conjugated to IR dye (LI-COR Biosciences) and blots imaged using an Odyssey Clx near-infrared fluorescence imaging system (LICOR Biosciences).

## Abbreviations used are

2AA: 2-aminobenzoic acid;
2A4MBA: 2-amino-4-methoxybenzoic acid;
3AA: 3aminobenzoic acid;
3APY: 3-aminopyrazole;
4AA: 4-aminobenzoic acid;
4AI: 4-aminoindan;
5AI: 5aminoindan;
HA_n_^AN^: (n = number of saccharides in HA oligomer) with non-reducing terminal GlcA (A) and reducing terminal GlcNAc (N);
HABD: Hyaluronan-binding domain;
HABP: Hyaluronan-binding protein;
TSG6: secreted product of Tumor Necrosis Factor-stimulated Gene-6;
IαI: Inter-a-Inhibitor;
ITC: Isothermal Titration Calorimetry;
MST: Microscale Thermophoresis;
NMR: Nuclear Magnetic Resonance spectroscopy.

## Supporting Information

This article contains supporting information.

## Acknowledgements

We are very grateful to Wojtek Augustinyak for technical support with NMR experiments and Thorsten Nowak for thoughtful advice on chemical modification strategies (both at C4X Discovery). We thank Reynard Spiess (Mass Spectrometry Facility, Manchester Institute of Biotechnology) for MALDI-TOF analysis of oligosaccharides and Emma Jayne Keevil (Biological Mass Spectrometry Facility) for intact-mass spectrometry analysis of proteins. We also thank Matthew Cliff (Biomolecular NMR Facility, Manchester Institute of Biotechnology) for his assistance with NMR experiments and training. All biomolecular analyses were carried out in the Biomolecular Analysis Core Facility (University of Manchester), with helpful advice from Tom Jowitt, which is supported by Centre funding from the Wellcome Trust (203128/Z/16/Z and 220926/Z/20/Z). This work was funded by a BBSRC CASE DTP (1792538), to AJD and CMM, sponsored by C4X Discovery, and a BBSRC SYNBIOCHEM FTMA2 Innovation Placement award (to RJD and AJD).

## Author Contributions

**Conceptualization**: A.J.D, C.D.B, R.J.D.; **Formal analysis**: R.J.D, C.D.B, B.M.S.; **Investigation**: R.J.D, C.D.B, B.M.S.; **Resources**: A.J.D., J.J.E.; **Writing original draft**: R.J.D.; **Writing review & editing**: R.J.D, A.J.D, C.D.B., B.M.S., J.J.E., C.C.M; **Supervision**: A.J.D., C.D.B, C.M.M.; **Funding acquisition**: A.J.D, C.D.B, C.M.M.

