## Supplemental Information for "Chemical modification of hyaluronan oligosaccharides differentially modulates hyaluronan- hyaladherin interactions"

**Supporting Information Contents**


| | | Page |
|---|---|---|
| ***A. Characterization of modified HA oligosaccharides.*** | | |
| Table S1 | Amine-containing reagents used to chemically modify $HA_n^{AN}$ oligosaccharides. | S-3 |
| Table S2 | Molecular masses for unmodified and modified HA oligosaccharides. | S-3 |
| Figure S1 | Example MALDI-TOF mass spectra of modified HA. | S-4 |
| ***B. Investigation of the interaction of HA-2AA modified oligosaccharides with Link_TSG6 by NMR spectroscopy.*** | | |
| Figure S2 | [$^1$H-$^{15}$N]-HSQC spectra of Link_TSG6 in the presence of unmodified and modified HA hexasaccharides. | S-5 |
| Figure S3 | [$^1$H-$^{15}$N]-HSQC spectra of Link_TSG6 in the presence of unmodified and modified HA octasaccharides. | S-5 |
| ***C. Purification and characterization of the new CD44_HABD construct.*** | | |
| Figure S4 | Purification and characterization of hisCD44_HABD$^{20\text{-}169}$. | S-6 |
| ***D. Determination of the solution dynamic 3D structure of $HA_4$-2AA.*** | | |
| Table S3 | $^1$H chemical shifts for $HA_4$-2AA. | S-7 |
| Table S4 | $^{13}$C chemical shifts for $HA_4$-2AA. | S-8 |
| Table S5 | Temperature coefficients ($^1$H $\Delta\delta_{HN}/\Delta T$) of exchangeable amide and aniline hydrogen nuclei in $HA_4$-2AA compared to $HA_4^{AN}$. | S-9 |
| Table S6 | Conformation-dependent scalar coupling constants for residue T4 of $HA_4$-2AA (modified GlcNAc) compared to β-configuration GlcNAc rings in $HA_4^{AN}$. | S-9 |
| Table S7 | Conformational restraints derived from scalar couplings and their $\chi^2$ fits. | S-9 |
| Table S8 | NOE conformational restraints for $HA_4$-2AA and their $\chi^2$ fits. | S-10 |
| Table S9 | noNOE conformational restraints for residues G3-X5 in $HA_4$-2AA and their $\chi^2$ fits. | S-11 |
| Table S10 | Summary of $\chi^2$ fits for all structural restraints for residues G3-X5 in $HA_4$-2AA. | S-12 |
| Table S11 | Torsion angle conformational preferences for $HA_4$-2AA in aqueous solution. | S-13 |
| Figure S5 | 2D structure and nomenclature of $HA_4$-2AA. | S-14 |
| Figure S6 | pH titration of $HA_4$-2AA. | S-14 |
| Figure S7 | Chemical shift perturbations in $HA_4$-2AA compared to $HA_4^{AN}$. | S-15 |
| Figure S8 | Solution dynamic 3D structure of $HA_4$-2AA in a conformational ensemble representation. | S-16 |
| ***E. Generation of a model of the $HA_6$-2AA/Link_TSG6 complex.*** | | |
| Figure S9 | Comparison of conformations of $HA_8^{AN}$ bound to Link_TSG6 with the modeled conformations of $HA_6$-2AA used for docking into the binding site of Link_TSG6. | S-17 |
| ***F. Supporting Information References.*** | | S-18 |

***A. Characterization of modified HA oligosaccharides.***

**Table S1. Amine-containing reagents used to chemically modify $HA_n^{AN}$ oligosaccharides.**

| Chemical | Abbreviated name | Modified HA oligosaccharide |
|---|---|---|
| 2-aminobenzoic acid | 2AA | $HA_n$•2AA |
| 3-aminobenzoic acid | 3AA | $HA_n$•3AA |
| 4-aminobenzoic acid | 4AA | $HA_n$•3AA |
| 2-amino-4-methoxy benzoic acid | 2A4MBA | $HA_n$•2A4MBA |
| 3-amino-pyrazole | 3APY | $HA_n$•3APY |
| 4-aminoindan | 4AI | $HA_n$•4AI |
| 5-aminoindan | 5AI | $HA_n$•5AI |
| 6-amino-quinolone | 6AQ | $HA_n$•6AQ |

**Table S2. Molecular masses for unmodified and modified HA oligosaccharides.**

| HA | Formula | $[M-H]^-$ (Da) [a] | Observed mass (Da) [b] | Difference (Da) [c] |
|---|---|---|---|---|
| **$HA_4^{AN}$** | $C_{28}H_{44}N_2O_{23}$ | 775.6 | 775.7 ±0.1 | 0.1 |
| **$HA_4$-2AA** | $C_{35}H_{49}N_3O_{24}$ | 895.6 | 894.3 ± 0.1 | 1.3 |
| **$HA_4$-3AA** | $C_{35}H_{49}N_3O_{24}$ | 895.6 | 896.5 ± 0.2 | 0.9 |
| **$HA_4$-4AA** | $C_{35}H_{49}N_3O_{24}$ | 895.6 | 896.9 ± 0.7 | 1.3 |
| **$HA_4$-2A4MBA** | $C_{36}H_{52}N_3O_{25}$ | 927.3 | 926.1 ± 0.3 | 1.2 |
| **$HA_6^{AN}$** | $C_{42}H_{65}N_3O_{34}$ | 1154.9 | 1155.1 ±0.2 | 0.2 |
| **$HA_6$-2AA** | $C_{49}H_{70}N_4O_{35}$ | 1275.5 | 1275.6 ± 0.1 | 0.1 |
| **$HA_6$-3AA** | $C_{49}H_{70}N_4O_{35}$ | 1275.5 | 1275.3 ± 0.1 | 0.2 |
| **$HA_6$-4AA** | $C_{49}H_{70}N_4O_{35}$ | 1275.5 | 1274.8 ± 0.2 | 0.7 |
| **$HA_6$-2A4MBA** | $C_{50}H_{73}N_4O_{36}$ | 1306.6 | 1305.5 ± 0.7 | 1.1 |
| **$HA_6$-3APY** | $C_{45}H_{69}N_6O_{33}$ | 1221.1 | 1221.5 ± 0.1 | 0.4 |
| **$HA_6$-4AI** | $C_{51}H_{75}N_4O_{33}$ | 1271.1 | 1271.5 ± 0.1 | 0.4 |
| **$HA_6$-5AI** | $C_{51}H_{75}N_4O_{33}$ | 1271.1 | 1271.7 ± 0.1 | 0.6 |
| **$HA_6$-6AQ** | $C_{51}H_{72}N_5O_{33}$ | 1282.1 | 1282.3 ± 0.0 | 0.2 |
| **$HA_8^{AN}$** | $C_{56}H_{86}N_4O_{45}$ | 1534.3 | 1534.5 ±0.2 | 0.2 |
| **$HA_8$-2AA** | $C_{63}H_{91}N_5O_{46}$ | 1654.7 | 1654.9 ± 0.1 | 0.2 |
| **$HA_8$-3AA** | $C_{63}H_{91}N_5O_{46}$ | 1654.7 | 1654.6 ± 0.6 | 0.1 |
| **$HA_8$-4AA** | $C_{63}H_{91}N_5O_{48}$ | 1654.7 | 1654.9 ± 0.1 | 0.2 |
| **$HA_8$-2A4MBA** | $C_{64}H_{94}N_5O_{47}$ | 1686.0 | 1687.3 ± 0.4 | 1.3 |
| **$HA_8$-3APY** | $C_{59}H_{90}N_7O_{44}$ | 1600.4 | 1600.4 ± 0.1 | 0.0 |
| **$HA_8$-4AI** | $C_{65}H_{96}N_5O_{44}$ | 1650.5 | 1651.1 ± 0.4 | 0.6 |
| **$HA_8$-5AI** | $C_{65}H_{96}N_5O_{44}$ | 1650.5 | 1651.1 ± 0.1 | 0.6 |
| **$HA_8$-6AQ** | $C_{65}H_{94}N_6O_{44}$ | 1661.5 | 1661.7 ± 0.5 | 0.2 |

[a] Theoretical value of $[M-H]^-$ based on the average molecular mass of oligosaccharide. [b] Determined from negative-ion MALDI-TOF mass spectra; value is the average mass recorded on three individual sample spots ± standard deviation. [c] Difference between the theoretical mass and the observed mass.

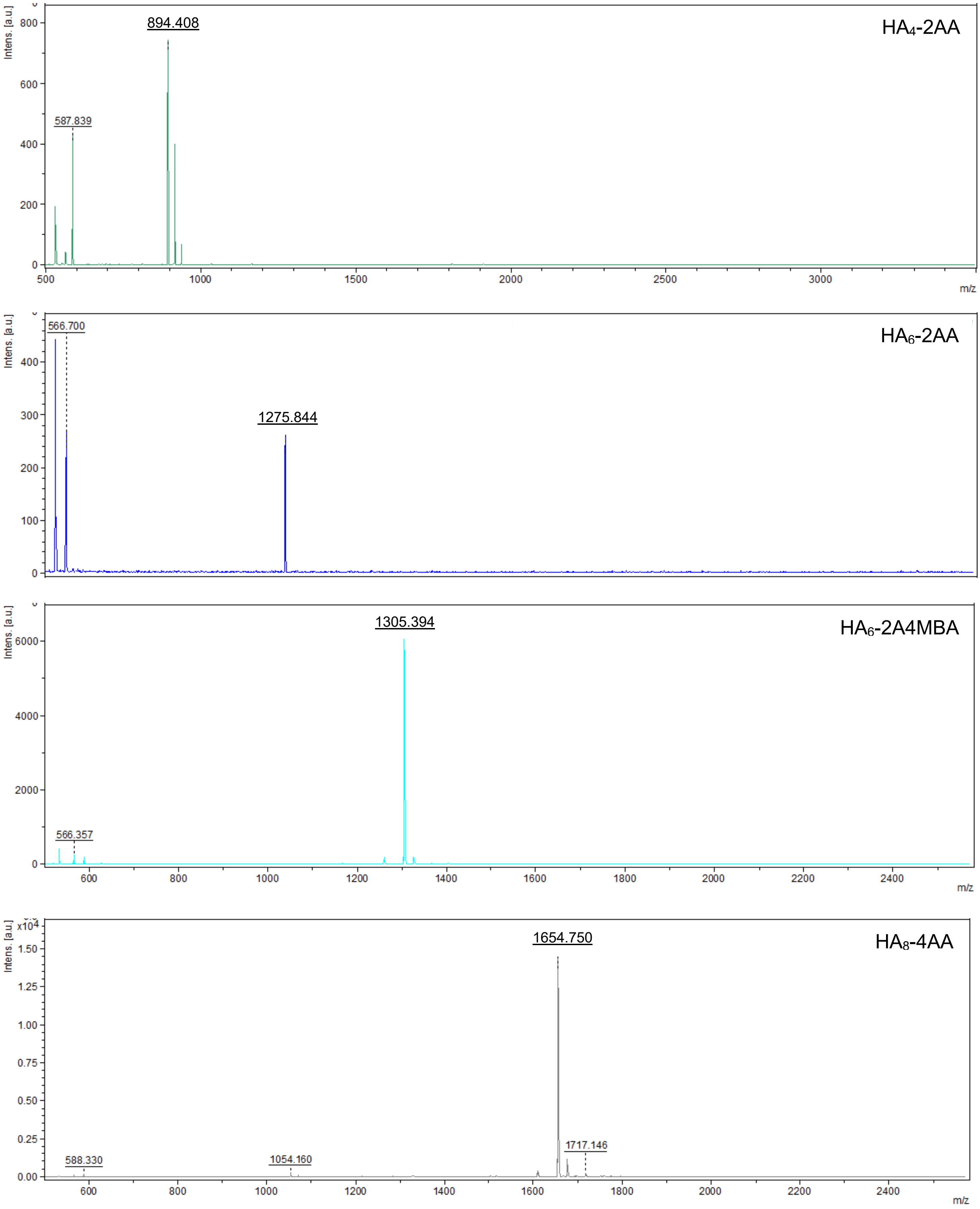


**Figure S1. Example MALDI-TOF mass spectra of modified HA**. Spectra are representative of those recorded on three individual samples. Additional species (500 Da) can be attributed to the MALDI analysis matrix (6-aza-2-thiothymine).

***B. Investigation of the interaction of HA-2AA modified oligosaccharides with Link_TSG6 by NMR spectroscopy.***

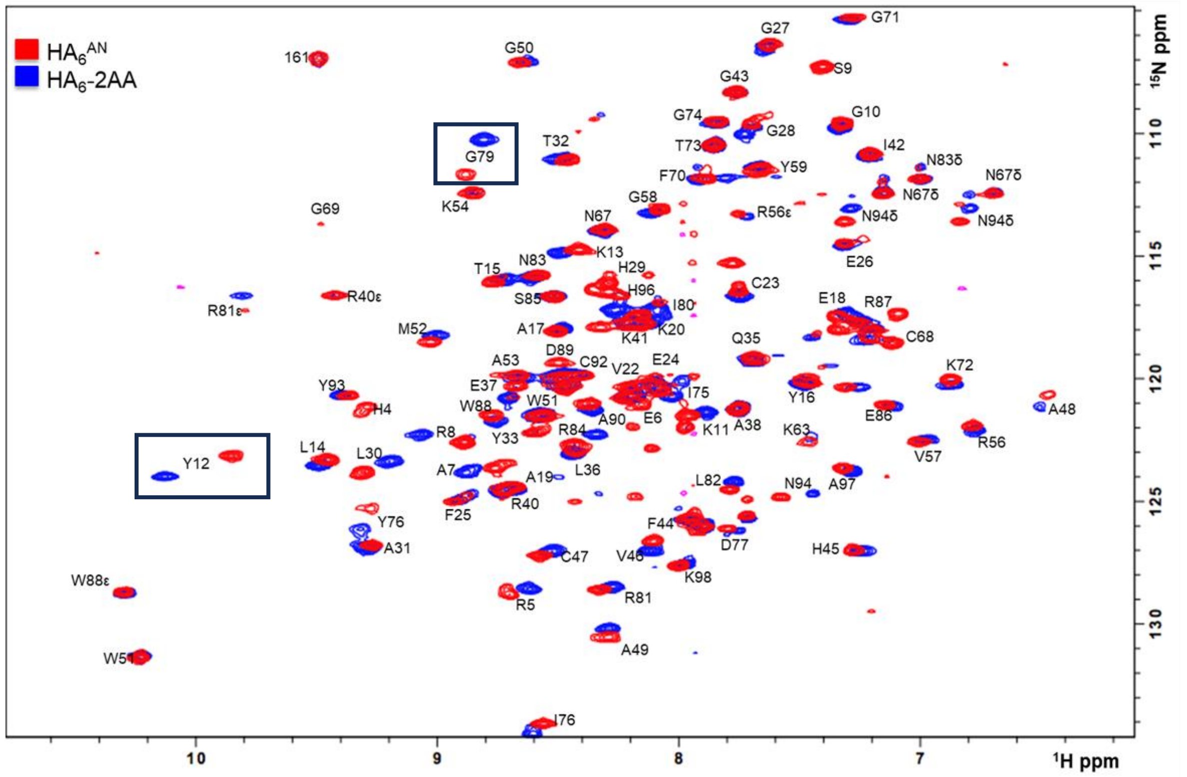


**Figure S2. [$^1$H-$^{15}$N]-HSQC spectra of Link_TSG6 in the presence of unmodified and modified HA hexasaccharides.** [$^1$H-$^{15}$N]-HSQC NMR spectra of 0.1 mM $^{15}$N-Link_TSG6 in the presence of 10-fold molar excess of either $HA_6^{AN}$ (red) or $HA_6$-2AA (blue). NH resonances are labelled in one letter code (backbone and sidechain) according to the assignment for Link_TSG6 reported previously (PDB accession code: 1o7c, (1)). Chemical shift perturbations of Tyr12 (Y12) and Gly79 (G79) are highlighted (square boxes).

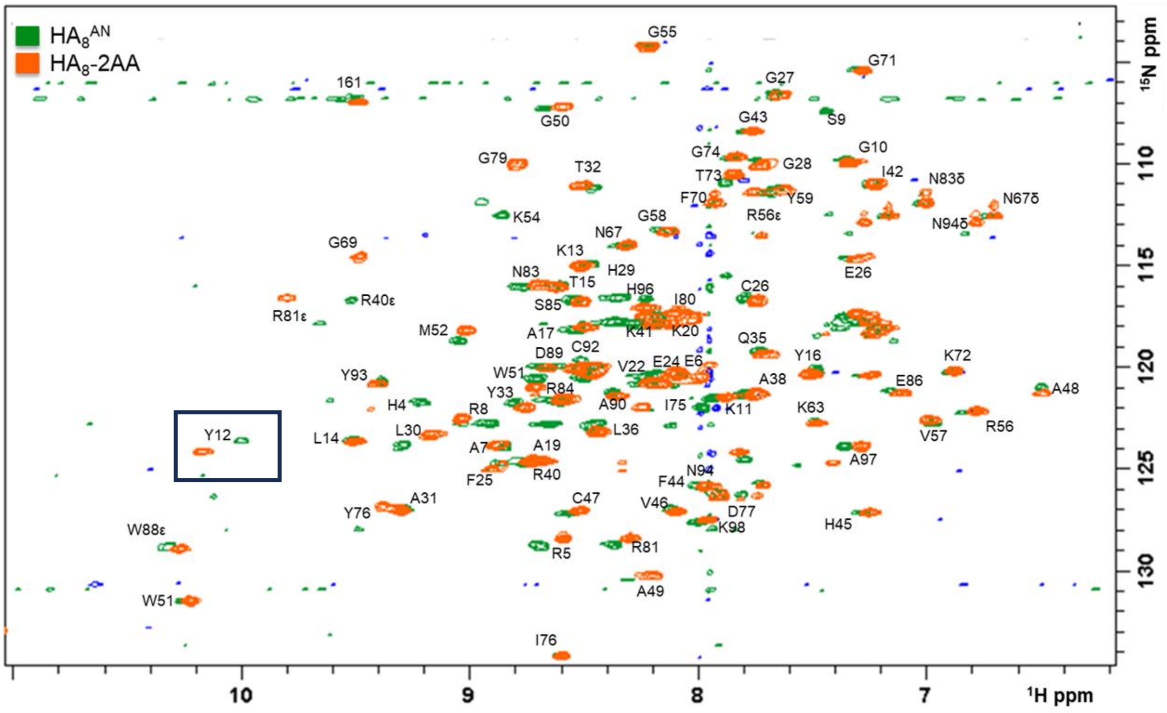


**Figure S3. [$^1$H-$^{15}$N]-HSQC spectra of Link_TSG6 in the presence of unmodified and modified HA octasaccharides.** [$^1$H-$^{15}$N]-HSQC NMR spectra of 0.1 mM $^{15}$N-Link_TSG6 in the presence of 10-fold molar excess of either $HA_8^{AN}$ (green) or $HA_8$-2AA (orange). NH resonances are labelled in one letter code (backbone and sidechain) according to the assignment for Link_TSG6 reported previously (pdb accession code: 1o7c (1)). Chemical shift perturbation of Tyr12 (Y12) is highlighted (square box).

*C. Purification and characterisation the new construct for the HA-binding domain of CD44*

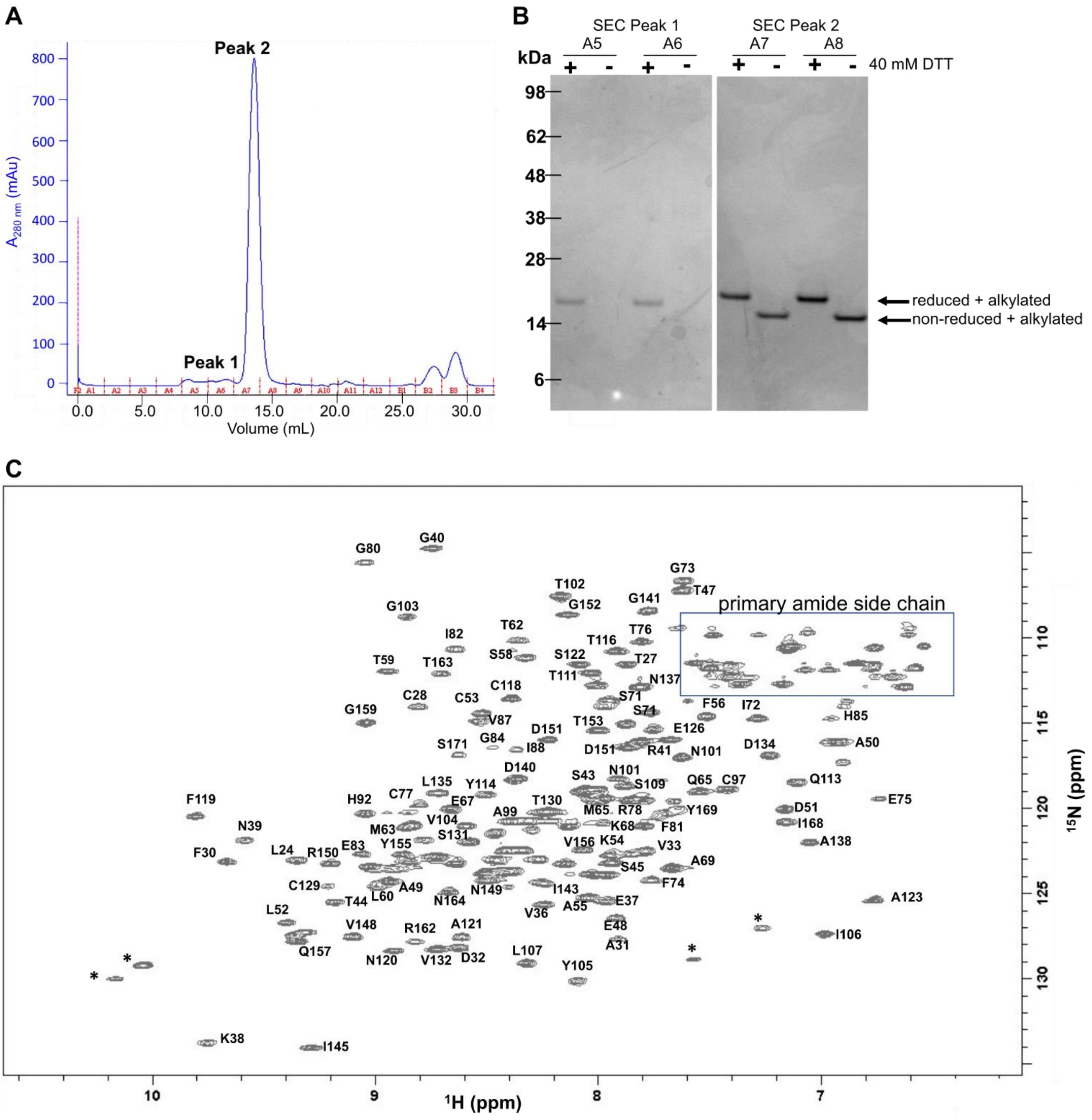


**Figure S4. Purification and characterization of hisCD44_HABD$^{20\text{-}169}$. A**) Size Exclusion Chromatography (SEC) of the hisCD44_HABD$^{20\text{-}169}$ construct following its expression *in E. coli*, refolding and initial purification on IMAC (not shown); the eluent was monitored continuously at A280$_{nm}$ and 2.5-mL fractions were collected. **B**) Fractions across Peak 1 (A5, A6) and Peak 2 (A7, A8) from SEC in (**A**) were analyzed under reduced (+ 40 mM DTT and alkylated) or non-reduced (alkylated) conditions. The majority of the hisCD44_HABD$^{20\text{-}169}$ protein was present in Peak 2, where this had a different apparent molecular weight on SDS-PAGE under reducing and non-reducing conditions (~20 and ~15 kDa, respectively), consistent with the presence of disulfide bonds; a band was only seen under reduced conditions for Peak 1 indicating this corresponds to aggregated material. **C**) 0.1 mM $^{15}$N-hisCD44_HABD$^{20\text{-}169}$ was analyzed by [$^{1}$H-$^{15}$N]-HSQC NMR at pH 7.5 on a spectrometer operating at 500 MHz. The spectrum was essentially identical to that for CD44_HABD$^{20\text{-}178}$ (2), allowing for its assignment, except that it lacks the resonances from the absent 9 C-terminal residues, which considerably improves the resolution at the crowded center. The primary amide side chain region for Asn (N) and Qlu (Q) residues is indicated (box).

*D. Determination of the solution dynamic 3D structure of $HA_4$-2AA.*

**Table S3. $^1H$ chemical shifts for $HA_4$-2AA.**

| Residue | Atom | **$^1H$ δ (ppm)** [a] **$HA_4^{AN}$** pH 6.0, 24.4 °C 10% (v/v) $D_2O$ 5-20 mM | **$^1H$ δ (ppm)** **$HA_4$-2AA** pH 9.4, 5.0 °C 10% (v/v) $D_2O$, 1 mM | **$^1H$ δ (ppm)** **$HA_4$-2AA** pH 6.0, 5 °C 100% (v/v) $D_2O$, 3 mM | **Δδ (ppm)** [b] **$HA_4^{AN}$** pH 6.0 vs **$HA_4$-2AA** pH 9.4 | **Δδ (ppm)** [c] **$HA_4$-2AA** pH 6.0 vs **$HA_4$-2AA** pH 9.4 |
|---|---|---|---|---|---|---|
| **G1 GlcA** | H1 [d] | 4.458 [e] | 4.468 | 4.468 | -0.010 | 0.000 |
| | H2 | 3.321 | 3.323 | 3.317 | -0.002 | 0.006 |
| | H3 | 3.496 | 3.486 | 3.486 | -0.010 | 0.000 |
| | H4 | 3.497 | 3.485 | 3.485 | 0.012 | 0.000 |
| | H5 | 3.723 | 3.725 | 3.725 | -0.059 | 0.000 |
| **S2 GlcNAc** | H1 | 4.560 | 4.561 | 4.561 | -0.001 | 0.000 |
| | H2 | 3.851 | 3.858 | 3.858 | -0.007 | 0.000 |
| | H3 | 3.707 | 3.705 | 3.705 | 0.002 | 0.000 |
| | H4 | 3.542 | 3.554 | 3.554 | -0.012 | 0.000 |
| | H5 | 3.484 | 3.477 | 3.477 | 0.007 | 0.000 |
| | H61 | 3.921 | 3.924 | 3.918 | -0.003 | 0.006 |
| | H62 | 3.780 | 3.792 | 3.792 | -0.012 | 0.000 |
| | HNA1 | - [f] | 8.202 | 8.198 | - | 0.004 |
| | HA3* | 2.028 | 2.034 | 2.024 | -0.006 | 0.010 |
| **G3 GlcA** | H1 | 4.467 | 4.578 | 4.578 | -0.111 | 0.000 |
| | H2 | 3.363 | 3.411 | 3.404 | -0.048 | 0.007 |
| | H3 | 3.580 | 3.620 | 3.620 | -0.040 | 0.000 |
| | H4 | 3.744 | 3.728 | 3.728 | 0.016 | 0.000 |
| | H5 | 3.703 | 3.728 | 3.728 | -0.025 | 0.000 |
| **T4 GlcNAc** | H11 | - [g] | 3.646 | 3.646 | - | 0.000 |
| | H12 | - [g] | 3.247 | 3.244 | - | 0.003 |
| | H2 | 3.806 | 4.441 | 4.441 | -0.635 | 0.000 |
| | H3 | 3.716 | 4.065 | 4.062 | -0.349 | 0.003 |
| | H4 | 3.514 | 3.567 | 3.567 | -0.053 | 0.000 |
| | H5 | 3.472 | 3.846 | 3.846 | -0.374 | 0.000 |
| | H61 | 3.750 | 3.831 | 3.824 | -0.081 | 0.007 |
| | H62 | 3.890 | 3.606 | 3.606 | 0.284 | 0.000 |
| | HNA1 | - [f] | 8.243 | 8.237 | - | 0.006 |
| | HA3* | 2.015 | 1.933 | 1.933 | 0.082 | 0.000 |
| **X5 2AA** | HN11 | - [g] | 7.965 | - [h] | - | - |
| | H2 | - [g] | 6.914 | 6.919 | - | -0.005 |
| | H3 | - [g] | 7.760 | 7.760 | - | 0.000 |
| | H4 | - [g] | 6.771 | 6.776 | - | -0.005 |
| | H5 | - [g] | 7.374 | 7.374 | - | 0.000 |

[a] Chemical shifts for the β-anomer of $HA_4^{AN}$ as reported in (3). [b] Chemical shift changes from $HA_4^{AN}$ (as reported in (3)) including temperature and pH difference. [c] Refer to Figure S5 for nomenclature. [d] Chemical shift changes resulting from replacement of $H_2O$ with $D_2O$ and increase in pH. [e] All chemical shifts were referenced relative to internal DSS-$d_6$. Standard error on measurements: $^1H$ ± 0.001 ppm. [f] Resonance not visible due to exchange with $D_2O$. [g] Equivalent atom not present in $HA_4^{AN}$. [h] Resonance not visible at pH 6.0.

**Table S4. $^{13}$C chemical shifts for $HA_4$-2AA.**

| Residue | Atom | $^{13}$C δ (ppm) $HA_4^{AN}$ [a]<br>pH 6.0, 24.4 °C<br>10% (v/v) $D_2O$<br>5-20 mM | $^{13}$C δ (ppm) $HA_4$-2AA<br>pH 9.4, 5.0 °C<br>10% (v/v) $D_2O$<br>1 mM | Δδ (ppm) [b]<br>$HA_4^{AN}$ pH 6.0, 24.4 °C<br>vs.<br>$HA_4$-2AA pH 9.4, 5.0 °C |
|---|---|---|---|---|
| **G1 GlcA** | C1 [c] | 105.715 [d] | 105.707 | 0.008 |
| | C2 | 75.578 | 75.364 | 0.214 |
| | C3 | 78.213 | 77.971 | 0.242 |
| | C4 | 74.560 | 74.443 | 0.117 |
| | C5 | 78.477 | 78.614 | -0.137 |
| **S2 GlcNAc** | C1 | 103.325 | 103.827 | -0.502 |
| | C2 | 57.049 | 56.990 | 0.059 |
| | C3 | 85.839 | 85.623 | 0.216 |
| | C4 | 71.357 | 71.100 | 0.257 |
| | C5 | 78.183 | 77.963 | 0.220 |
| | C6 | 63.390 | 63.094 | 0.296 |
| | CA3 | 25.326 | 25.212 | 0.114 |
| **G3 GlcA** | C1 | 105.818 | 105.608 | 0.072 |
| | C2 | 75.294 | 75.345 | -0.051 |
| | C3 | 76.506 | 76.159 | 0.347 |
| | C4 | 82.781 | 82.551 | 0.230 |
| | C5 | 79.098 | 78.586 | -0.512 |
| **T4 GlcNAc** | C1 | 97.879 | 46.087 | 51.502 |
| | C2 | 58.454 | 53.001 | 5.453 |
| | C3 | 85.151 | 79.482 | 5.669 |
| | C4 | 71.351 | 73.363 | -2.012 |
| | C5 | 78.264 | 73.217 | 5.047 |
| | C6 | 63.567 | 65.347 | -1.780 |
| | CA3 | 25.065 | 24.764 | 0.301 |
| **X5 2AA** | C2 | - [e] | 115.966 | - |
| | C3 | - [e] | 134.273 | - |
| | C4 | - [e] | 119.769 | - |
| | C5 | - [e] | 135.268 | - |

[a] Chemical shifts for the β-anomer of $HA_4^{AN}$ as reported in (3). [b] Chemical shift changes from $HA_4^{AN}$ including temperature and pH changes. [c] Refer to Figure S5 for nomenclature. [d] All chemical shifts were referenced indirectly relative to internal DSS-$d_6$. Standard error on measurements: $^{13}$C ± 0.020 ppm. [e] Atom not present in $HA_4^{AN}$.

**Table S5. Temperature coefficients ($\Delta\delta_H/\Delta T$) of exchangeable amide and aniline hydrogen nuclei in $HA_4$-2AA compared to $HA_4^{AN}$.**

| Residue | Atom | **$HA_4$-2AA** pH 6.0, 3 mM 100% (v/v) $D_2O$ ppb/K | **$HA_4$-2AA** pH 9.4, 1 mM 10% (v/v) $D_2O$ ppb/K | **$HA_4^{AN}$, pH 6.0** [a] pH 6.0, 5-20 mM 10% (v/v) $D_2O$ ppb/K |
|---|---|---|---|---|
| **S2** | HNA1 [b] | -6.8 [c] | -6.9 | -7.0 |
| **T4** | HNA1 | -6.8 | -6.8 | -[d] |
| **X5** | HN11 | -[e] | -2.6 | -[d] |
| $H_2O$ [f] | | -11.3 | -11.3 | -[d] |

[a] Temperature coefficient as reported in (3). [b] Refer to Figure S5 for nomenclature. [c] Standard error on measurements: ± 0.1 ppb/K. [d] Equivalent atom not present in $HA_4^{AN}$. [e] HN was no longer observed due to fast chemical exchange with solvent at this pH value. [f] Temperature coefficient of water reported as internal control.

**Table S6. Conformation-dependent scalar coupling constants for residue T4 of $HA_4$-2AA (modified GlcNAc) compared to β-configuration GlcNAc rings in $HA_4^{AN}$.**

| $^3J_{HH}$ coupling constant | $HA_4$-2AA [a] T4-GlcNAc residue (Hz) | $HA_4^{AN}$ [b] β-GlcNAc (Hz) | internal GlcNAc (Hz) | Mean $\Delta^3J_{HH}$ [c] (Hz) |
|---|---|---|---|---|
| H2-HNA1/HN [d] | 9.8 [e] | 9.70 | 9.68 | 0.1 |
| H2-H3 | 6.1 | - | 10.37 | 4.3 |
| H3-H4 | 1.7 | 8.81 | - | 7.2 |
| H4-H5 | 8.5 | 9.73 | 10.00 | 1.4 |
| H5-H61 | -[f] | 5.47 | 5.32 | - |
| H5-H62 | 4.1 | 2.20 | 2.31 | -1.7 |

[a] Values measured at pH 9.4, 5 °C, 1 mM in 10% (v/v) $D_2O$. [b] Values for β-GlcNAc and internal (I)-GlcNAc rings in $HA_4^{AN}$ as reported in (3)).[c] Mean difference between values in residue T4 (of $HA_4$-2AA) compared to the same coupling constant in either the β-GlcNAc or internal GlcNAc ring of $HA_4^{AN}$. [d] Refer to Figure S5 for nomenclature. [e] All coupling constants shifts were determined by direct measurement from 1D spectra. Standard error on all couplings ± 0.05 Hz. [f] Value not measured due to spectral overlap and lineshape distortion caused by strong coupling.

**Table S7. Conformational restraints derived from scalar couplings and their $\chi^2$ fits.**

| **$^3J_{HH}$ coupling constant** | **Karplus parameters** [a] **A** | **B** | **C** | **phase Ψ** | **observed value (Hz)** | **calculation error applied (Hz)** [c] | **mean predicted value (Hz)** | **mean χ2** |
|---|---|---|---|---|---|---|---|---|
| T4 H11-H2 | 8.804 | -0.990 | 1.452 | 1 | 3.7 ± 0.05 | ± 0.75 | 3.22 | 0.497 |
| T4 H12-H2 | 9.036 | -0.983 | 1.336 | -7 | 11.0 ± 0.05 | ± 0.75 | 1.05 | 0.489 |
| T4HNA1-H2 | 9.450 | -2.080 | 0.630 | -5 | 9.8 ± 0.05 | ± 0.75 | 9.99 | 0.155 |
| T4 H2-H3 | 7.564 | -0.896 | 0.847 | 10 | 6.1 ± 0.05 | ± 0.75 | 5.58 | 0.068 |
| T4 H3-H4[b] | 7.516 | -0.879 | 0.567 | -15 | 1.7 ± 0.05 | ± 0.75 | 1.61 | 0.046 |
| T4 H4-H5 | 6.578 | -0.910 | 1.036 | 0 | 8.5 ± 0.05 | ± 0.75 | 8.35 | 0.099 |
| T4 H62-H5 | 8.317 | -0.990 | 1.371 | -1 | 4.1 ± 0.05 | ± 0.75 | 4.17 | 0.123 |
| X5 HN11-T4 H11 | 7.900 | -1.050 | 0.650 | -5 | 6.6 ± 0.05 | ± 0.75 | 6.25 | 0.369 |
| X5 HN11-T4 H12 | 7.900 | -1.050 | 0.650 | -5 | 4.5 ± 0.05 | ± 0.75 | 4.59 | 0.415 |

[a] Karplus equation in the form: $^3J = A\cos^2(\theta+\Psi) + B\text{os}(\theta+\Psi) + C$, with Ψ in degrees (4). [b] Refer to Figure S5 for nomenclature. [c] Error applied to observed $^3J_{HH}$ value is that of predictive capability of the Karplus relation, rather than measurement error (5).

**Table S8. NOE conformational restraints for $HA_4$-2AA and their $\chi^2$ fits.**

| donor residue [a, b] | donor nuclei | acceptor residue | acceptor nuclei | measured height [c] | calculation error applied | mean predicted height | mean χ2 |
|---|---|---|---|---|---|---|---|
| G3 | H1 | G3 | H1 | 9100 | 3640 | 5690 | 0.885 |
| G3 | H3 | G3 | H1 | 1090 | 435 | 1120 | 0.018 |
| G3 | H4 & H5 [d] | G3 | H1 | 2290 | 916 | 1940 | 0.151 |
| T4 | H12 | G3 | H1 | 293 | 117 | 108 | 2.510 |
| T4 | H2 | G3 | H1 | 365 | 146 | 224 | 0.940 |
| T4 | H3 | G3 | H1 | 1860 | 742 | 1520 | 0.209 |
| T4 | H4 | G3 | H1 | 131 | 52 | 181 | 0.979 |
| G3 | H4 & H5 | G3 | H2 | 1640 | 656 | 1380 | 0.151 |
| T4 | H2 | G3 | H2 | 54 | 22 | 48 | 0.104 |
| T4 | H3 | G3 | H2 | 138 | 57 | 85 | 0.864 |
| T4 | H5 & H61 | G3 | H2 | 201 | 80 | 143 | 0.747 |
| T4 | HA3* | G3 | H2 | 28 | 12 | 25 | 0.194 |
| G3 | H1 | G3 | H3 | 1350 | 539 | 1120 | 0.196 |
| G3 | H3 | G3 | H3 | 10900 | 4360 | 11200 | 0.009 |
| G3 | H1 | T4 | H11 | 211 | 86 | 103 | 1.620 |
| T4 | H4 | T4 | H11 | 333 | 133 | 312 | 0.059 |
| X5 | H2 | T4 | H11 | 351 | 141 | 243 | 0.612 |
| G3 | H1 | T4 | H12 | 218 | 87 | 108 | 1.620 |
| T4 | H4 | T4 | H12 | 379 | 152 | 178 | 1.760 |
| X5 | H2 | T4 | H12 | 229 | 93 | 273 | 0.241 |
| G3 | H1 | T4 | H2 | 306 | 123 | 224 | 0.458 |
| T4 | H2 | T4 | H2 | 5820 | 2330 | 6650 | 0.167 |
| T4 | H4 | T4 | H2 | 370 | 148 | 443 | 0.261 |
| T4 | H5 & H61 | T4 | H2 | 123 | 49 | 98 | 0.268 |
| X5 | H2 | T4 | H2 | 339 | 136 | 308 | 0.138 |
| G3 | H1 | T4 | H3 | 2020 | 809 | 1520 | 0.387 |
| G3 | H2 | T4 | H3 | 89 | 36 | 85 | 0.019 |
| G3 | H4 & H5 | T4 | H3 | 495 | 198 | 361 | 0.457 |
| T4 | H5 & H61 | T4 | H3 | 430 | 172 | 326 | 0.384 |
| T4 | HA3 | T4 | H3 | 90 | 36 | 18 | 3.960 |
| X5 | H2 | T4 | H3 | 177 | 71 | 119 | 0.670 |
| T4 | H12 | T4 | H4 | 283 | 113 | 178 | 0.861 |
| T4 | H2 | T4 | H4 | 391 | 156 | 443 | 0.128 |
| X5 | H2 | T4 | H4 | 85 | 34 | 135 | 2.130 |
| G3 | H1 | X5 | H2 | 56 | 23 | 54 | 0.063 |
| T4 | H11 | X5 | H2 | 288 | 116 | 243 | 0.175 |
| T4 | H12 | X5 | H2 | 179 | 72 | 273 | 1.750 |
| T4 | H2 | X5 | H2 | 336 | 135 | 308 | 0.132 |
| T4 | H3 | X5 | H2 | 73 | 29 | 119 | 2.510 |
| X5 | H4 | X5 | H4 | 9780 | 3910 | 10500 | 0.041 |
| X5 | H5 | X5 | H5 | 7570 | 3040 | 10200 | 0.734 |

[a] The donor and acceptor nuclei are those on the indirect and direct dimensions of the spectrum, respectively. [b] Refer to Figure S5 for nomenclature. [c] Measured height, calculation error applied, predicted heights, and calculated mean $\chi^2$, as described in (5)). Data measured from a 2D-NOESY recorded with 700 ms mixing time on a 3 mM sample at pH 6.0 in 100% (v/v) $D_2O$ and at 5 °C. [d] These restraints were included as overlaps between the NOE cross-peaks indicated.

**Table S9. noNOE conformational restraints for residues G3-X5 in $HA_4$-2AA and their $\chi^2$ fits.**

| donor residue [a, b] | donor nuclei | acceptor residue | acceptor nuclei | measured height [c] | calculation error applied | mean predicted height | mean $\chi^2$ |
|---|---|---|---|---|---|---|---|
| X5 | H3 | G3 | H1 | 0.0 | 33.3 | 1.6 | 0.002 |
| X5 | H4 | G3 | H1 | 0.0 | 32.3 | 2.1 | 0.004 |
| X5 | H5 | G3 | H1 | 0.0 | 41.1 | 9.6 | 0.055 |
| T4 | H12 | G3 | H2 | 0.0 | 55.5 | 23.9 | 0.192 |
| X5 | H2 | G3 | H2 | 0.0 | 18.0 | 9.7 | 0.294 |
| X5 | H3 | G3 | H2 | 0.0 | 14.3 | 4.1 | 0.088 |
| X5 | H4 | G3 | H2 | 0.0 | 19.4 | 1.7 | 0.008 |
| X5 | H5 | G3 | H2 | 0.0 | 13.2 | 2.9 | 0.050 |
| T4 | H12 | G3 | H3 | 0.0 | 29.4 | 19.8 | 0.463 |
| T4 | H2 | G3 | H3 | 0.0 | 44.8 | 27.1 | 0.365 |
| T4 | HA3 | G3 | H3 | 0.0 | 30.1 | 18.9 | 0.404 |
| X5 | H2 | G3 | H3 | 0.0 | 30.0 | 10.5 | 0.125 |
| X5 | H3 | G3 | H3 | 0.0 | 18.4 | 1.13 | 0.005 |
| G3 | H2 | T4 | H11 | 0.0 | 22.9 | 14.9 | 0.428 |
| T4 | H5 | T4 | H11 | 0.0 | 36.2 | 40.3 | 1.240 |
| T4 | H61 | T4 | H11 | 0.0 | 36.7 | 16.3 | 0.202 |
| T4 | HA3 | T4 | H11 | 0.0 | 17.5 | 5.1 | 0.085 |
| X5 | H3 | T4 | H11 | 0.0 | 13.3 | 6.1 | 0.212 |
| X5 | H4 | T4 | H11 | 0.0 | 14.3 | 15.1 | 1.130 |
| G3 | H2 | T4 | H12 | 0.0 | 39.0 | 23.9 | 0.389 |
| T4 | H5 | T4 | H12 | 0.0 | 30.0 | 25.2 | 0.705 |
| T4 | H61 | T4 | H12 | 0.0 | 26.8 | 8.62 | 0.106 |
| T4 | HA3 | T4 | H12 | 0.0 | 20.4 | 6.53 | 0.103 |
| X5 | H3 | T4 | H12 | 0.0 | 19.8 | 6.25 | 0.101 |
| X5 | H4 | T4 | H12 | 0.0 | 23.1 | 14.2 | 0.385 |
| G3 | H3 | T4 | H2 | 0.0 | 47.5 | 27.1 | 0.325 |
| T4 | H62 | T4 | H2 | 0.0 | 63.4 | 17.5 | 0.077 |
| T4 | HA3 | T4 | H2 | 0.0 | 46.9 | 37.9 | 0.652 |
| X5 | H3 | T4 | H2 | 0.0 | 22.7 | 6.1 | 0.071 |
| X5 | H4 | T4 | H2 | 0.0 | 21.7 | 8.9 | 0.169 |
| X5 | H3 | T4 | H3 | 0.0 | 22.4 | 2.4 | 0.011 |
| X5 | H4 | T4 | H3 | 0.0 | 27.4 | 3.5 | 0.016 |
| X5 | H5 | T4 | H3 | 0.0 | 23.3 | 18.0 | 0.600 |
| G3 | H2 | T4 | H4 | 0.0 | 50.8 | 21.6 | 0.182 |
| T4 | HA3 | T4 | H4 | 0.0 | 19.9 | 12.0 | 0.367 |
| X5 | H3 | T4 | H4 | 0.0 | 14.3 | 2.1 | 0.028 |
| X5 | H4 | T4 | H4 | 0.0 | 13.8 | 3.5 | 0.066 |
| G3 | H2 | T4 | HA3 | 0.0 | 91.8 | 24.5 | 0.073 |
| G3 | H3 | T4 | HA3 | 0.0 | 54.8 | 18.9 | 0.122 |
| G3 | H4 | T4 | HA3 | 0.0 | 49.1 | 4.3 | 0.008 |
| G3 | H5 | T4 | HA3 | 0.0 | 50.5 | 8.1 | 0.026 |
| T4 | H11 | T4 | HA3 | 0.0 | 53.2 | 5.1 | 0.009 |
| T4 | H12 | T4 | HA3 | 0.0 | 120.0 | 6.5 | 0.003 |
| T4 | H2 | T4 | HA3 | 0.0 | 105.0 | 37.9 | 0.129 |
| T4 | H4 | T4 | HA3 | 0.0 | 52.0 | 12.0 | 0.054 |
| T4 | H5 | T4 | HA3 | 0.0 | 55.1 | 5.9 | 0.012 |
| T4 | H61 | T4 | HA3 | 0.0 | 50.4 | 1.9 | 0.001 |
| T4 | H62 | T4 | HA3 | 0.0 | 57.1 | 1.8 | 0.001 |
| X5 | H2 | T4 | HA3 | 0.0 | 237.0 | 6.0 | 0.001 |
| X5 | H5 | T4 | HA3 | 0.0 | 178.0 | 5.6 | 0.001 |
| G3 | H2 | X5 | H2 | 0.0 | 10.3 | 9.7 | 0.891 |
| G3 | H3 | X5 | H2 | 0.0 | 26.6 | 10.5 | 0.159 |
| G3 | H4 | X5 | H2 | 0.0 | 11.5 | 2.3 | 0.040 |
| G3 | H5 | X5 | H2 | 0.0 | 8.88 | 12.1 | 1.870 |
| T4 | H61 | X5 | H2 | 0.0 | 10.4 | 6.4 | 0.388 |
| T4 | H62 | X5 | H2 | 0.0 | 11.9 | 5.2 | 0.199 |
| T4 | HA3 | X5 | H2 | 0.0 | 14.6 | 6.0 | 0.170 |
| G3 | H1 | X5 | H3 | 0.0 | 12.0 | 1.6 | 0.017 |
| G3 | H2 | X5 | H3 | 0.0 | 10.2 | 4.1 | 0.175 |
| G3 | H3 | X5 | H3 | 0.0 | 12.1 | 1.1 | 0.010 |
| G3 | H4 | X5 | H3 | 0.0 | 13.3 | 0.6 | 0.002 |
| G3 | H5 | X5 | H3 | 0.0 | 13.3 | 0.4 | 0.001 |
| T4 | H11 | X5 | H3 | 0.0 | 12.8 | 6.1 | 0.227 |
| T4 | H12 | X5 | H3 | 0.0 | 11.9 | 6.3 | 0.280 |
| T4 | H2 | X5 | H3 | 0.0 | 14.4 | 6.1 | 0.177 |
| T4 | H3 | X5 | H3 | 0.0 | 13.0 | 2.4 | 0.033 |
| T4 | H4 | X5 | H3 | 0.0 | 12.1 | 2.1 | 0.029 |
| T4 | H5 | X5 | H3 | 0.0 | 13.3 | 0.6 | 0.002 |
| T4 | H61 | X5 | H3 | 0.0 | 13.3 | 0.2 | 0.000 |
| T4 | H62 | X5 | H3 | 0.0 | 12.1 | 0.2 | 0.000 |
| T4 | HA3 | X5 | H3 | 0.0 | 14.8 | 4.3 | 0.088 |

| | | | | | | | |
|---|---|---|---|---|---|---|---|
| G3 | H1 | X5 | H4 | 0.0 | 10.2 | 2.1 | 0.043 |
| G3 | H2 | X5 | H4 | 0.0 | 11.1 | 1.7 | 0.025 |
| G3 | H3 | X5 | H4 | 0.0 | 15.8 | 1.1 | 0.005 |
| G3 | H4 | X5 | H4 | 0.0 | 14.0 | 0.3 | 0.000 |
| G3 | H5 | X5 | H4 | 0.0 | 14.0 | 0.6 | 0.002 |
| T4 | H11 | X5 | H4 | 0.0 | 15.8 | 15.1 | 0.920 |
| T4 | H12 | X5 | H4 | 0.0 | 14.1 | 14.2 | 1.040 |
| T4 | H2 | X5 | H4 | 0.0 | 12.9 | 8.9 | 0.479 |
| T4 | H3 | X5 | H4 | 0.0 | 11.0 | 3.5 | 0.100 |
| T4 | H4 | X5 | H4 | 0.0 | 15.8 | 3.5 | 0.050 |
| T4 | H5 | X5 | H4 | 0.0 | 14.0 | 0.7 | 0.003 |
| T4 | H61 | X5 | H4 | 0.0 | 14.0 | 0.4 | 0.001 |
| T4 | H62 | X5 | H4 | 0.0 | 15.8 | 0.4 | 0.001 |
| T4 | HA3 | X5 | H4 | 0.0 | 15.0 | 3.2 | 0.050 |
| G3 | H1 | X5 | H5 | 0.0 | 14.9 | 9.6 | 0.418 |
| G3 | H2 | X5 | H5 | 0.0 | 12.3 | 2.9 | 0.058 |
| G3 | H3 | X5 | H5 | 0.0 | 17.0 | 3.5 | 0.044 |
| G3 | H4 | X5 | H5 | 0.0 | 17.0 | 0.6 | 0.001 |
| G3 | H5 | X5 | H5 | 0.0 | 17.0 | 2.7 | 0.026 |
| T4 | H5 | X5 | H5 | 0.0 | 12.9 | 3.0 | 0.054 |
| T4 | H61 | X5 | H5 | 0.0 | 12.9 | 1.7 | 0.018 |
| T4 | H62 | X5 | H5 | 0.0 | 17.0 | 1.5 | 0.009 |
| T4 | HA3 | X5 | H5 | 0.0 | 13.3 | 5.6 | 0.192 |

[a] The donor and acceptor nuclei are those on the indirect and direct dimensions of the spectrum, respectively. [b] Refer to Figure S5 for nomenclature. [c] Measured height, calculation error applied, predicted heights, and calculated mean $\chi^2$, as described in (5)). Data measured from a 2D-NOESY recorded with 700ms mixing time on a 3 mM sample at pH 6.0 in 100% (v/v) $D_2O$ and at 5 °C.

**Table S10. Summary of $\chi^2$ fits for all structural restraints for residues G3-X5 in $HA_4$-2AA.**

| Restraint | Number of restraints | Total $\chi^2$ | $\chi^2$/ restraint |
|---|---|---|---|
| $^3J_{HH}$ | 9 | 2.8 | 0.31 |
| NOEs | 41 | 29.6 | 0.72 |
| noNOEs | 94 | 18.4 | 0.20 |
| Total | 144 | 50.8 | 0.35 |

**Table S11. Torsion angle conformational preferences for $HA_4$-2AA in aqueous solution.**

| Torsion | Torsion definition [a] | Number of modes [b] | Mean angle (μ) of mode (°) | Libration angle (σ) of mode (°) [c] | Occupancy (π) [d] |
|---|---|---|---|---|---|
| **Based on identity of chemical shifts and coupling constants, the solution dynamic 3D structure parameters for the unmodified HA oligosaccharide portion (residues G1-G3) were taken from) (5):** | | | | | |
| G1 C5 – C6 | G1 C4 – G1 C5 – G1 C6 – G1 O61 | 1 | 98 | 15 | 1.00 |
| G1 – S2 Φ | G1 C2 – G1 C1 – S2 O3 – S2 C3 | 1 | 171 | 23 | 1.00 |
| G1 – S2 Ψ | G1 C1 – S2 O3 – S2 C3 – S2 C2 | 1 | -122 | 23 | 1.00 |
| S2 C2 – NA1 | S2 C1 – S2 C2 – S2 NA1 – SA CA2 | 1 | 120 | 24 | 1.00 |
| S2 C5 – C6 | S2 C4 – S2 C5 – S2 C6 – S2 O6 | 2 | 180 | 15 | 0.53 |
| | | | 60 | 15 | 0.47 |
| S2 – G3 Φ | S2 C2 – S2 C1 – G3 O4 – G3 C4 | 1 | 177 | 19 | 1.00 |
| S2 – G3 Ψ | S2 C1 – G3 O4 – G3 C4 – G3 C5 | 1 | 145 | 19 | 1.00 |
| G3 C5 – C6 | G3 C4 – G3 C5 – G3 C6 – G3 O61 | 1 | 98 | 15 | 1.00 |
| **Solution dynamic 3D structure parameters for the modified HA oligosaccharide portion (residues G3-X5) determined here:** | | | | | |
| Conformationally independent torsions: | | | | | |
| G3 – T4 Φ | G3 C2 – G3 C1 – T4 O3 – T4 C3 | 1 | 159 | 13 | 1.00 |
| G3 – T4 Ψ | G3 C1 – T4 O3 – T4 C3 – T4 C2 | 1 | -123 | 3 | 1.00 |
| T4 C2 – NA1 | T4 C1 – T4 C2 – T4 NA1 – T4 CA2 | 1 | 114 | 22 | 1.00 |
| T4 C3 – C4 | T4 C2 – T4 C3 –T4 C4 – T4 C5 | 1 | -60 | 19 | 1.00 |
| T4 C4 – C5 | T4 C3 – T4 C4 – T4 C5 – T4 C6 | 1 | 180 | 8 | 1.00 |
| T4 C5 – C6 | T4 C4 – T4 C5 – T4 C6 – T4 O6 | 3 | 60 | 19 | 0.29 |
| | | | -60 | 19 | 0.13 |
| | | | 180 | 19 | 0.58 |
| X5 HN11 – X5 C1 | T4 C1 – X5 N11 – X5 C1 – X5 C6 | 1 | 180 | 2 | 1.00 |
| X5 C6 –C7 | X5 C1 – X5 C6 – X5 C61 – X5 O611 | 1 | 0 | 2 | 1.00 |
| Conformationally co-dependent torsions: [e] | | | | | |
| T4 C2 – C3 | T4 NA1 – T4 C2 – T4 C1 – T4 O3 | 3 | 60 | 20 | 0.35 |
| | | | -60 | 20 | 0.08 |
| | | | -180 | 20 | 0.57 |
| T4 C2 – C1 | T4 NA1 – T4 C2 – T4 C1 – X5 HN11 | 2 | -60 | 5 | 0.92 |
| | | | -180 | 5 | 0.08 |
| T4 C1 – X5 HN11 | T4 C2 – T4 C1 – X5 HN11 – X5 C1 | 5 | -127 | 4 | 0.35 |
| | | | 117 | 4 | 0.35 |
| | | | -96 | 4 | 0.22 |
| | | | -147 | 4 | 0.06 |
| | | | 85 | 4 | 0.02 |
| **Dynamic model parameters not measurable by NMR that were assigned modeled values:** [f] | | | | | |
| T4 Me | T4 NA1 – T4 CA2 – T4 CA3 – T4 HA31 | 3 | 60 | 20 | 0.33 |
| | | | -60 | 20 | 0.33 |
| | | | 180 | 20 | 0.33 |
| T4 HO4 | T4 C3 – T4 C4 – T4 O4 – T4 HO4 | 3 | 60 | 20 | 0.33 |
| | | | -60 | 20 | 0.33 |
| | | | 180 | 20 | 0.33 |
| T4 HO5 | T4 C6 – T4 C5 – T4 O5 – T4 HO5 | 1 | 180 | 20 | 1.00 |
| T4 HO6 | T4 C5 – T4 C6 – T4 O6 – T4 HO6 | 3 | 60 | 20 | 0.33 |
| | | | -60 | 20 | 0.33 |
| | | | 180 | 20 | 0.33 |

[a] Refer to Figure S5 for atom nomenclature. [b] Modes (macrostates) are classified as distinct if rotation of the bond between the two conformers passes through a van der Waals maximum. [c] The libration amplitude is the standard deviation of the distribution of dihedral angle values from the mean value that the torsion typically adopts for a given mode. [d] The occupancy of each mode is expressed as a proportion of the total for that torsion. [e] Refer to Table 3 for the co-dependency relationships between these 5 modes. [f] Torsions involving hydroxyls and carboxylate groups were given values based on the dynamic 3D structure used for $HA_6^{AN}$ (6), expectations from mining the Cambridge Structural Database (7) and consideration of their positions within each determined conformer.

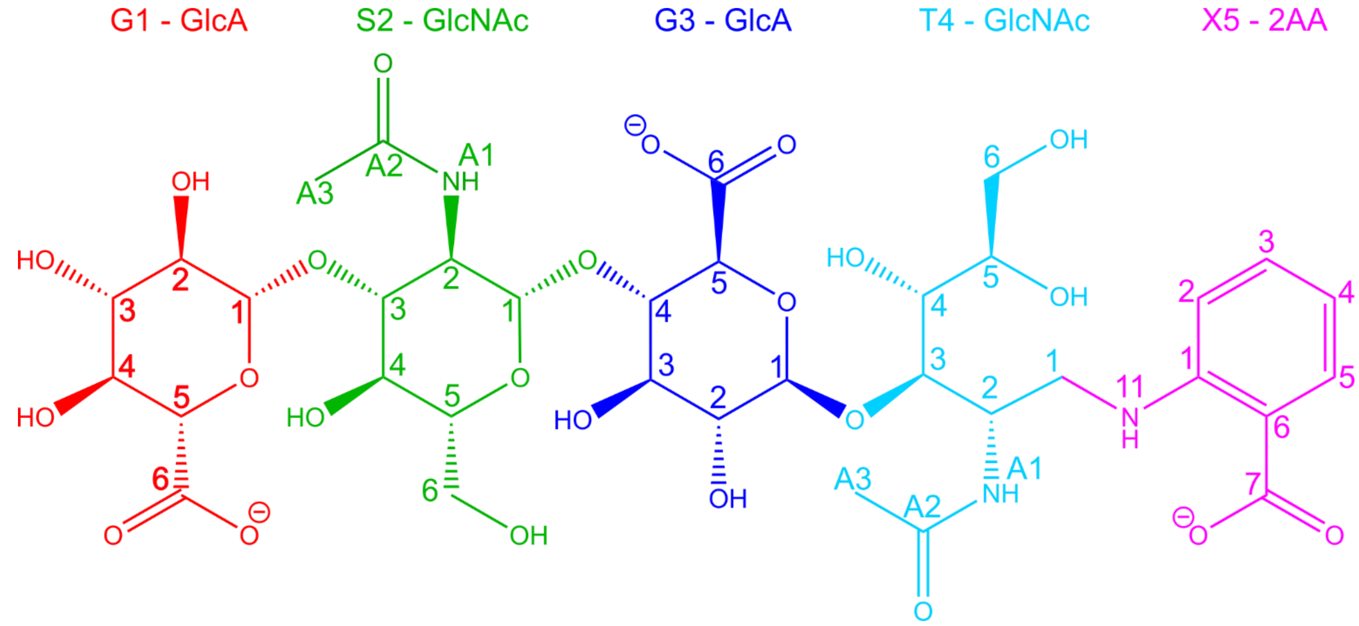


**Figure S5. 2D structure and nomenclature of HA$_4$-2AA.** The molecule is divided into 5 residues (corresponding to the monosaccharide rings and the 2AA group) and the atom labels within each residue are shown.

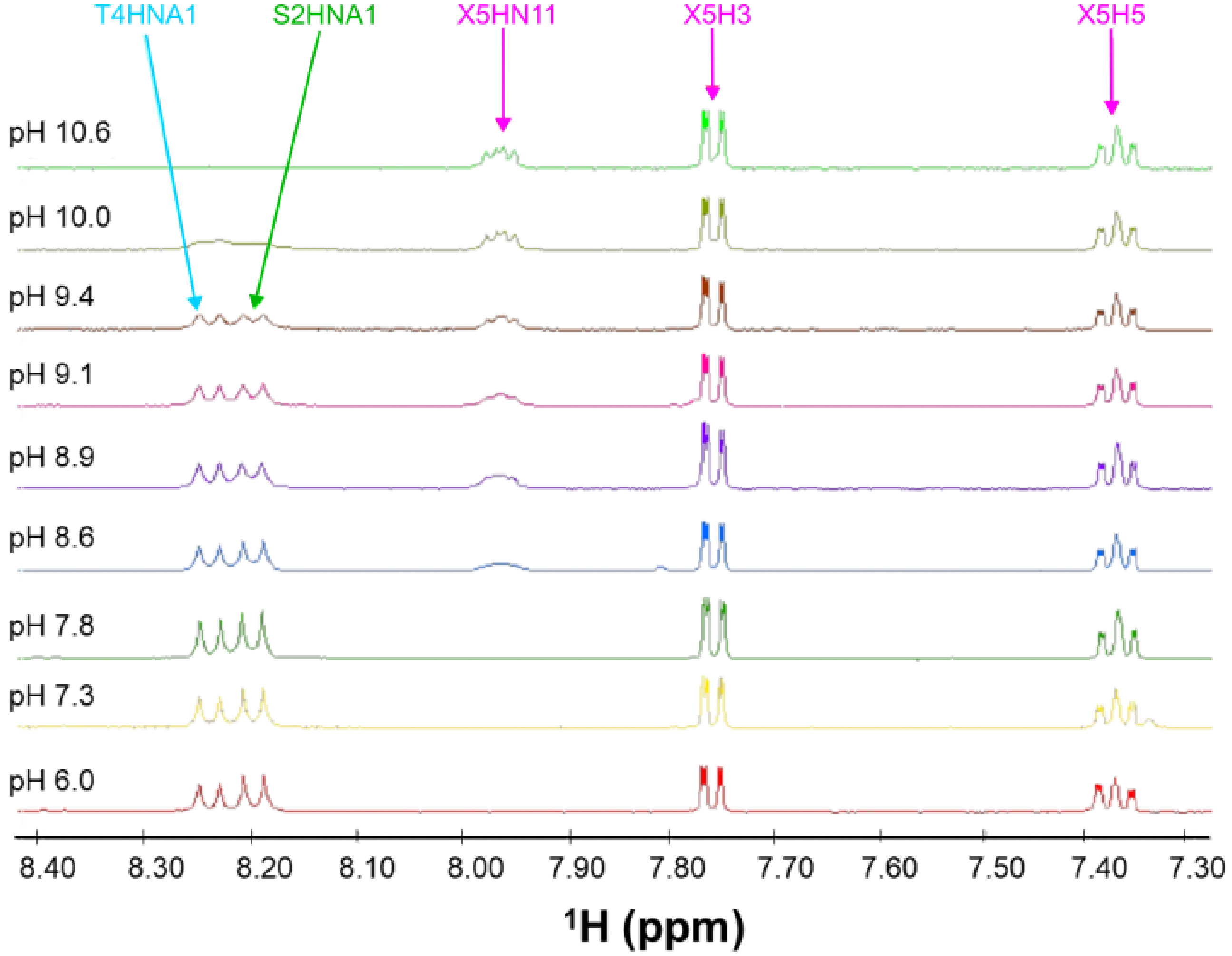


**Figure S6. pH titration of HA$_4$-2AA.** Portions of 1D $^1$H spectra recorded at 5 °C on a sample of HA$_4$-2AA (1 mM, 10% (v/v) $D_2O$) at different pH values (shown from pH 6.0 - 10.6). The difference in exchange rate of the amide (S2 HNA1, T4 HNA1) and aniline (X5 HN11) hydrogens with pH is clearly visible in the differential line-broadening as the pH value is changed. Aromatic hydrogens (X5 H3, X5 H5) do not exchange with solvent and are equally sharp at all pH values.

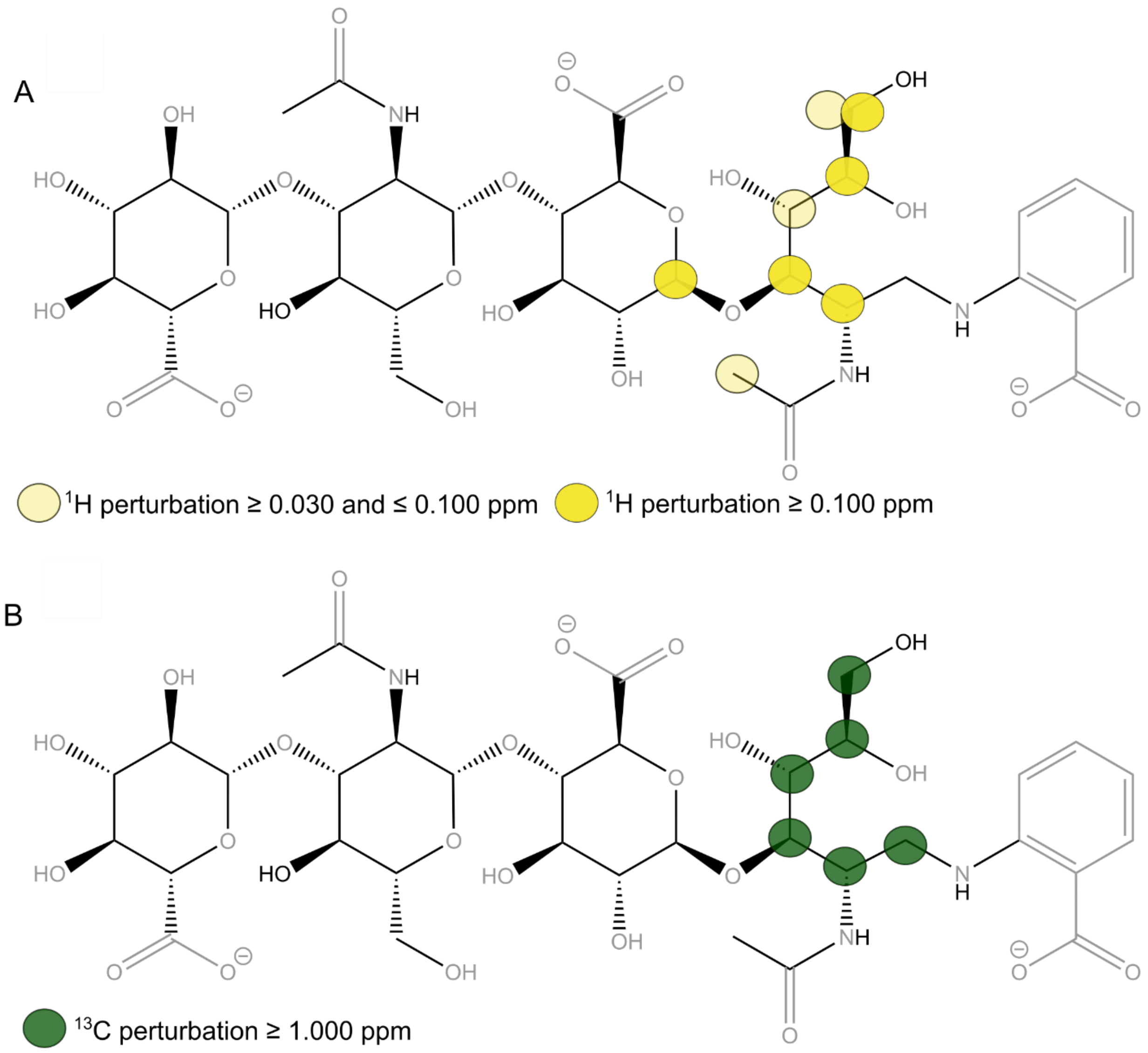


**Figure S7. Chemical shift perturbations in $HA_4$-2AA compared to $HA_4^{AN}$.** The 2D structure of $HA_4$-2AA is shown with colored circles indicating where in the structure $^{1}$H (**A**) and $^{13}$C (**B**) chemical shifts are perturbed compared to $HA_4^{AN}$. Comparison is between $HA_4^{AN}$ at 5 – 20 mM, pH 6.0, 24.4 °C, 10% (v/v) $D_2O$ (values from (3)) and $HA_4$-2AA at 1 mM, pH 9.4, 5.0 °C, 10% (v/v) $D_2O$. Atoms in grey are those for which chemical shifts were not measurable (O and H in OH), not measured (N atoms or C in carboxylate groups), or those for which comparison of chemical shift assignment (to $HA_4^{AN}$) is not possible (*e.g.,* residue X5-2AA).

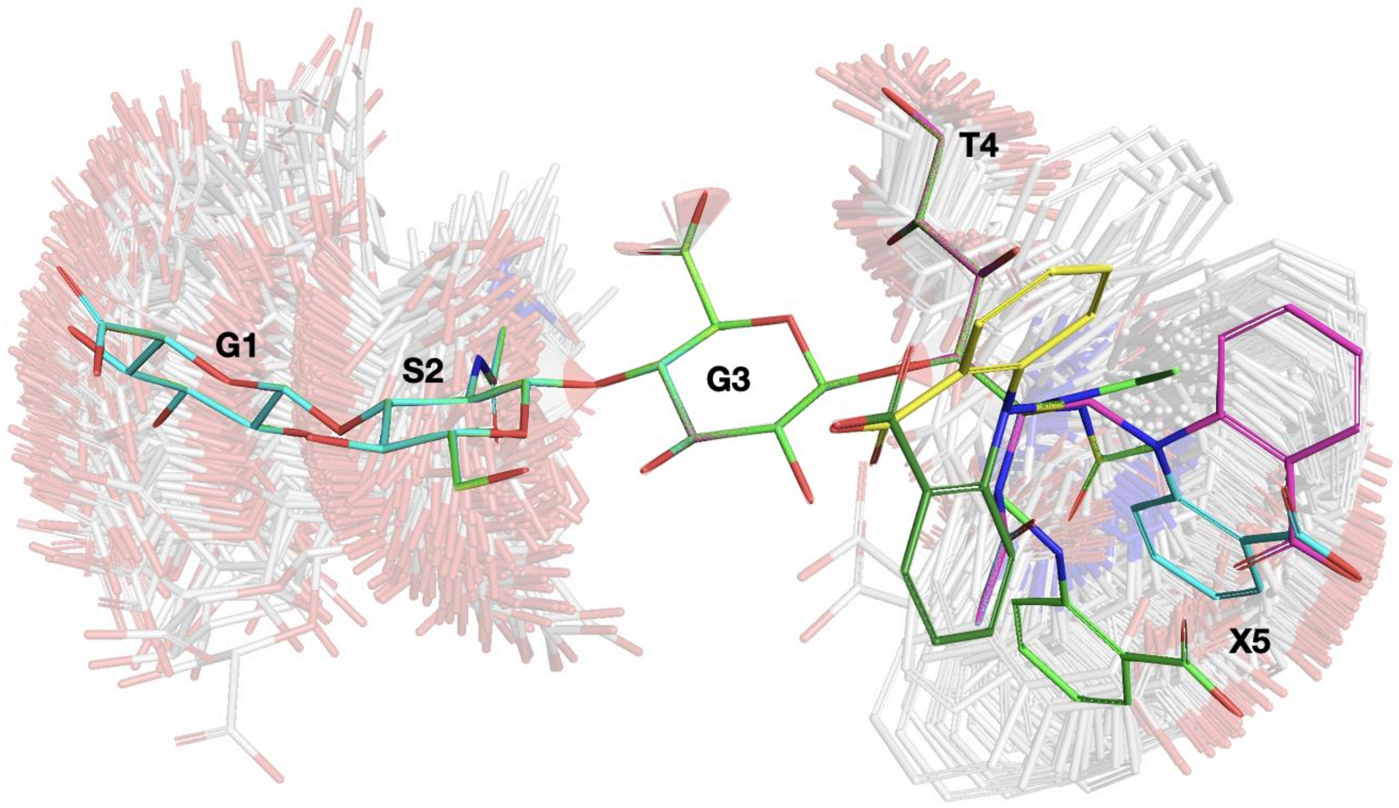


**Figure S8. Solution dynamic 3D structure of $HA_4$-2AA in a conformational ensemble representation**. The 5 Conformers generated by the co-dependent behavior of T4 C2 – C3, T4 C2 – C1 and T4 C1 – X5 HN11 (bright conformations) are overlaid together on an ensemble of conformations representing the Gaussian libration about those mean positions (faded conformations). Carbon atoms are colored differently for each Conformer: 1 (green), 2 (cyan), 3 (magenta), 4 (yellow), and 5 (dark green). All Conformers and conformations are overlaid onto the ring atoms of residue G3 (GlcA). Oxygen atoms are shown in red and nitrogen atoms in blue; hydrogen atoms are omitted for clarity. See Figure S5 for residue nomenclature.

*E. Generation of a model of the $HA_6$-2AA/Link_TSG6 complex*

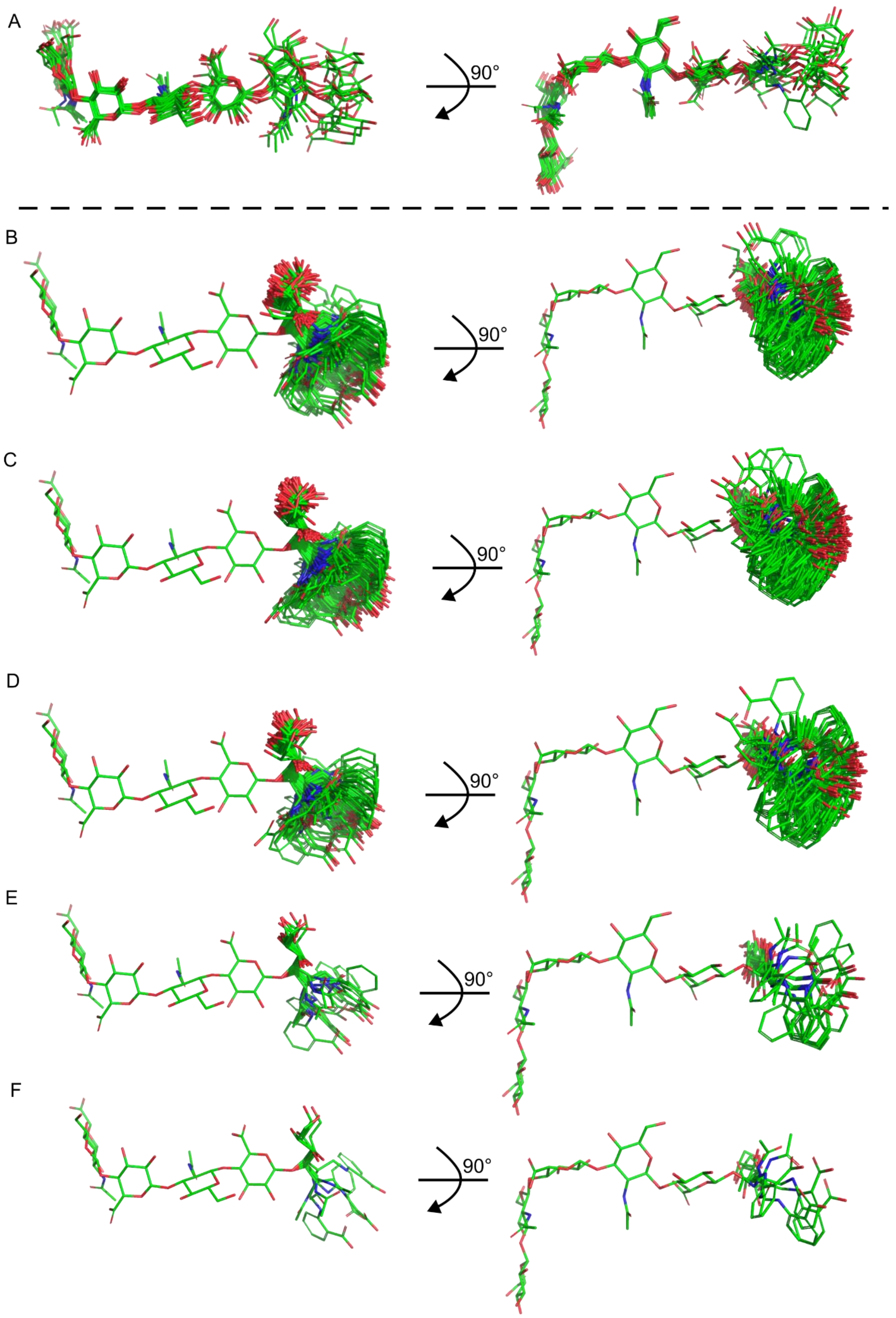


**Figure S9. Comparison of conformations of $HA_8^{AN}$ bound to Link_TSG6 with the modeled conformations of $HA_6$-2AA used for docking into the binding site of Link_TSG6. A)** An overlay of the 10 conformations of residues 1-7 of $HA_8^{AN}$ when bound to Link_TSG6 as determined from experiment-based modeling (8). **B-F)**. 250 modeled conformations for the chemically modified residues of $HA_6$-2AA (6 sugar residues and 2AA) attached to the conserved residues from the lowest energy bound conformation of $HA_8^{AN}$ (residues 1-4); the distribution and number of conformations in B-F represents the behavior and occupancy of each Conformer 1-5 in solution.

*F. Supporting Information References*

6. Blundell, C. D., and Almond, A. (2010) Method for determining three-dimensional structures of dynamic molecules. US Patent number US10482997B2

7. Groom, C. R., Bruno, I. J., Lightfoot, M. P., and Ward, S. C. (2016) The Cambridge structural database. *Acta Crystallogr B Struct Sci Cryst Eng Mater*. **72**, 171–179

8. Higman, V. A., Briggs, D. C., Mahoney, D. J., Blundell, C. D., Sattelle, B. M., Dyer, D. P., Green, D. E., DeAngelis, P. L., Almond, A., Milner, C. M., and Day, A. J. (2014) A Refined Model for the TSG-6 Link Module in Complex with Hyaluronan. *Journal of Biological Chemistry*. **289**, 5619–5634
